# A Tyrosine Phospho-switch within the Longin Domain of VAMP721 modulates SNARE functionality

**DOI:** 10.1101/2023.03.18.533296

**Authors:** Martiniano Maria Ricardi, Niklas Wallmeroth, Cecilia Cermesoni, Dietmar Gerald Mehlhorn, Sandra Richter, Lei Zhang, Josephine Mittendorf, Ingeborg Godehardt, Kenneth Wayne Berendzen, Edda von Roepenack-Lahaye, York-Dieter Stierhof, Volker Lipka, Gerd Jürgens, Christopher Grefen

## Abstract

The final step in secretion is membrane fusion facilitated by SNARE proteins that reside in opposite membranes. The formation of a trans-SNARE complex between one R and three Q coiled-coiled SNARE domains drives the final approach of the membranes providing the mechanical energy for fusion. Biological control of this mechanism is exerted by additional domains within some SNAREs. For example, the N-terminal Longin domain (LD) of R-SNAREs (also called Vesicle-associated membrane proteins, VAMPs) can fold back onto the SNARE domain blocking interaction with other cognate SNAREs. The LD may also determine the subcellular localization via interaction with other trafficking related proteins. Here, we provide cell-biological and genetic evidence that phosphorylation of the Tyrosine57 residue regulates the functionality of VAMP721. We found that an aspartate mutation mimics phosphorylation, leading to protein instability and subsequent degradation in lytic vacuoles. The mutant SNARE also fails to rescue the defects of *vamp721vamp722* loss-of-function lines in spite of its wildtype-like localization within the secretory pathway and the ability to interact with cognate SNARE partners. Most importantly, it imposes a dominant negative phenotype interfering with root growth, normal secretion and cytokinesis in wildtype plants generating large aggregates that mainly contain secretory vesicles. Non-phosphorylatable VAMP721^Y57F^ needs higher gene dosage to rescue double mutants in comparison to native VAMP721 underpinning that phosphorylation modulates SNARE function. We propose a model where a short-lived phosphorylation of Y57 serves as a regulatory step to control VAMP721 activity, favouring its open state and interaction with cognate partners to ultimately drive membrane fusion.

## Introduction

Eukaryotic cells have evolved an intricate trafficking system to enable transport of membrane material, proteins, metabolites, and/or other molecules between subcellular compartments and into the apoplast (More et al., 2020). Transport vesicles shuttling between these compartments are the key component of a functional secretory system (Cai et al., 2007); their genesis and maintenance requires a phalanx of different protein families that regulate budding at the donor membrane, transport to, tethering and docking at the target membrane (Kanazawa and Ueda, 2017). The final step – fusion of both membranes – is facilitated by a family of SOLUBLE N-ETHYLMALEIMIDE–SENSITIVE FACTOR ATTACHMENT PROTEIN RECEPTOR PROTEINS (SNAREs). SNAREs’ core function in enabling vesicle fusion places these proteins vital for a number of cellular and physiological processes in plants that rely on secretory traffic, such as cytokinesis (Jurgens, 2005; El Kasmi et al., 2013), pathogen defence (Lipka et al., 2007; Kwon et al., 2008) and polar growth (Enami et al., 2009; Ichikawa et al., 2014; Slane et al., 2017).

Most SNAREs are tail-anchored membrane proteins with a C-terminal transmembrane domain and are inserted into the ER membrane post-translationally via the so-called Guided-Entry of Tail-anchored proteins (GET) pathway (Stefanovic and Hegde, 2007; Schuldiner et al., 2008; Xing et al., 2017). Their eponymous ‘SNARE-motif’ – an α-helix coiled-coil domain – faces the cytosol and is responsible for the tight interaction with partner SNAREs ultimately driving membrane fusion. Depending on a conserved polar residue within this domain they are divided into Q- and R-SNAREs (Fasshauer et al., 1998). Membrane fusion requires the assembly of a trans-SNARE complex containing one R-SNARE and three Q-SNARE domains (Qa,b,c). Additionally, most SNAREs feature an N-terminal domain that can fold back on the SNARE-motif to regulate its interaction with cognate SNARE partners (Jahn and Scheller, 2006).

The VESICLE-ASSOCIATED MEMBRANE PROTEIN 7 (VAMP7) R-SNARE family in *Arabidopsis thaliana* is composed of two subsets, four *VAMP71* and eight *VAMP72* genes. All of them share sequence homology with the unique human VAMP7 and resulted from gene duplication and divergence events (Liu et al., 2014). Most of them share a common domain structure composed of an N-terminal Longin domain (LD) followed by the R-SNARE (RD) and C-terminal transmembrane domain (=’tail-anchor’). *VAMP721* and its closest homologue *VAMP722* are strongly and ubiquitously expressed in all developmental stages and tissues of *Arabidopsis thaliana* (Uemura et al., 2004; Lipka et al., 2007). Both proteins are mainly vesicle-localized and form different complexes with their cognate Q-SNARE partners, the Qa-SNAREs SYP121/PEN1 in general secretion (Kwon et al., 2008) or SYP111/KNOLLE (Qa), NPSN11 (Qb), SYP71 (Qc), and SNAP33 (Qbc) during cytokinesis (Zhang et al., 2011; El Kasmi et al., 2013).

While loss of one *VAMP* does not lead to an apparent phenotype in *Arabidopsis*, the *vamp721^−/−^vamp722^−/−^* double mutants are seedling lethal and show severe cytokinesis defects in seedlings (Zhang et al., 2011; El Kasmi et al., 2013). The ‘homo-heterozygous’ mutants *vamp721^−/−^vamp722^−/+^* and *vamp721^−/+^vamp722^−/−^* show wildtype appearance under normal growth conditions, but are haplo-insufficient in responses to biotic and abiotic stresses (Kwon et al., 2008; Yi et al., 2013).

Structural analyses of the mammalian VAMP7 revealed that the RD determines the interaction with Q-SNARE partners while the LD has regulatory effects determining subcellular localization through additional interactions with other proteins (Martinez-Arca et al., 2003). VAMP7 has two main configurations: a ‘closed’ state - fusogenically inactive conformation - with the LD folded back onto the RD, or ‘open’, with the RD available for interaction with Q-SNARE partners. In vitro assays showed that VAMP7 autoinhibition is not too strong since the protein can spontaneously form SNARE complexes *in-vitro.* This capacity is increased when the protein is deprived of its LD, underpinning the importance of the latter for spatio-temporal regulation of vesicle fusion (Martinez-Arca et al., 2003). Interaction with the endosomal VPS9 AND ANKYRIN REPEAT– CONTAINING PROTEIN (Varp) can strengthen the autoinhibitory mechanisms and trap the protein in a closed conformation (Schafer et al., 2012). Additionally, VAMP7 can also bind to AP-3 (for trafficking to synaptic vesicles, endosomes and lysosomes) or Hrb (for recycling) when exposing its LD. Varp, AP-3 and Hrb all bind to the same motif within the LD and it is through this sequential binding that VAMP7 can reach it’s different target compartments (Daste et al., 2015).

In recent years, SNAREs have also been implicated in controlling the activity of plasma membrane K^+^-channels through physical interaction of exclusive domains outside of the SNARE motif (Honsbein et al., 2009; Grefen et al., 2010; Grefen et al., 2015; Zhang et al., 2015). Interestingly, the cognate SNARE partners SYP121 and VAMP721 seem to antagonistically control the potassium channel KAT1, thereby regulating potassium influx in coordination with vesicle trafficking (Grefen et al., 2011; Honsbein et al., 2011; Zhang et al., 2015). In the course of these works, the mutation of a tyrosine residue within the LD of VAMP721 was discovered to prevent interaction with and control of the potassium channel by the mutated SNARE (Zhang et al., 2015). FRET experiments with a Y57D mutation in the Arabidopsis VAMP721 suggested that such mutation causes the R-SNARE to adopt an ‘open’ conformation (Zhang et al., 2017). The mutation also interferes with K^+^-channel interaction reducing potassium influx at the plasma membrane, and surprisingly, mutation of an Aspartate residue within the non-interacting, close VAMP721 homolog VAMP723 into VAMP723^D57Y^ rendered the latter interaction-competent with the K^+^-channel (Zhang et al., 2015; Zhang et al., 2017).

Here, we report our findings regarding the implications of the tyrosine motif for the VAMPs core function in vesicle trafficking. We were particularly interested to analyse, whether the VAMP721^Y57D^ mutant would still complement the phenotype of the seedling-lethal *vamp721^−/−^vamp722^−/−^* double mutants and if the Tyrosine 57 is indeed a phosphorylation site regulating VAMP721 function, localization and interactions.

## Results

### Mutation of Tyrosine 57 to Aspartate impedes rescue of the seedling lethal phenotype

The suggestion that the Y57D mutation leads to conformational changes sparked our interest with a view to its potential to rescue the cytokinesis defect of the *vamp721^−/−^vamp722^−/−^ loss of function* line (Zhang et al., 2011; El Kasmi et al., 2013) and its dependence on other SNARE related features such as interaction with cognate partners and/or subcellular localisation. We designed full-length genomic constructs of VAMP721, VAMP721^Y57D^ and VAMP723 (as negative control) and included an N-terminal fusion of mEGFP-myc for visualization and detection. To achieve native expression levels, we designated the 1333 bp fragment upstream of the VAMP721 transcription start as promoter region (‘*P_VAMP721_*’) for all constructs (Figure 1A). Due to the lethality of the *vamp721^−/−^vamp722^−/−^* genotype (Zhang et al., 2011) all transformations were performed in the *vamp721^−/−^vamp722^−/+^* genetic background.

**Figure 1:**
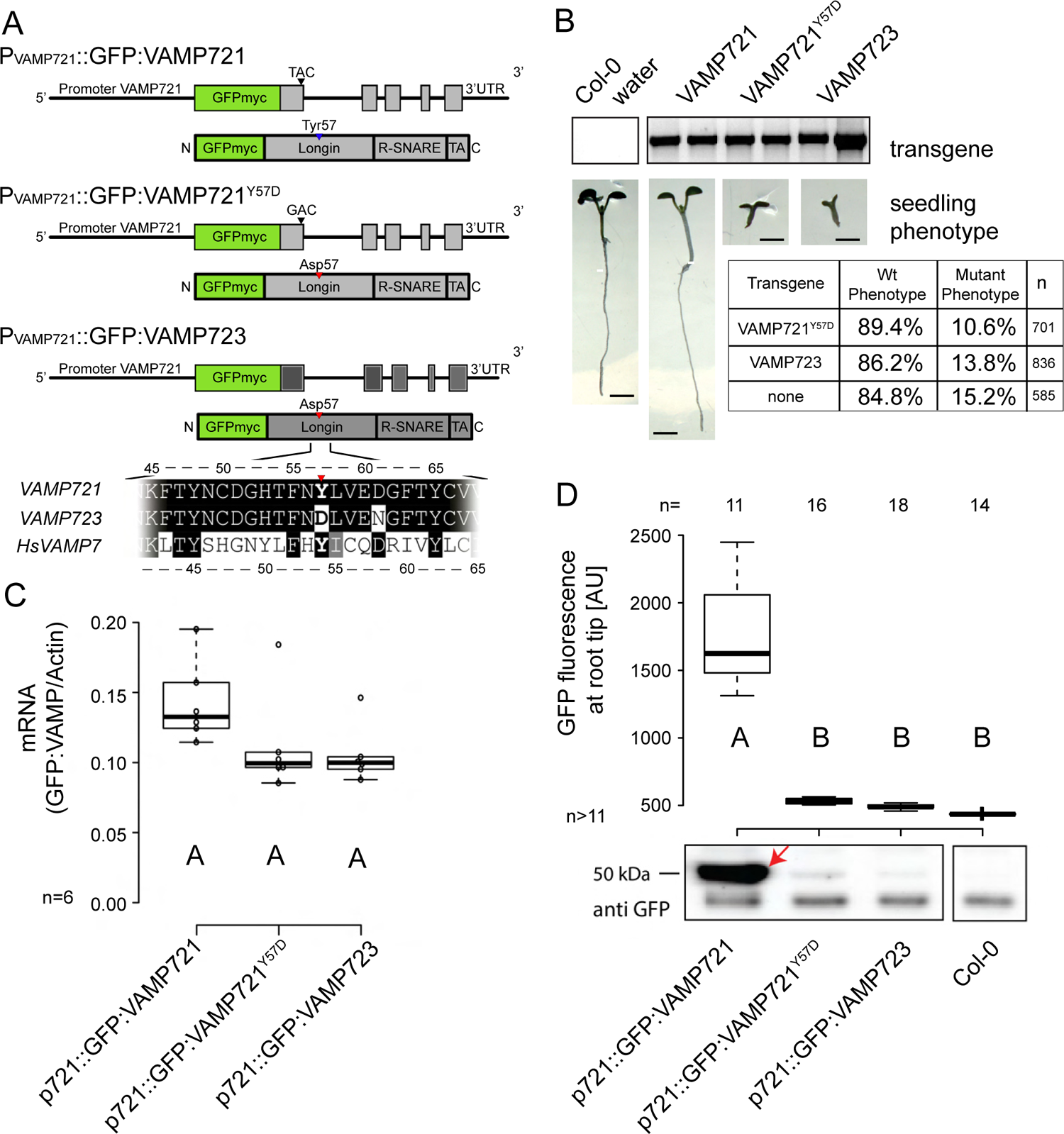
Expression of GFP-tagged VAMP constructs driven by the native VAMP721 promoter in the vamp721 vamp722 T-DNA insertion line. (A) Cartoon of genomic constructs above their resulting protein. Residue 57 is depicted with red or blue triangle, respectively. (B) Detection of the corresponding complementation transgene in vamp721−/−vamp722−/− background and its resulting phenotype underneath. Inset table lists the ratio of wildtype vs. mutant phenotype recorded in vamp721−/−vamp722-/+ descendants that contain the complementation constructs VAMP721Y57D and VAMP723, respectively. Assuming heterozygosity of the transgene we expect a yield of 3.1% offspring with mutant phenotype in case of full rescue. (C) Boxplot depicting qRT-PCR analysis of transcript abundancy in transgenic lines using primers specific for the complementation constructs. Values are normalized to the mRNA levels of Actin. (D) Boxplot depicting quantification of GFP fluorescence at the root tip of transformants (see Supplemental Figure 1B) and immunoblot underneath (anti-cYFP) of total seedling extracts. GFP-VAMP fusions are expected at a size of around 50 kDa (red arrow). Letters A and B indicate statistically different groups. Boxplot design: Center lines of boxes represent the median, with outer limits at the 25th and 75th percentiles. Tukey whiskers extend to 1.5 times the interquartile range (IQR). Outliers are depicted as hollow dots. Number of biological replicates is labelled by the letter ‘n’ throughout this figure. Scale bar = 2mm.

We screened more than 700 transformants but only observed rescue of double homozygous mutant lines when the *P_VAMP721_*::VAMP721 transgene was used for complementation (Figure 1B and Supplemental Figure 1). Conversely, *vamp721^−/−^vamp722^−/−^* seedlings carrying the respective *P_VAMP721_*::VAMP721^Y57D^ or *P_VAMP721_*::VAMP723 transgenes died at early seedling stage. Segregation analysis of T2 seedlings in the transformed *vamp721^−/−^vamp722^−/+^* progeny confirmed the lack of rescue (Figure 1B), and showed a lower segregation of the VAMP722 mutant allele (10.6-15.2% vs. 25% expected), a transmission defect previously reported (Kwon et al., 2008).

While mRNA levels in the transgenic lines (assessed by qRT-PCR) are not significantly different (Figure 1C), there are noticeable differences on protein level. The GFP-VAMP721 protein was readily detected by immunoblot analysis, however, both GFP-VAMP721^Y57D^, and GFP-VAMP723 proteins were below detection levels (Figure 1D). Using epi-fluorescence we corroborated the differences in protein levels as several independent VAMP721^Y57D^ and VAMP723 transgenic lines showed very low GFP signals (Figure 1D and Supplemental Figure 1B). Our data suggests cellular recognition of mutant SNAREs (Y57D) or aberrant (VAMP723) protein levels and their subsequent degradation.

To further understand this connection, we tested the expression strength of the VAMP723 promoter. *P_VAMP723_::*VAMP721 failed to rescue the double mutant phenotype and showed around 50-fold less mRNA in comparison to *P_VAMP721_* driven transcripts with virtually undetectable protein expression (Supplemental Figure 1C-E). Segregation analyses showed an average 7.5% of seedlings (74 out of 986) with mutant phenotype in three independent transgenic events differing from the expected 2.5%-3.75% for a full rescue in T2 plants and close to the 10-15% expected for lack of rescue.

Taken together, VAMP721 levels are required to be tightly controlled and maintained at high levels to facilitate successful secretory trafficking during cytokinesis and for cellular homeostasis. Also, VAMP723 is unable to complement the seedling lethal phenotype even under strong expression from the native VAMP721 promoter. However, this will also be a consequence of the sequence differences in critical positions within the RD of VAMP723 compared to VAMP721/722 (Sanderfoot, 2007; Liu et al., 2014).

### VAMP721^Y57D^ protein is unstable and traffics to the vacuole via the pre-vacuolar compartment (PVC) for degradation

Since the Aspartate containing constructs (VAMP721^Y57D^ and VAMP723) driven by the endogenous promoter *P_VAMP721_* were virtually impossible to detect (Figure 1D and Supplemental Figure 1B), we constructed Estradiol-inducible (EDI) lines (Figure 2A) (Zuo et al., 2000). The EDI lines were generated in a Col-0 background and showed detectable GFP-VAMP721 protein levels upon induction. After selecting the three strongest expression lines for each construct, we noticed that GFP-VAMP721 still yielded higher protein levels compared to the mutant version and VAMP723, respectively (Figure 2B and Supplemental Figure 1F), reminiscent of our previous observations using the endogenous promoter (Figure 1D).

**Figure 2:**
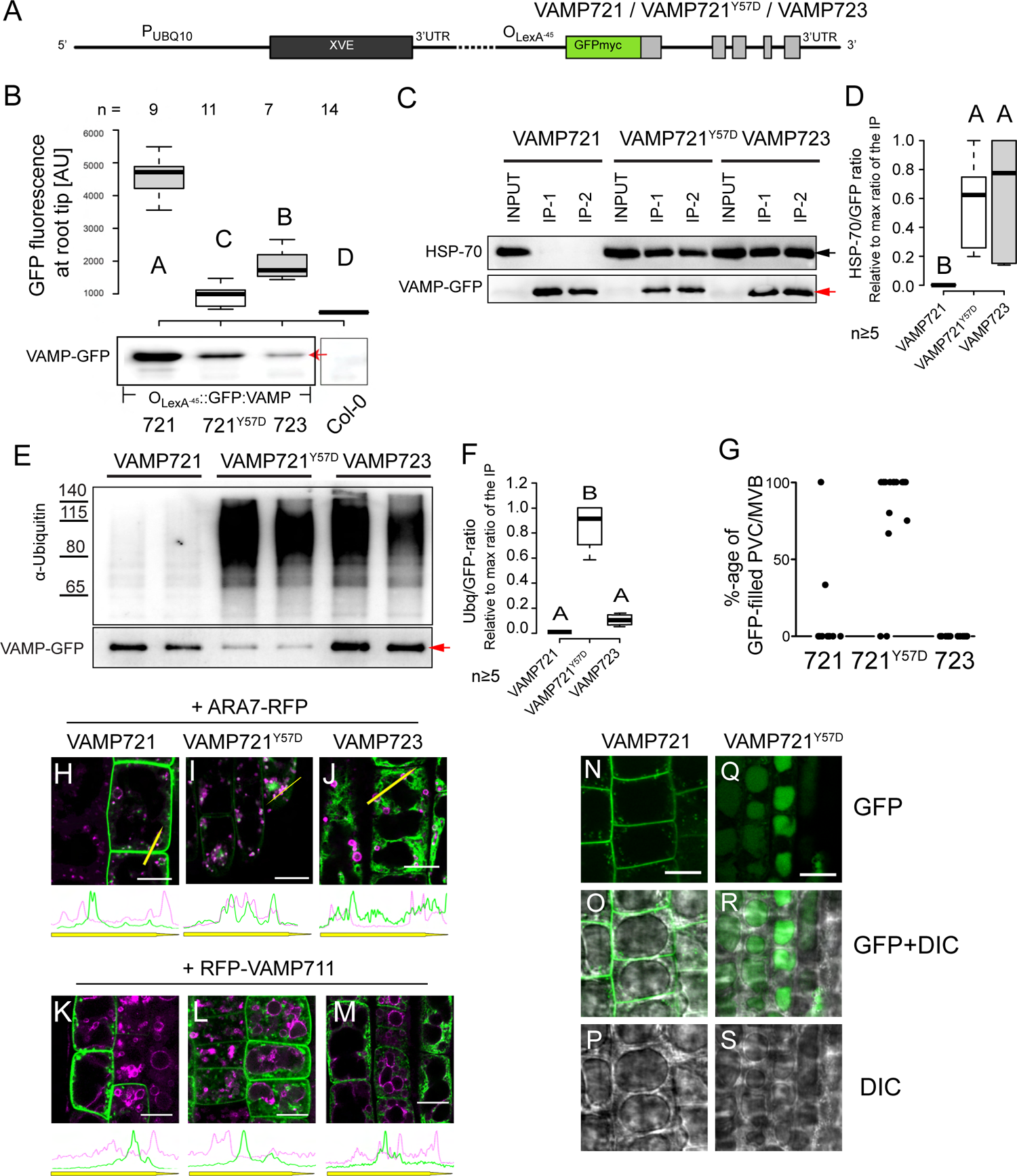
Inducible expression of GFP-VAMP fusion constructs. (A) Cartoon of Estradiol-inducible constructs (EDI) comprising a tandem repeat of LexA operators fused to the minimal, −45bp 35S promoter, here termed: OLexA-45 transformed into Col-0 plants. (B) Boxplot depicting quantification of GFP fluorescence at the root tip of transformants after Estradiol induction (see Supplemental Figure 2) and immunoblot underneath (anti-cYFP) of total seedling extracts. GFP-VAMP fusions are expected at a size of around 50 kDa (red arrow). Letters A and B indicate statistically different groups. Boxplot design: Center lines of boxes represent the median, with outer limits at the 25th and 75th percentiles. Tukey whiskers extend to 1.5 times the interquartile range (IQR). Outliers are depicted as hollow dots. Number of biological replicates is labelled by the letter ‘n’ throughout this figure. (C, E) Co-IP assays of GFP-VAMP lines after 6 hours of 10µM Estradiol induction. GFP-VAMP protein was immunoprecipitated with anti-GFP beads and protein blots were probed with antisera indicated to the left. (D, F) Signal intensities of bands from at least 5 repetitions were measured with ImageJ. Ratio of GFP-VAMP (red arrowhead indicating the expected 50kDa band) was also detected and used to calculate relative IP values in the blots where D) HSP70 (black arrowhead indicating the expected 70 kDa band) or F) Polyubiquitin were detected (n≥5). The Signal to VAMP ratios were then relativized to the max ratio observed for each row of IP assays to compensate for differences between rows. A and B indicate significantly different ratios. (G) Percentage of GFP-filled PVC/MVB vesicles and representative confocal images of crosses between ARA7-RFP with (H) VAMP721, (I) VAMP721Y57D, and (J) VAMP723 lines. Representative images of crosses between VAMP711 (staining the vacuole tonoplast) and (K) VAMP721, (L) VAMP721Y57D, and (M) VAMP723 lines. Line histograms underneath images (H-M) show fluorescence intensity along yellow arrows between VAMP constructs and respective marker line. Dark treated seedlings expressing (N-P) GFP-VAMP721 or (Q-S) GFP-VAMP721Y57D visualize presence of the latter in the vacuole. Scale bar = 10µm.

To explore the possibility of proteolytic degradation, we treated six days old EDI seedlings of VAMP721, VAMP721^Y57D^ and VAMP723 with Estradiol prior to performing co-Immunoprecipitation (co-IP) assays to check for potential interaction with the cytosolic isoforms of the HEATSHOCK PROTEIN 70 (HSP70) chaperone, a master regulator of protein degradation (Fernandez-Fernandez et al., 2017). Indeed, both VAMP721^Y57D^ and VAMP723 interact strongly with HSP70 in contrast to VAMP721 that shows no detectable interaction (Figure 2C, D). This interaction correlates with the detection of poly-ubiquitination in VAMP721^Y57D^ and to a lesser extent VAMP723 (Figure 2E, F).

Given that poly-ubiquitination of membrane proteins is a tag for their degradation at the vacuole (Scheuring et al., 2012), we crossed our constructs with marker lines for the PVC (ARA7, Figure 2H-J and Supplemental Figure 2A-C (Geldner et al., 2009)), the vacuole (VAMP711, Figure 2K-M and Supplemental Figure 2D-F (Geldner et al., 2009)), and autophagosomes (NBR1, Supplemental Figure 2G-I (Hafren et al., 2017)). For visualization of the PVC, plants were treated with wortmannin-A which induces the swelling of the PVC compartments labelled by the Rab GTPase Ara7 (Wang et al., 2009). We observed that VAMP721^Y57D^ was present inside the PVC (Figure 2 G, I and Supplemental Figure 2B) more frequently than the wildtype VAMP721 (Figure 2G, H). Using the tonoplast marker VAMP711 to better visualise the vacuole, neither of the three fusion proteins could be detected within or at this compartment when seedlings were grown in light. To visualize GFP inside the vacuole, we performed four hours of dark treatment on pre-induced lines (Tamura et al., 2003) yielding a clear signal inside of the vacuole for VAMP721^Y57D^, while VAMP721 was completely absent (Figure 2N-S). Co-localisation studies with the autophagosome marker NBR1-RFP (Supplemental Figure 2G-I) did not detect any of the VAMPs in that compartment.

### VAMP721^Y57D^ overlaps with normal VAMP721 subcellular localization in addition to its PVC/vacuole localization

Previous reports showed that wildtype VAMP721 localized to the trans-Golgi network (TGN), plasma membrane (PM), cell plate and endosomes, while VAMP723 is retained in the Endoplasmic Reticulum (ER). This ER retention, however, is solely based on transient and/or protoplast transformations (Uemura et al., 2004; Zhang et al., 2011; El Kasmi et al., 2013; Hecker et al., 2015; Zhang et al., 2015; Uemura et al., 2019). Using the stably transformed EDI lines, we performed immunostaining with anti-SYP111 antibodies to first assess VAMP721 localization at the cell plate (Touihri et al., 2011) (Figure 3A-C and Supplemental Figure 3A-C). Both VAMP721 and VAMP721^Y57D^ constructs showed a strong cell plate localization with weak signals in other endosomal compartments of dividing cells. By contrast, and in line with previous observations using transient overexpression (Uemura et al., 2004), VAMP723 localised to the ER (Figure 3C and Supplemental Figure 3C). Since the ER is ubiquitously present in dividing cells and squeezed in close proximity to the emerging cell plate during cytokinesis, it is not possible to exactly discern co-localisation of SYP111 and VAMP723 at the cell plate with these confocal microscopy images. Instead, we co-expressed the constructs with an RFP-tagged ER reporter line (CD3-959) (Nelson et al., 2007) to confirm a predominant ER localization for VAMP723 as opposed to VAMP721 and its Y57D mutant (Figure 3D-G and Supplemental Figure 3D-F).

**Figure 3:**
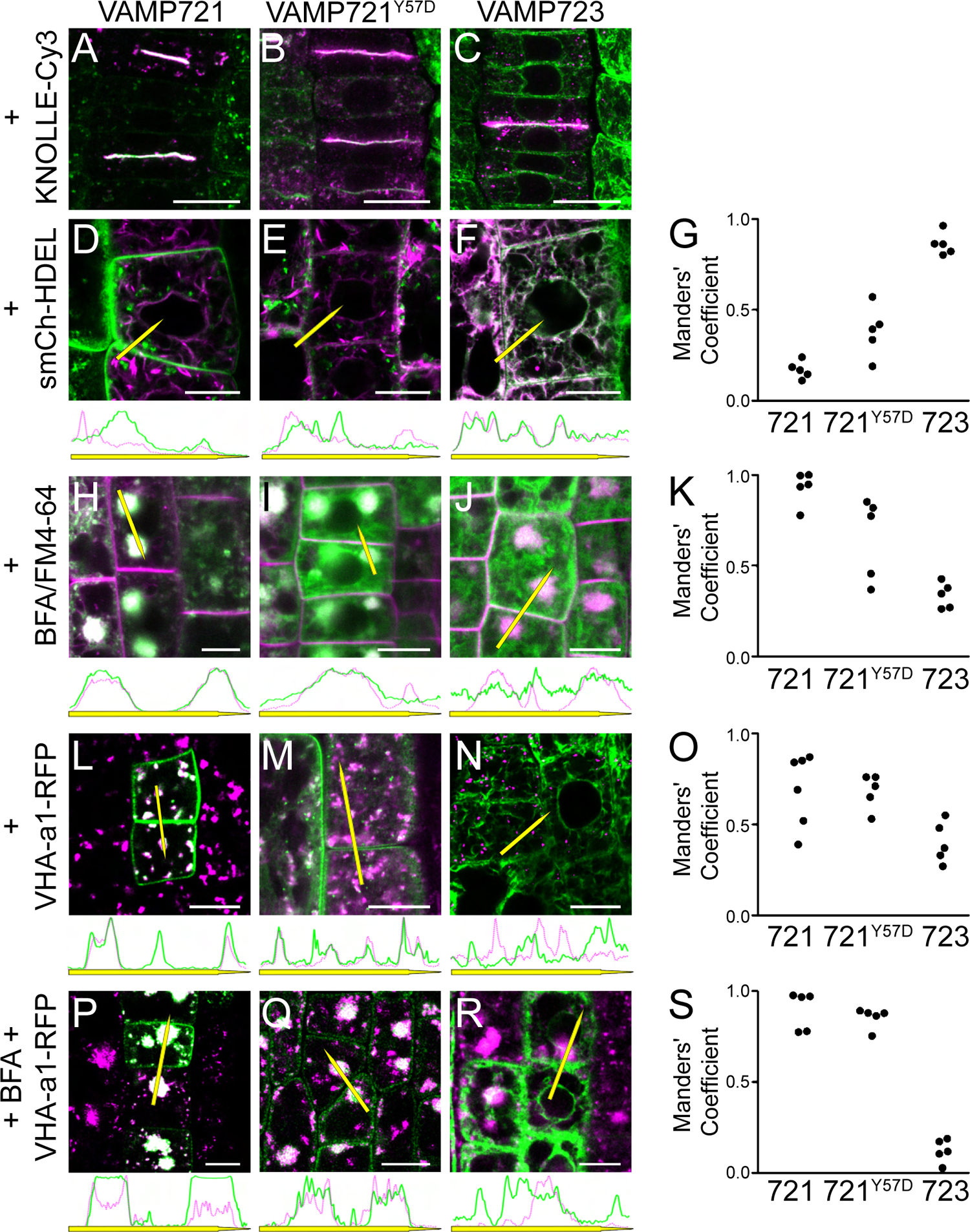
Co-expression of Estradiol-induced GFP-VAMP constructs with organelle marker lines in stably trans-formed Arabidopsis thaliana seedlings. Exemplary confocal triptych images showing from left to right: GFP-VAMP expression, RFP-tagged marker proteins, and the merged images to analyze colocalization with (A-C) the cell plate during cytokinesis using anti-Knolle immu-nostaining, (D-F) the ER using an ER retained, secreted modified (sm) Cherry (smCh-HDEL), (H-J) BFA compartments using FM4-64 staining after Brefeldin-A (BFA) treatment, the TGN using the VHAa1-RFP marker line either (L-N) untreated or (P-R) treated with BFA. (G, K, O, S) Manderś Coefficient was calculated for at least 5 individual cells. Line histograms underneath confocal images show fluorescence intensity along yellow arrows between VAMP constructs and respective marker line. All images were taken 6 hours after induction with 3µM Estradiol. Scale bar 10µm

To validate the localization of the constructs in post-Golgi compartments we applied the ARF-GEF inhibitor Brefeldin-A (BFA). Treatment with BFA (at a 25 µM concentration) leads to the formation of endosomal BFA-compartments which are aggregates of endocytic vesicles and TGN, specifically excluding ER and cis-Golgi (Robinson et al., 2008). Thus, if VAMP723 indeed leaves the ER to traffic along the secretory pathway via the TGN to the cell plate, it should be visible in BFA-compartments. We detected VAMP721 and – more importantly – VAMP721^Y57D^, but not VAMP723, inside of BFA-compartments visualized via accumulation of the endocytosed styryl dye FM4-64 (Figure 3H-K and Supplemental Figure 3G-I). It seems that the Aspartate 57 residue in the LD – as also present in the ER-retained VAMP723 – is not interfering with ER exit and correct localisation of VAMP721 in the endosomal system.

To further assess potential differences in the subcellular localization of the different VAMP721 constructs post ER, we crossed the EDI lines with RFP-tagged reporter lines for the TGN (VHA-a1-subunit) (Dettmer et al., 2006) and treated with mock control or BFA (Figure 3L-S and Supplemental Figure 3J-O). Both, VAMP721 and VAMP721^Y57D^ co-localised significantly with the TGN in untreated samples and in contrast to VAMP723 corroborating the latter’s absence from post-ER compartments. However, VAMP721^Y57D^ shows significantly less correlation with the TGN in comparison to unmutated VAMP721, underpinning its path to degradation compartments (Figure 3L-O, and Supplemental Figure 6J, K). This notion is mirrored by the BFA treatment analysis showing residual accumulation of the Y57D mutant in endomembrane compartments (Figure 3P-S and Supplemental Figure 3M-O).

### VAMP721^Y57D^ overexpression induces vesicular aggregates

During the study of subcellular localization, we noticed that EDI lines expressing VAMP721^Y57D^ showed a tendency for aggregated signal accumulation when expressed for longer time periods (Figure 4A and Supplemental Figure 4). Upon Estradiol-treatment, light-grown plants that strongly expressed VAMP721^Y57D^ were incubated with FM4-64 dye. We observed a partial overlap of the early endocytosed FM4-64 dye with the aggregates within the first 6 hours of staining, specifically surrounding the aggregates (Figure 4B-E). After 24 hours, the FM4-64 dye located at the tonoplast, showing a very weak but homogenous signal in the VAMP721^Y57D^ aggregates (Figure 4F-I). As the seedlings are light-grown, VAMP protein was not detected in the vacuolar lumen, but it is clearly visible that the aggregates are formed outside of the vacuole in the cytosol (Figure 4H and Supplemental Figure 4E). The FM4-64 staining of the tonoplast, however, implies that endocytosis from PM to vacuole is not disrupted – at least not completely – by either overexpression of VAMP721^Y57D^ or aggregate formation.

**Figure 4:**
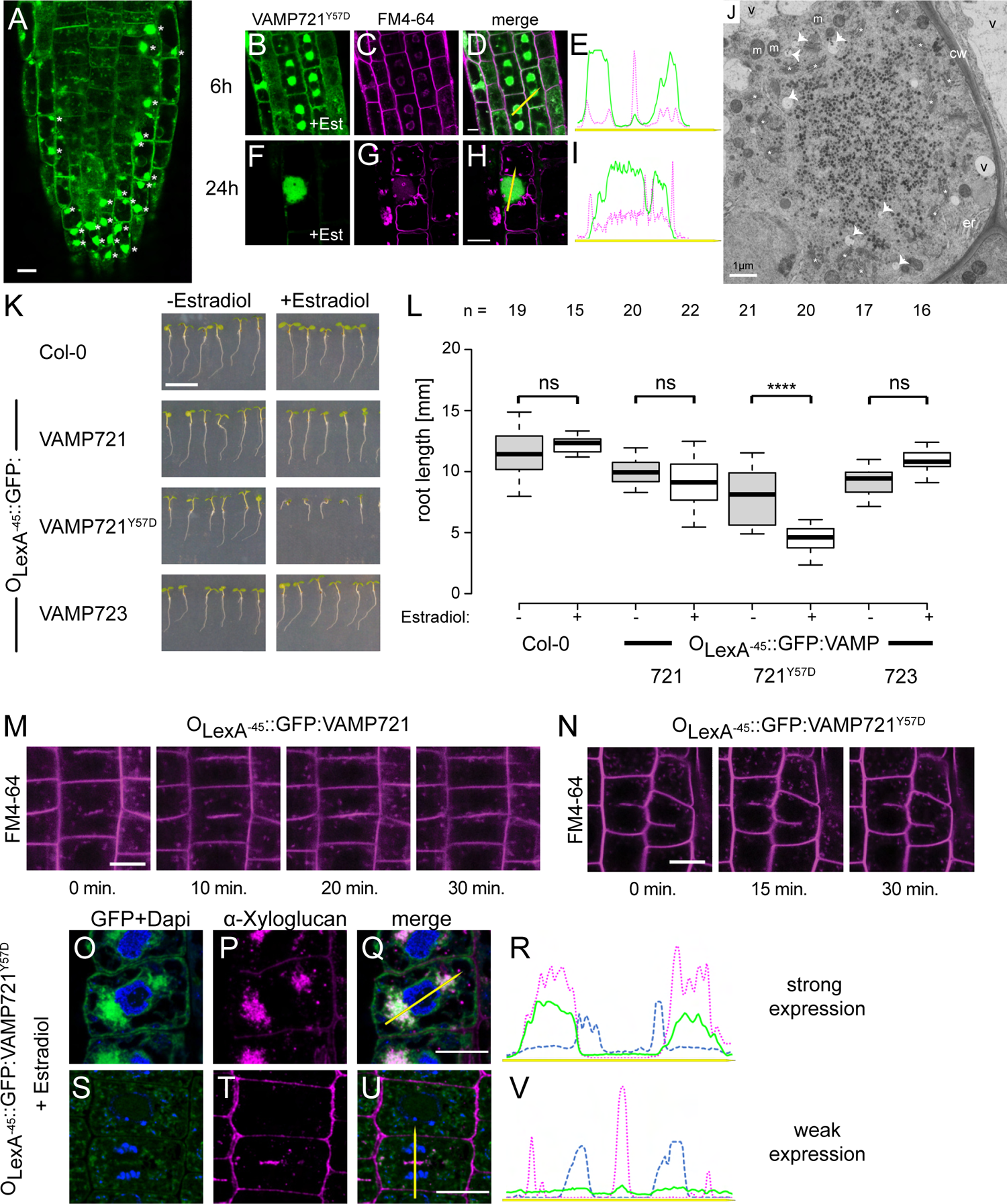
Effects of VAMP721Y57D overexpression. (A) VAMP721Y57D aggregate formation in the rhizodermis after Estradiol induction. Scale bar = 10µm. (B-I) Exemplary confocal images of the co-expression of the styryl dye FM4-64 with VAMP721Y57D (B-E) 6 hours and (F-I) 24 hours after induction with 3µM Estradiol. Line histograms next to the merged confocal images show fluorescence intensity along yellow arrows between VAMP constructs (green line) and FM4-64 (magenta dots). Scale bar = 10µm. (J) Transmission electron microscopy ultrastructural analysis of VAMP721Y57D (over-)expressing cells after cryofixation and upon resin embedding. a = apoplast; cw = cell wall; er = ER; m = mitochondrion; n = nucleus; v = vacuole; asterisk = Golgi stack/TGN; arrow head = MVB. Scale bar 1µm. (K) Exemplary images of roots of EDI seedlings grown either in 3µM Estradiol or DMSO control for 5 days in constant light. Scale bar = 10mm (L) Box plot showing the total root length of different constructs with or without induction. Significant reduction in root growth of VAMP721Y57D expression seedlings is calculated via ANOVA and comparing effect of Estradiol treatment for each line. **** for p<0.0001 and ns for non-significant differences (Bonferroni post-test). Boxplot design: Center lines of boxes represent the median, with outer limits at the 25th and 75th percentiles. Tukey whiskers extend to 1.5 times the interquartile range (IQR).(M-N) Time course confocal images (timing underneath) of an individual cell undergoing cell division in plants inducibly expressing (M) VAMP721 or (N) VAMP721Y57D. Membranes were stained with the styryl dye FM4-64 to visualize the cell plate. Scale bar = 10µm. (O-V) Immunofluorescence labelling of xyloglucan after Estradiol induced expression of GFP-VAMP721Y57D. GFP fluorescence + DAPI, xyloglucan and overlay images of cells showing aggregates. (O-Q) Representative images from cells (400 nm cryosections) showing high GFP signal fluorescence, and (S-U) cells with almost no expression of GFP-VAMP721Y57D. Scale bar = 10µm. (R,V) Line histograms next to the merged confocal images show fluorescence intensity along yellow arrows between VAMP aggregates (green line), xyloglucan accumulation (magenta dots) and Dapi as nuclear counterstain (blue hyphen). Scale bars = 10µm.

Further analyses corroborated the findings from our extensive localization studies: The aggregates do not correlate with the expression of the autophagy marker (Supplemental Figure 4F) and only weakly colocalise with Golgi, TGN or PVC which are found at the periphery of the aggregates (Supplemental Figure 4B-D). However, the resolution limits of light microscopy do not allow to completely dissect the composition of the aggregates as also the ER marker slightly overlaps with it (Supplemental Figure 4A). This may well be signal bleed through from the cell’s periphery above and below the focal plane or ‘overloading’ of the ER due to the strong overexpression of the mutated VAMP.

To overcome the limitation of confocal microscopy, we also used transmission electron microscopy (TEM) to dissect the exact composition of the VAMP721^Y57D^ aggregates. Ultrathin resin sections showed that PVC, TGN and Golgi are trapped in the periphery of the aggregate which seems to be mainly comprised of SV (Figure 4J and Supplemental Figure 5A-C). Immunogold staining (cryosections) showed the presence of VAMP721^Y57D^ mostly within the aggregate and mainly at the vesicular membranes or within the TGN (Supplemental Figure 5D-F). However, strongly expressing cells, also reveal gold particles at the ER and Golgi, corroborating the slight signal overlaps we had observed in our confocal analyses (Supplemental Figure 5E,G).

### VAMP721^Y57D^ overexpression affects cytokinesis and secretion

During the short induction times used for the localization studies the seedlings did not show any relevant phenotype. However, when EDI seedlings were directly germinated on Estradiol, they showed significant growth defects when expressing VAMP721^Y57D^ as compared to wildtype VAMP721 or VAMP723 (Figure 4K-L). Seedling development was impaired on ½ MS agar plates supplemented with 3µM ED, resulting in reduced root growth. Conversely, lines supplemented with DMSO grew normally. We followed dividing cells at the root tip by confocal microscopy using FM4-64 as counterstain. Several cells showed delayed or no progress in cytokinesis, cell wall stubs and abnormal shape in VAMP721^Y57D^ expressing lines, reminiscent of the phenotype of *vamp721^−/−^vamp722^−/−^* double mutants (Zhang et al., 2011) (Figure 4N). Overexpression of wildtype VAMP721, however, did neither cause defects in cytokinesis, nor shorter root growth or other aberrant phenotypes (Figure 4K-M) suggesting that the observed effect is not due to R-SNARE overexpression. In addition, immunoblot analyses of EDI lines showed that VAMP721 levels were much higher under Estradiol-induction compared with the endogenous pVAMP721 driven expression or Estradiol-induced VAMP721^Y57D^ (Supplemental Figure 6).

As both, cell expansion and division, are affected by overexpression of the mutant VAMP721^Y57D^ protein we hypothesize that this mutant imposes a dominant-negative effect on secretion. We therefore used xyloglucan as a secretion marker since it is synthetized at the Golgi apparatus and secreted to the apoplast (Kim and Brandizzi, 2016). Xyloglucan accumulated heavily within cells that showed strong VAMP721^Y57D^ expression (Figure 4O-R), while cells that less strongly expressed the mutant VAMP721 protein secreted normally (Figure 4S-V).

### Interaction of VAMP721 with its cognate partners

Block of secretory traffic can be caused through the use of dominant-negative protein fragments, for example the soluble parts of the Qa-SNARE SYP121 (Grefen et al., 2015). To better understand how the vesicles, accumulate in VAMP721^Y57D^, we tested the interaction capacity of the mutant VAMP721^Y57D^ with exemplary cognate SNARE partners that would usually facilitate vesicle fusion at the plasma membrane or cell plate – the Qa-SNAREs SYP121 (PEN1) and SYP111 (KNOLLE).

We performed co-IP analyses using the stably transformed EDI lines. Seedlings were Estradiol treated and the GFP-tagged VAMP721 constructs were purified via GFP-traps. We tested eluates for the presence of SYP111 or SYP121 using specific antibodies (El Kasmi et al., 2013; Zhang et al., 2017). In both cases, VAMP723 was used as biological negative control and did not interact with neither Qa-SNARE partner; while wildtype VAMP721 served as biological positive control, binding both, SYP111 and SYP121 as had been demonstrated previously (Kwon et al., 2008; El Kasmi et al., 2013). Purification of GFP-tagged VAMP721^Y57D^ immunoprecipitated both Qa-SNAREs, SYP111 and SYP121 to a similar extent as with wildtype VAMP721 (Figure 5A-C).

**Figure 5:**
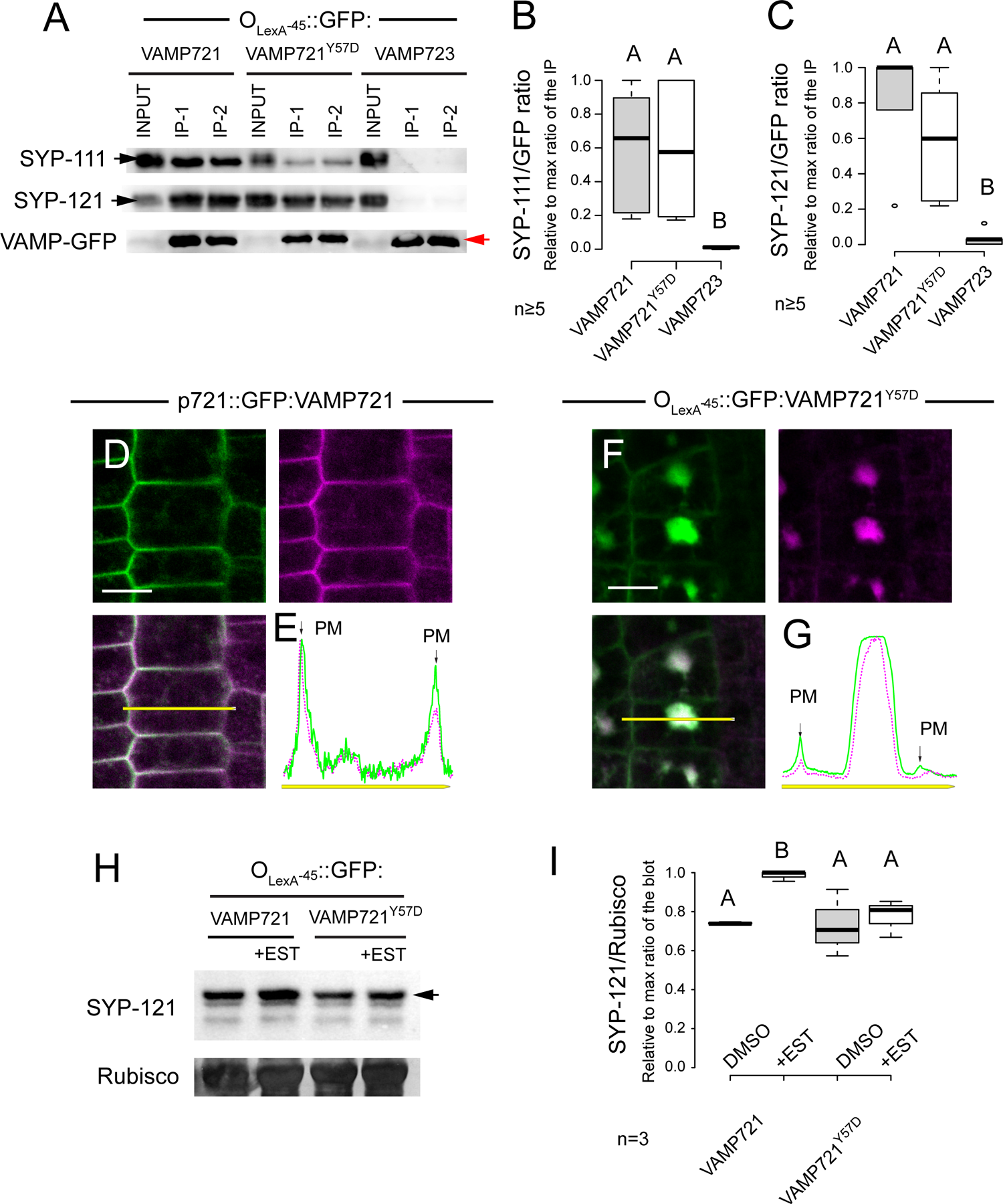
VAMP721Y57D interactions with cognate SNARE partners. (A) Co-IP analyses of Arabidopsis seedlings expressing Estradiol-induced GFP-VAMP721, -VAMP721Y57D, and -VAMP723. GFP-VAMP protein was immunoprecipitated and probed with specific antibodies for (A, B) SYP111 (KNOLLE) (arrowhead indicating the expected 35kDa band) or (A,C) SYP121 (arrowhead indicating the expected 38kDa band). The SYP111/SYP121 to GFP-VAMP (red arrowhead indicating the expected 50kDa band) ratios were normalized to the maximum ratio detected for each row of IP assays to compensate for differences between rows. Statistically different groups are indicated with the letter A or B with n ≥ 5. (D-G) Exemplary confocal images and line histograms of VAMP721 or VAMP721Y57D EDI lines crossed with SYP121-mCherry and induced with 10µM Estradiol for 18 hours. Line histograms show signal overlap between SYP121 and VAMP constructs. Scale bar = 10µm. (H) Representative anti-SYP121 (arrowhead indicating the expected 38 kDa band) immunoblot of 5 days old seedlings’ extracts from the indicated lines treated or not for 24 hours with 3µM Estradiol and their quantification (I).

Since the Y57D mutant readily interacted with SYP121 we questioned whether VAMP721^Y57D^ overexpression might be altering SYP121 localization. To investigate this, we performed crosses between pH4:mCherry-SYP121 (Reichardt et al., 2011) and VAMP721 or VAMP721^Y57D^ EDI lines. We located a large fraction of SYP121 signal colocalizing with the VAMP721^Y57D^ induced aggregates – at the expense of its PM localization – while VAMP721 lines showed normal SYP121 distribution in the PM, Cell plate and SV (Uemura et al., 2004; Rybak et al., 2014) (Figure 5D-G and Supplemental Figure 7A). Trapping of SYP121 only happens in cells with high overexpression of the mutant VAMP and the formation of aggregates (Supplemental Figure 7B). Next, we reasoned that since VAMP721^Y57D^ is actively transported to the vacuole for degradation, the same could happen to its interaction partner SYP121. This, however, was not the case as we could not detect mCherry-SYP121 signal inside the vacuole of EDI lines expressing VAMP721^Y57D^ after dark treatment (Supplemental Figure 7C-E). Immunoblots of these lines did not show any decrease in total SYP121 levels (Figure 5H-I). If anything, SYP121 levels increased slightly in lines overexpressing the wildtype VAMP721.

### Tyrosine 57 is a phosphorylation site

Overexpression of VAMP721^Y57D^ (Figure 4K, L) resembled a previously described effect of sodium orthovanadate (SO; an inhibitor of tyrosine phosphatases) showing reduced root growth among other phenotypes (Yemets et al., 2008). The Y57D mutation supposedly causes a phosphomimetic change – albeit not entirely from a structural point of view – which is why we reasoned that if VAMP721 is phosphorylated, blocking tyrosine dephosphorylation could lead to an increase in phospho-tyrosine at position 57 (pVAMP721^pY57^). We treated *P_VAMP721_*::GFP:VAMP721 and Col-0 lines with SO and observed a strong reduction (15 fold, p<0.0001, n>15) in daily root growth rate after 2 days of SO treatment (Figure 6A). Confocal images showed abnormal cell morphology and incomplete cytokinesis in SO treated p721::GFP:VAMP721 seedlings but did not reveal the formation of large aggregates (Figure 6B-E). It’s important to point out that chemical inhibitors usually have broader effects like in the case of SO that also inhibits plasma membrane ATPases (Palmgren, 2001).

**Figure 6:**
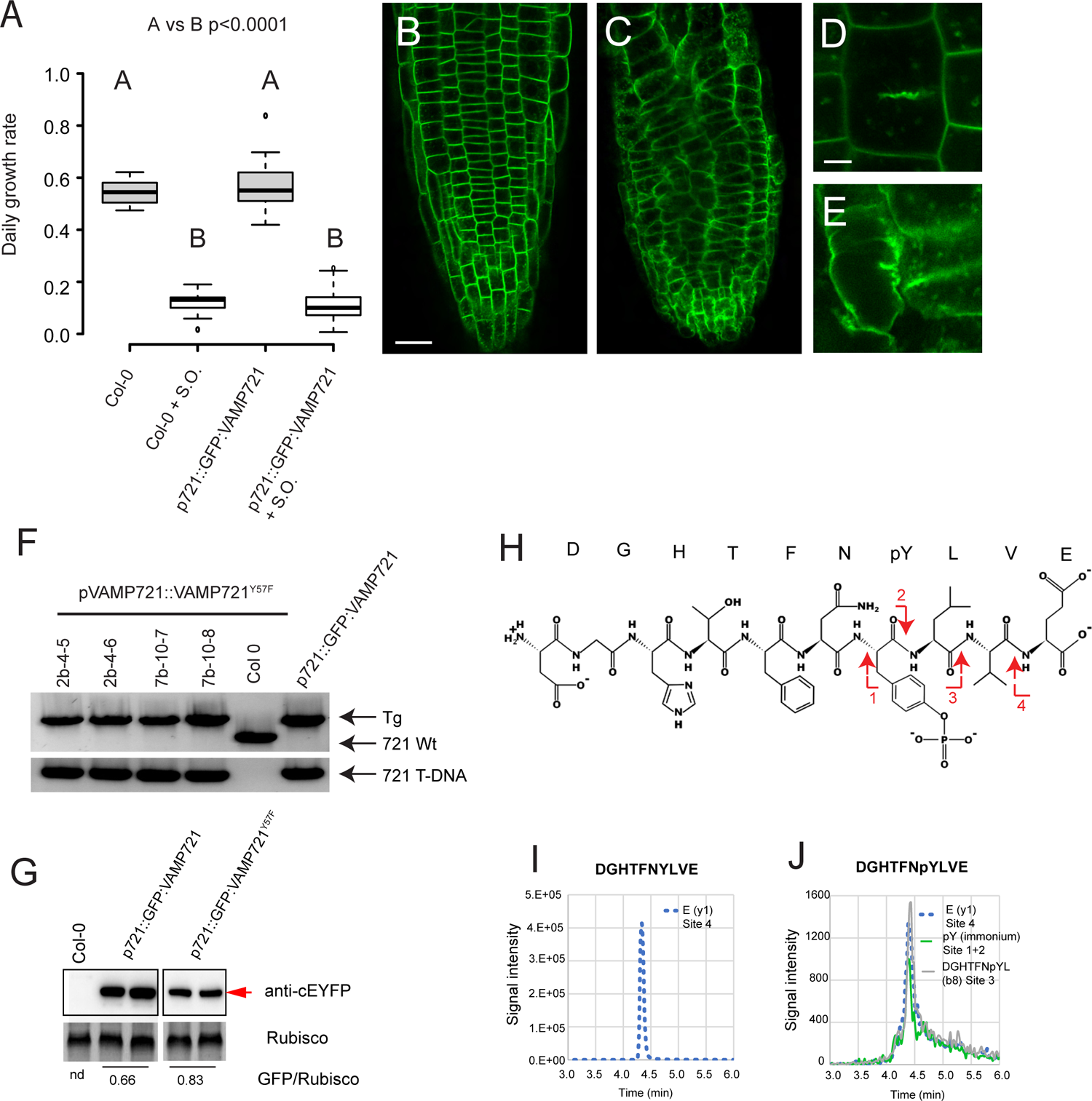
Genetic and biochemical evidence of Tyrosine 57 phosphorylation. (A) Daily root growth of Col-0 and VAMP721 rescued seedlings after two days treatment with or without Sodium orthovanadate (SO). (B-E) Exemplary confocal images of roots of GFP-VAMP721 expressing plants growing either without (B, D) or with the application of SO (C, E). Close-up images show disruption of cell plate formation and aberrant cell shape under the effect of SO (E) in comparison to the untreated plant (D). (F) Genotyping of T3 vamp721−/−vamp722−/− mutants carrying homozygous copies of the VAMP721Y57F transgene under the p721 endogenous promoter. (G) Immunoblot of independent lines expressing VAMP721 and VAMP721Y57F and rescuing the vamp721−/−vamp722−/− phenotype (red arrowhead indicating the expected 50kDa band). (H) Chemical structure indicating the peptide sequence containing the Tyrosine 57 residue after digestion of affinity purified VAMP721 by ASP-N endopeptidase. Fragmentation sites that generated the different species followed in the multiple reaction monitoring (MRM) are indicated with red arrows. Only the phosphopeptide is shown for simplicity. (I-J) Extracted ion chromatograms for the transitions belonging to (I) the DGHTFNYLVE and (J) DGHTFNpYLVE peptides. Cleavage sites that gave rise to the followed species and type of ions (indicated between parenthesis) are marked with red arrows. (B, C) Scale bar = 25 µm, (D-E) Scale bar = 10 µm.

To gain a better understanding of Tyr57 as potential phosphorylation residue we created additional constructs replacing the Tyr^57^ by Phe^57^, an amino acid that shares the aromatic structure of the tyrosine, but lacks its phosphorylatable hydroxyl group (=conservative change). During our initial T1 and T2 screen, we were unable to isolate pVAMP721::GFP:VAMP721^Y57F^ rescued *vamp721^−/−^vamp722^−/−^* lines; instead, we frequently detected plants with the seedling lethal phenotype carrying the mutant construct (Supplemental Figure 8). However, when analysing T3 decedents of segregating complemented lines, we isolated several plants deriving from different independent T1 lines that showed a strong GFP signal within the *vamp721^−/−^vamp722^−/−^* background (Figure 6F). Lack of an initial rescue in T1 lines suggests that the hemizygous gene dosage of the mutated VAMP721^Y57F^ might not be sufficient to overcome the lack of wildtype VAMP721. In terms of protein abundance our immunoblot assays suggest that plants rescued by the mutant VAMP721^Y57F^ require slightly higher protein abundance than the wildtype version (Figure 6 G). The blot also implies that the Y57F mutation does not destabilise the protein.

Taken together, our biochemical and genetic assays suggest that phosphorylation of Y57 might play a role in VAMP721 regulation. Therefore, we focused on the detection of such phosphorylation. For this, we developed a customized UPLC-MS strategy using synthetic peptide standards (DGHTFNYLVE and DGHTFNpYLVE) to adjust the conditions for the detection of the phospho-tyrosine containing peptide. We purified GFP-VAMP721 protein from stable transgenic lines treated with SO during induction. The purified protein was digested with ASP-N before the MS-MS run. The DGHTFNYLVE peptide that contains the Tyr^57^ (Figure 6H) was analysed detecting the species common for both the phosphorylated and unphosphorylated peptides used as internal control as well as phosphorylation specific species for each peptide (Figure 6I-J). The detection was very challenging due to the low signal of the ions used for the pY detection, but we could reliably identify phosphorylation of the residue. Considering this limitation in abundancy, we qualitatively estimated a ratio of 30 Tyrosine residues to each phosphorylated Tyrosine.

### VAMP721 and VAMP721^Y57D^ interactome highlights common SNARE related interactors and supports differences in VAMP721^Y57D^ localization and stability

Detecting – albeit very low levels of VAMP721^pY57^ – we hypothesise that residue 57 is a phosphorylation site which enhances VAMP721 activity and needs to be quickly dephosphorylated to avoid the dominant negative effects that we observed in the phosphomimic VAMP721^Y57D^. Due to the short-lived phosphorylation of VAMP721, and to gain further knowledge regarding its biological function, we used VAMP721^Y57D^ in comparison to VAMP721 and aimed to identify possible differential interactions. We performed additional co-IP experiments with VAMP721, VAMP721^Y57D^ and soluble GFP as a background control, followed by tandem mass spectrometry analysis. As expected, the most abundant peptides corresponded to the precipitated protein VAMP721 or VAMP721^Y57D^. We were also able to detect the well described partners SYP121 and SNAP33 (Kwon et al., 2008) along with several other SNARE and SNARE related proteins (Supplemental Table 1). Other known interactors like SYP111 were detected, but filtered out due to iBAQ values below our determined threshold (Supplemental Table 1). Using iBAQ VAMP721/VAMP721^Y57D^ ratios we ranked the identified proteins according to their preferred interaction with either VAMP721 (ratio > 3), VAMP721^Y57D^ (ratio < 0.33) or intermediates (ratio between 0.33 and 3.0). The list of preferred VAMP721 interacting proteins comprises mostly SNARE and SNARE-related proteins such as the core Qbc SNARE SNAP33 as well as the SNAREs SYP61, VTI12, NPSN11 and NPSN12 which form complexes with KNOLLE/SYP111 at the cell plate (Sanderfoot et al., 2001; Zheng et al., 2002; Drakakaki et al., 2012; Karnahl et al., 2017). VAMP721 also co-immunoprecipitated SNAP2 (ASNAP1 or ASNAP2, and GSNAP2), a homolog of the yeast Sec17p/α-SNAP which activates ATP hydrolysis by NSF and the disassembly of the cis-SNARE complex (Carter et al., 2004; Jun and Wickner, 2019). We also found the Sec1/Munc18-like (SM) protein SLY1 a known SNARE chaperone required to prime the SNARE proteins before complex formation (Kosodo et al., 2003), preferentially with VAMP721 at a ratio of 3.6.

The group of proteins that were identified with both VAMP721 constructs ranges from more SNARE proteins (ratios around 1.5-2.9) to membrane proteins such as plasma membrane ATPases, ABC transporters and Aquaporins (ratios between 0.38-1.3). A number of proteins related to protein folding and stability regulation were found to preferably bind to VAMP721^Y57D^, in particular chaperoning and proteasome-related proteins. This reinforces the idea of an increased degradation of VAMP721^Y57D^. The RPT proteins (26S proteasome regulatory subunits) (1A, 2A/B, 5A/B) as well as the RPN proteins (26S proteasome non-ATPase regulatory subunit) (5A/B, 6, 7,10) showed a clear preference for VAMP721^Y57D^. There is also favoured interaction with several ERAD related proteins like BAG6 (part of BAG6/BAT3 complex), TIF4A (part of the Hrd1 complex (Lin et al., 2019) and DNAJ proteins (e.g. ERDJ2A and ERDJ3B) which aid in handling misfolded proteins via HSP70 at the ER (Liu and Li, 2014). We also identified several other chaperones (HSP70-2 or 18, HSP70-4, HSP70-9, HSP90-2, HSP90-5, USP17, CCT8) and accessory chaperoning proteins related to their activity (ATJ2 and ATJ3 activators of HSP70 and MEC18.18 activator of HSP90). Also, trafficking related proteins like SYNAPTOTAGMIN 1 & 7 (SYT1, SYT7) (Yamazaki et al., 2010) and ARF-like GTPase ARL8b (Nishikiori et al., 2011) bound more strongly to the mutated R-SNARE.

## Discussion

The main chore of VAMP721/722 is to facilitate the final step of vesicle trafficking – membrane fusion. To achieve this, SNAREs must align precisely in a parallel manner and assemble into a four-helix bundle (trans-/cis-complex). SNAREs on their own, however, would tend to form kinetically trapped dead-end structures which are unable to drive fusion (Brunger, 2006). To prevent this, additional autoinhibitory mechanisms evolved to block SNARE interaction until required for fusion. For example, both Qa- and R-SNAREs not only feature additional domains that can fold back onto the SNARE motif (Vivona et al., 2010), but have also been shown to interact with additional proteins that can lock onto such ‘closed’ confirmation.

A prominent example of this mechanism in plants is the Sec1/Munc18 (‘SM’)-protein KEULE, named after its club-shaped *loss-of-function* phenotype which is testament to its vital role during cytokinesis (Jurgens et al., 2015). Here, the mitosis-specific Qa-SNARE KNOLLE/SYP111 facilitates establishing of the cell plate through homotypic vesicle fusion (Karnahl et al., 2017). This process is tightly regulated through binding of KEULE to KNOLLE (Park et al., 2012). While regulating (SM) proteins have been shown to act on Qa-SNAREs in animal and plant systems, R-SNARE regulating proteins have only been described in mammalian cell culture. Binding of the endosomal VARP to the ‘closed’ mammalian VAMP7 slows down SNARE-SNARE binding by a factor of 65 preventing premature complex formation and in turn exerting temporal control over fusion events (Schafer et al., 2012).

Current models propose that after tethering, the SNAREs are opened or “primed” to form the four-helix (=SNARE-motifs) bundle. SM proteins play a crucial role because not only do they prevent Qa-SNAREs to open prematurely, but they are also able to bind them in an open configuration thereby presenting an additional site for R-SNARE binding (Zhang and Hughson, 2021). The R-SNARE itself must be in an ‘open’ state to interact with both Qa and SM proteins. VAMP7 can spontaneously adopt it’s open configuration and be further activated by phosphorylation of the Y^45^ residue within its LD (Burgo et al., 2013).

We provide evidence that a Tyrosine residue at position 57 within the Arabidopsis VAMP721 protein – which is not corresponding to the conserved Y^45^ of the human VAMP7 residue (Figure 1A) – can be phosphorylated and thereby facilitate R-SNARE opening and activation, potentially leading to a faster trans-SNARE assembly and subsequent cis-complex formation after membrane fusion. Following complex disassembly by the NSF/α-SNAP chaperone complex, dephosphorylation is required to allow proper backfolding of the LD. According to this model – and in line with the low amount of pY^57^ that we detected – phosphorylation probably lasts only for the short time window between formation of the trans-complex and until disassembly of the post-fusion cis-complex. Failure to rescue the cytokinesis defect of the *vamp721vamp722 loss-of-function* lines and toxicity of its overexpression in wildtype *Arabidopsis* plants demonstrate the physiological defects that the mutated, constitutively opened VAMP721^Y57D^ imposes on vesicle traffic. It also underpins that dysregulation of LD opening outcompetes wildtype copies of VAMP721.

One way for the plant to cope with such misfolded VAMPs seems to be degradation. In our work we showed that this artificially and constitutively opened R-SNARE is recognized by cytosolic HSP70 isoforms and other chaperones as a misfolded protein and subsequently poly-ubiquitinated (Figure 2C-F). The interaction with ER-associated protein degradation (ERAD) related proteins (Supplemental Table 1) and proteasome subunits, point towards proteasomal degradation, while the interaction with several vacuole-resident proteins (Supplemental Table 1) as well as the localisation inside PVC and vacuole support the possibility of vacuolar degradation, too (Figure 2 G,I,Q-R). In other words, after ER insertion a fraction of the protein may be recognized as misfolded and subsequently get degraded via ERAD, while the remainder exits the ER to traffic along the secretory pathway where again, the protein is recognized as misfolded and sent to the vacuole for degradation. This demonstrates that the plant cell attempts to get rid of the mutated protein all along the secretory pathway.

In contrast to the Y57D mutation, the Y57F mutation that neither mimics nor allows phosphorylation at that residue, was able to rescue the *vamp721 vamp722* double mutant; however, only as homozygous transgenic, in other words, at a higher gene dosage, whereas wildtype VAMP721 already rescues in heterozygous transformants. This means that in case of the Y57F mutation, fusion with cognate SNARE partners is not completely prevented but rather left to a chance event, reducing the R-SNAREs fusogenic activity which would explain the need for higher amounts of the mutated VAMP721^Y57F^ to fulfil its housekeeping functions. It is possible that the effect is not as strong since other, additional (phosphorylation) sites (for instance Y^48^) might co-regulate the opening mechanism of the VAMPs LD as is the case for Y^45^ of the human VAMP7 (Burgo et al., 2013). Potentially, pY^57^ may gain more relevance under special developmental/environmental conditions where fine-tuning of vesicle trafficking is required. Previous work demonstrated that environmental conditions such as biotic and abiotic stresses can alter abundance of *VAMP721/722* proteins considerably (Kwon et al., 2008; Yun et al., 2013).

Protein levels do seem to be a relevant factor in these fine-tuning of trafficking and degradation events. While expression from the native promoter resulted in virtually complete degradation, the artificial, inducible overexpression of the Tyrosine-to-Aspartate mutation caused drastic cellular trafficking defects resulting in severe phenotypic consequences (Figure 4): Default secretion/recycling is blocked inducing the formation of aggregates. Though trafficking of VAMP721^Y57D^ from ER to TGN is unaffected since the protein was able to reach post TGN locations (PM, cell plate, endosomes) and is neither retained at the ER nor Golgi. This is not surprising since traffic of membrane proteins along these initial compartments of the secretory route depends on the R-SNARES SEC22 in ER-to-Golgi traffic (El-Kasmi et al., 2011) and YLT61/62 in Golgi-TGN traffic (Chen et al., 2005). The aggregates we observed comprised SV with Golgi and TGN in close proximity suggesting that the VAMP721^Y57D^ induced blocking occurs most likely at post TGN compartments. It is tempting to speculate that during vesicle approach to the PM the phosphomimetic Y57D in its constitutively opened state interacts prematurely with its partner Q-SNAREs residing on other vesicles before proper tethering of the vesicle to the PM. This would block secretion, while keeping the vesicles physically docked eventually leading to the large number of accumulated SV that we observed in VAMP721^Y57D^ induced aggregates.

The fact that we find SYP121 co-localised in the aggregates underpins the notion that VAMP721 could also trap its cognate SNARE partners in – potentially – non-fusogenic complexes. Though limited by the observation of 100nm sections, we repeatedly detected structures that suggest SV docking but failure to fuse within the aggregates. Under normal circumstances vesicular aggregates appear during cytokinesis as prelude to homotypic vesicle fusion which was well described during cell plate formation. Here, VAMP721 and its mitotic-phase partners SYP111/SNAP33 or SYP111/SYP71/NPSN11 are trafficking together as cis-SNARE complexes (Karnahl et al., 2017). These cis-SNARE complexes are then disassembled by NSF and SNAP to favour the later formation of trans-SNARE complexes that can drive membrane fusion (two trans-SNARE complexes formed from two original cis-complexes) (Karnahl et al., 2017).

This could support a model whereby the VAMP mutation leads to an initial homotypic vesicle fusion within the forming aggregate, however, during cis-complex disassembly, VAMP721^Y57D^ is unable to mask its LD by folding back on it (Zhang et al., 2017). This in turn leads to a prolonged state of cis-complex interaction blocking function of the involved SNAREs as well as preventing individuals of these complexes to participate in further fusion events. It would also explain why the mutant R-SNAREs seem to outcompete their wildtype relatives – probably through sheer number and for being in an activated state trapping cognate SNARE partners in either dysfunctional antiparallel trans-SNARE or constitutively reformed cis-SNARE complexes.

Another possibility, however, would be that the dominant-active VAMP titrates out non-SNARE interaction partners such as tethering or docking factors thereby inhibiting vesicle fusion. Indeed, our IP-MS highlighted ARL8b and SYT1 as potential candidates to be preferably bound by the mutated Y57D. ARL8b is an ADP Ribosylation Factor Like GTPase highly conserved (Khatter et al., 2015) and is involved in bidirectional lysosomal trafficking in mammalian cells (Wu et al., 2020). Hence, titration of ARL8b could lead to the vesicular aggregates observed that are reminiscent of BFA-compartments.

Another potential candidate could be SYT-1 that has an important role in Ca^2+^ activated exocytosis in neurons. Two modes of action have been postulated, clamp and release. While the former blocks fusion of docked SVs with the PM in absence of Ca^2+^, sensing of local Ca^2+^ increase leads to release of the block and fast fusion (Tagliatti et al., 2020). It may be that the dominant-active binding of VAMP721^Y57D^ to the Synaptotagmin maintains SYT1 in its clamp function, preventing vesicle fusion indefinitely.

Either way, and as a result, all secretory cargo including xyloglucan will stay trapped within the aggregates eventually leading to the demise of the plant if *de novo* synthesis of the mutant SNARE is maintained. Otherwise (under native or lower expression levels), the various degradation pathways attempt and succeed to clean the cell of the toxic mutant. Taken together, our data suggests the Tyrosine 57 residue within the LD to act as a ‘phospho-switch’ that regulates the VAMPs activity and stability, possibly by way of modulating the open-closed state of the R-SNARE.

## Materials and Methods

### Plants material and growth conditions

The Arabidopsis mutant lines used in this work are all in the Columbia-0 genetic background. Plants were grown in soil at 22°C and long day light conditions (16/8h light/dark). For plate assays, individual seeds were grown under constant light conditions in 120 mm square plates filled with 70 ml of half strength MS media and 1% agar. Transgenic lines were generated using the standard floral dipping protocol (Clough and Bent, 1998) using Agrobacterium tumefaciens strain GV3101. Plants were selected in soil by BASTA spraying or in plants with hygromycin according to the fast selection method (Harrison et al., 2006). The *vamp721^−/−^ vamp722^−/−^* T-DNA insertion lines correspond to accessions SALK_106594 and SALK_103189, respectively, and were characterized in (Kwon et al., 2008). Genotyping was performed using primer combinations indicated in Supplemental Figure 8 and Supplemental Methods.

### Molecular cloning and construct design

The entry clones were generated by PCR amplification of the insert followed by PB recombinase reaction in the specified vectors. Site-directed mutagenesis was performed as previously described (Karnik et al., 2013) to obtain the different mutant variants. The final destination clones used for plant transformation were generated by LR recombination in case of the EDI lines and by restriction ligation procedure for the lines with the endogenous promoter. Cloning details and vector maps with annotations are available in the supplemental methods and Supplemental Table 2.

### Expression analyses via real-time qPCR

RNA extractions were performed with Universal RNA extraction Kit from Roboklon. After RNA quantification with nanodrop and running the RNA for integrity assessments, ProtoScript II reverse transcriptase from NEB was used to generate cDNA. Real time reactions were performed using Luna® Universal qPCR Master Mix from NEB. Oligonucleotides were used to detect GFP-VAMP (56 and 57) and Actin2 (58 and 59). Quantification was performed with the software Linereg.

### Co-IP and MS analyses

Induction of protein expression in EDI lines was performed by flooding the plates in 1/2 strength MS supplemented with 3µM Estradiol and subsequent removal of the solution after 5 minutes. The remaining liquid with Estradiol was left for 6h before harvesting the seedlings. We used up to 4g of tissue as starting material from which co-IP experiments were performed as previously described (Karnahl et al., 2017), using 7 days old seedlings. Samples were eluted from the GFP-trap beads in Laemli buffer 2x for immunoblot or send directly before elution for MS-MS analysis at the University of Tübingen Proteome Center as described previously (Asseck et al., 2021).

### Detection of phosphorylation

We performed an IP with the following modifications. The washes were more stringent and carried 0,5% SDS to remove all interacting proteins. Seedlings were treated with 250µM SO during the induction, and all buffers used for the IP also included 1mM SO. Samples were analyzed with a UPLC-MS protocol further described in Supplemental Methods using the peptides DGHTFNpYLVE and DGHTFNYLVE as standards.

### Laser Scanning Confocal microscopy

Microscopy images of 6d old seedlings were acquired with either an SP8 (Leica) of LSM880 (Zeiss) confocal microscope using 63x NA1.2 water immersion objective. Excitation/emission parameters for GFP and RFP were 488 nm/490 to 552 nm and 561 nm/569 to 652 nm, respectively. Sequential scanning mode was used for colocalization of both fluorophores. Unless stated otherwise all inductions with Estradiol were performed for 6h placing the seedlings into a new plate supplemented with 3µM Estradiol. For all co-localization analyses, we used F1 seedlings derived from the different crosses with marker lines ER/PM: CD3-959, Golgi: SYP32, TGN: VHAa1, Tonoplast: VAMP711, autophagosome: NBR1, and PVC: ARA7, pH4:SYP121 (Dettmer et al., 2006; Nelson et al., 2007; Geldner et al., 2009; Reichardt et al., 2011; Hafren et al., 2017). Seedlings were treated for 1 hour with 20 μM Wortmanin, 25 μM BFA or 2 μM FM4-64 dye prior imaging. Immunogold and immunofluorescence details in supplemental methods.

### Immunoblots

Seedling samples were ground to a fine dust with tissuelyzer II (Qiagen) and later extracted with UREA buffer (200mM Tris HCl pH6.8, 10% SDS, 6M Urea, 20%, Glycerol, 50mM DTT, 0.05% Bromophenol Blue). Samples were heated for 10 minutes at 80°C and run in standard 12% SDS-PAGE gels and transferred into PVDF membranes for immunoblotting. As primary antibodies we used anti-SYP111 (1:6000, rabbit) (Lauber et al., 1997), anti-SYP121 (1:4000, rabbit) (Zhang et al., 2007), anti-HSP70 cytosolic isoform (1:10000, rabbit) from Agrisera (AS08 371), anti-Ubiquitin (1:1000, mouse) from Santa Cruz Biotech (sc-8017) and anti-cEGFP (1.2000) from Agrisera (AS11 1775).

### Ultrastructural TEM analysis

We used 6 days old seedlings induced for 18 hs with 3µM ED. Seedling root tips were high-pressure frozen, freeze-substituted in acetone containing 2.5% OsO4 and embedded in epoxy resin (for details see Singh et al, 2018). Ultrathin sections were stained with uranyl acetate (in 50% ethanol) and lead citrate and viewed in a JEM-1400plus (Jeol) transmission electron microscope. Images were taken with a 4K CMOS TemCam-F416 camera (TVIPS) and processed using Adobe Photoshop CS5.

## Author contributions

MR, VL, GJ and CG designed all experiments. MR, NW, CC, DM, SR, JM, IG, ER, YS performed experiments. MR, SR, ER, YS, VL and CG analysed the data. MR and CG wrote the manuscript with input from SR, YS, VL and GJ. All authors approved the final version of this paper.

## Acknowledgements

We thank Misoon Park for useful advice in the co-IP procedure and immunoblots with SYP111 and SYP121 antibodies, Rebecca Stahl for technical support and Suayib Üstün for providing RFP-NBR1 seeds. The MS analysis was done at the Proteomics Centre of the University of Tübingen, and we are grateful to Mirita Franz-Wachtel for her help in interpreting the data.

---

**Supplementary Figure 1:**
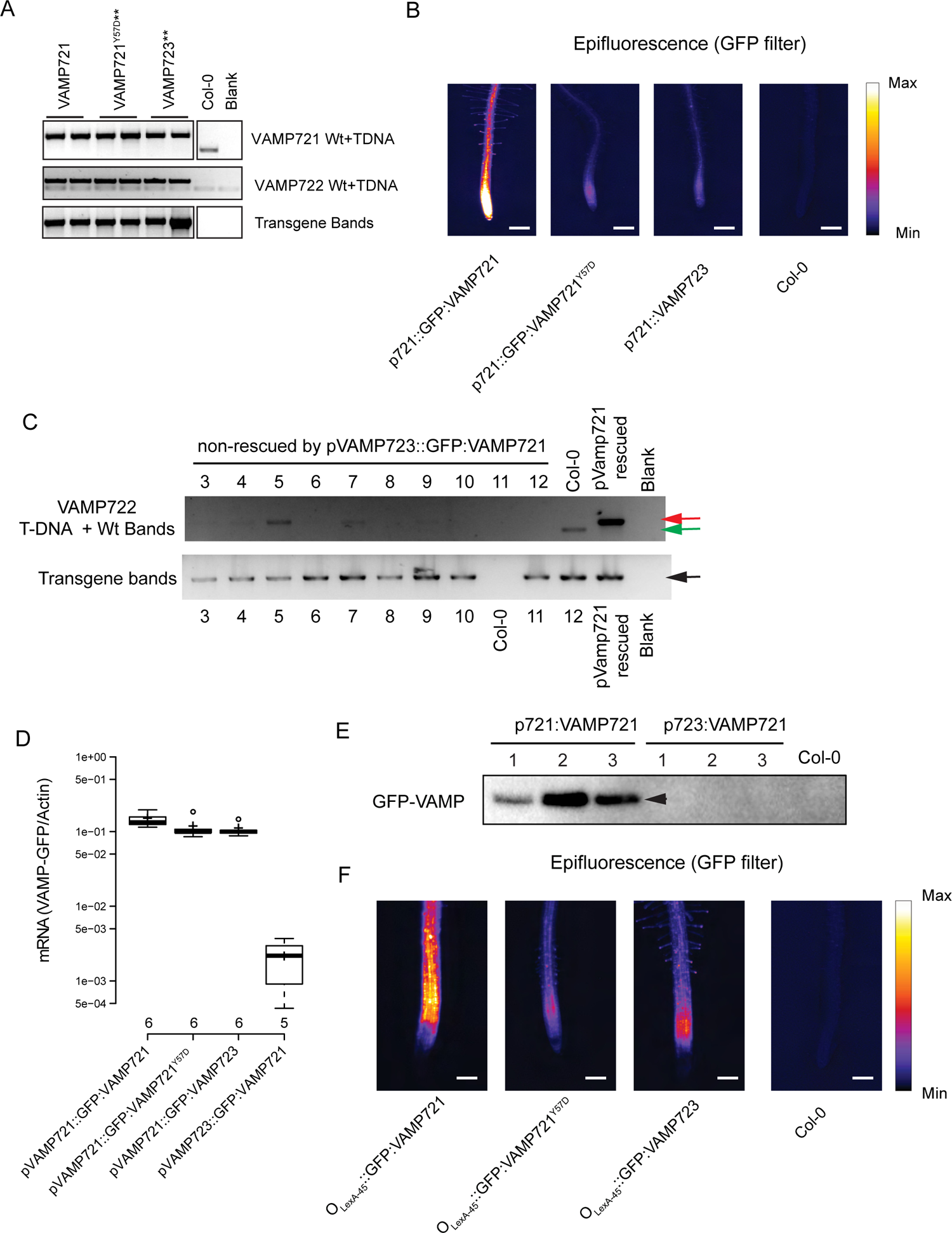
Genotyping and expression analysis of VAMP constructs. (A) Genotyping of complementation assays. PCR amplification of T-DNA and Wt alleles for VAMP721 (top panel), VAMP722 (middle panel) and transgenic constructs (bottom panel ‘transgene bands’ is identical to the data in Figure 1B) ** indicates mutant seedling phenotype. (B) Representative epifluorescence pictures of GFP-VAMP transformed lines used for quantification in Figure 1D. Intensity-based false-color labelling is based on the“fire” look up table (LUT) of the ImageJ software. (C) Genotyping of T2 seedlings from 10 independent T1 lines transformed with pVAMP723::GFP:VAMP721 in the vamp721−/−vamp722-/+ background that display the mutant phenotype compared to wildtype Col-0 and a VAMP721 complemented line. Red (T-DNA), green (WT) and black (Transgene) arrows indicate the PCR bands for VAMP722 T-DNA, VAMP722 Wt or transgene detection. (D) qRT-PCR assessment of VAMP723 promoter strength in comparison to data shown in Figure 1C. (E) Immunoblot (anti-cYFP) of 3 independently transformed seedlings expressing GFP-VAMP721 either from the native or VAMP723 promoter (expected size at 50kDa marked with black arrows). (F) Representative epifluorescence pictures of GFP-VAMP transformed lines and induced using Estradiol induction that were used for quantification in Figure 2B. Intensity-based false-color labelling is based on the “fire” look up table (LUT) of ImageJ. Scale bar = 200µm.

**Supplementary Figure 2:**
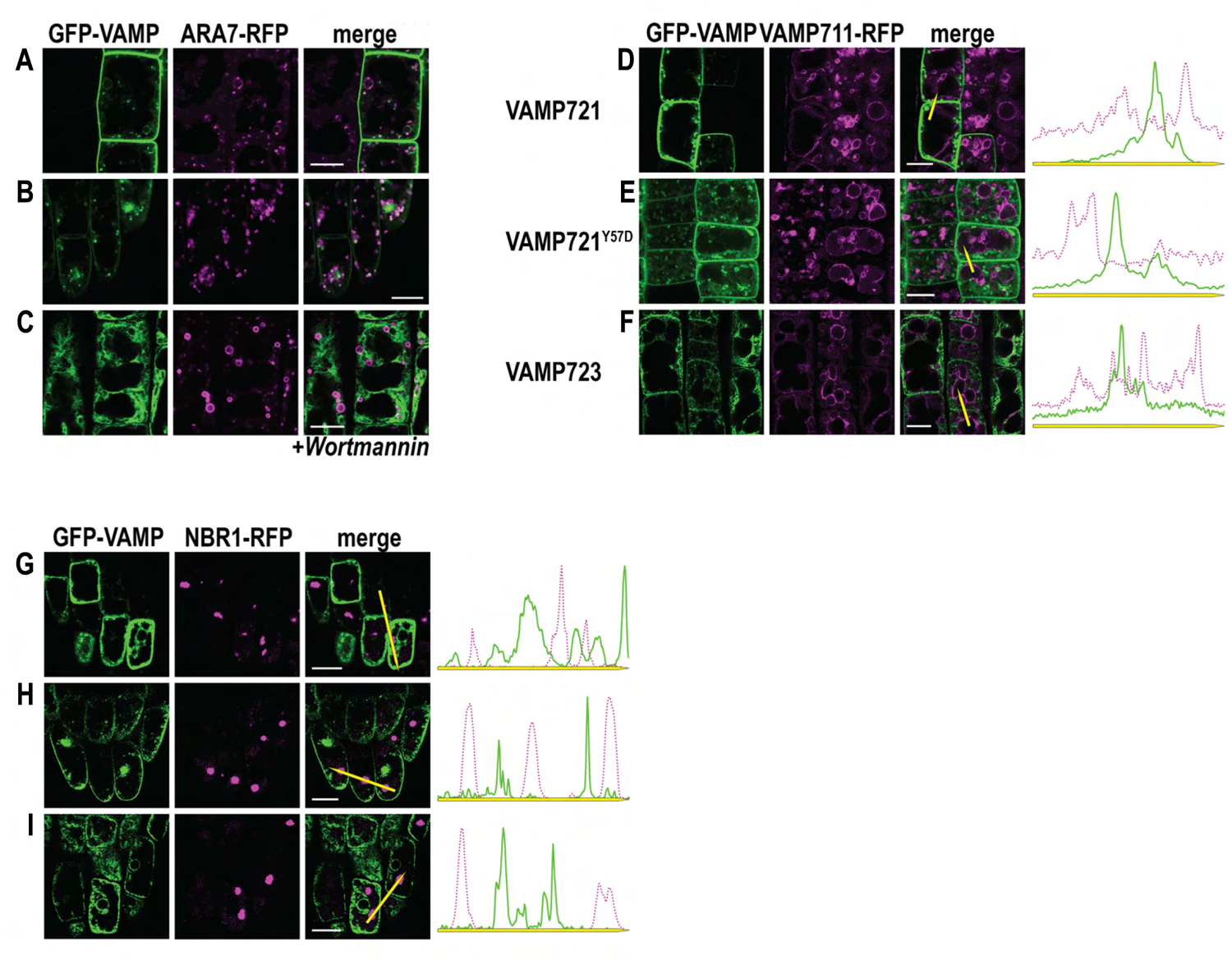
Co-expression of Estradiol-induced GFP-VAMP constructs with organelle marker lines in stably transformed Arabidopsis thaliana seedlings. Representative confocal triptych images showing from left to right: GFP-VAMP expression, RFP-tagged marker proteins, and the merged images to analyze colocalization with (A-C) the PVC using the ARA7-RFP marker line treated with Wortmannin, (D-F) the vacuole using the tonoplast marker RFP-VAMP711, or (G-I) the autophagosome using the NBR1-RFP marker line. Line histograms in D-I show fluorescence intensity along yellow arrows between VAMP constructs and respective marker line. All images were taken 6 hours after induction with 3µM Estradiol. Scale bar = 10 µm.

**Supplementary Figure 3:**
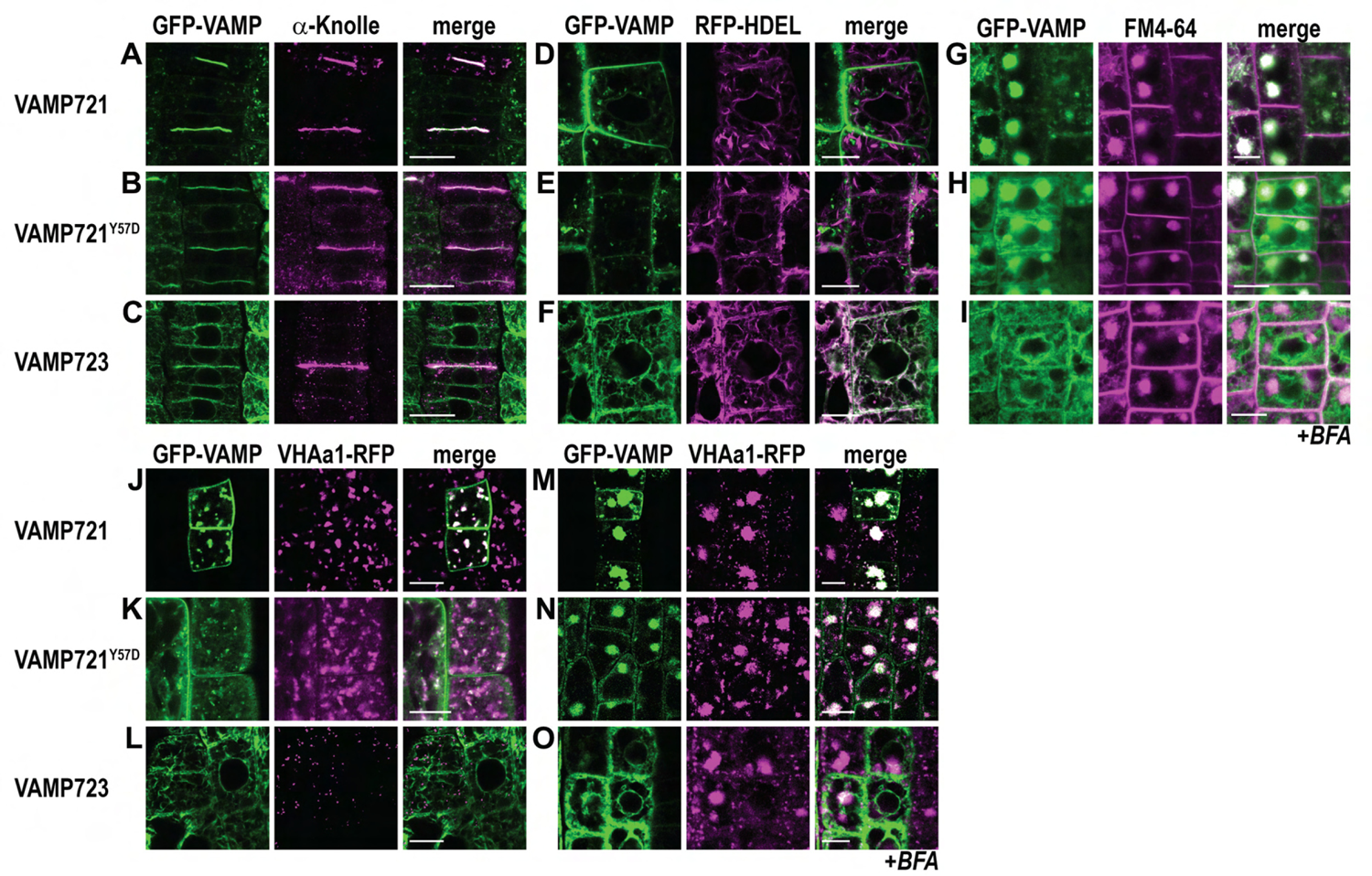
Co-expression of Estradiol-induced GFP-VAMP constructs with organelle marker lines in stably transformed Arabidopsis thaliana seedlings. Representative confocal triptych images showing from left to right: GFP-VAMP expression, RFP-tagged marker proteins, and the merged images to analyze colocalization with (A-C) the cell plate during cytokinesis using anti-Knolle immunostaining, (D-F) the ER using secreted RFP-HDEL, (G-I) BFA compartments using FM4-64 staining after Brefeldin-A (BFA) treatment, (J-L) the TGN using the VHAa1-RFP marker line either (J-L) mock-treated, or (M-O) treated with BFA. All images were taken 6 hours after induction with 3µM Estradiol. Scale bar = 10 µm.

**Supplementary Figure 4:**
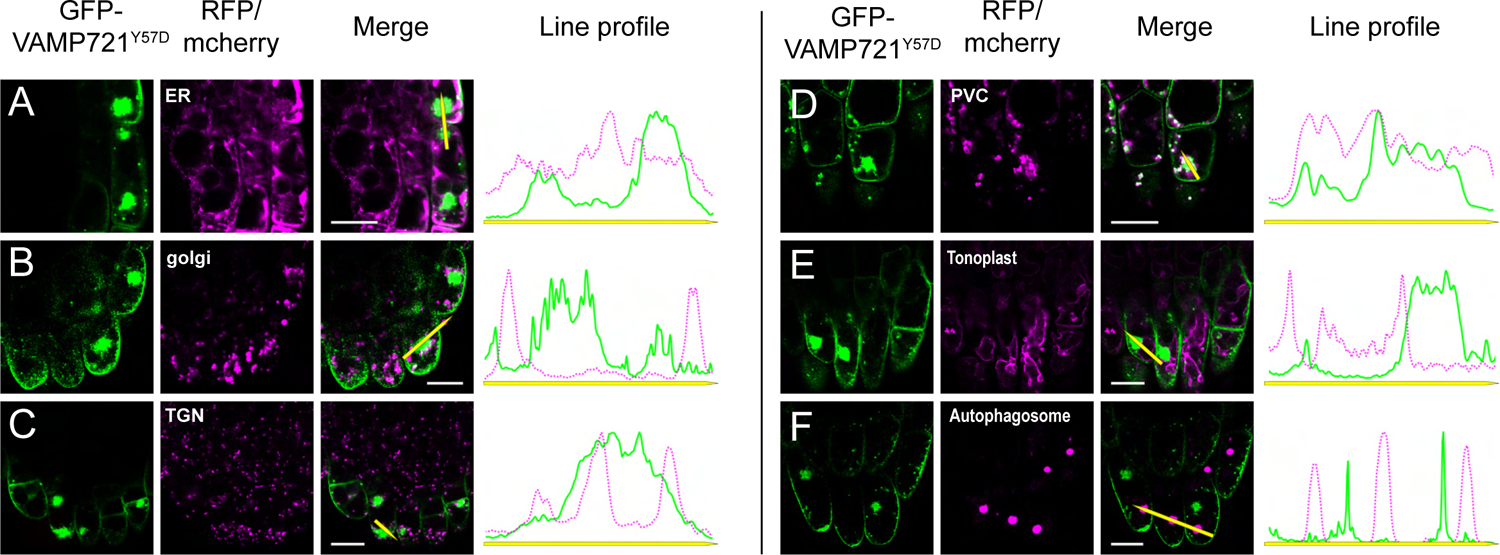
Aggregate colocalization with organelle markers. Representative confocal triptych images showing from left to right: GFP-VAMP expression, RFP- or mCher-ry-tagged marker proteins, and the merged images to analyse colocalization of aggregates with marker for (A) the ER, (B) Golgi, (C) TGN, (D) PVC, (E) Tonoplast, and (F) autophagosome. VAMP721Y57D seedlings were analyzed after 6 hours of induction with Estradiol. Line histograms next to the merged confocal images show fluorescence intensity along yellow arrows between VAMP constructs (green line) and FM4-64 (magenta dots). Scale bar = 10 µm.

**Supplementary Figure 5:**
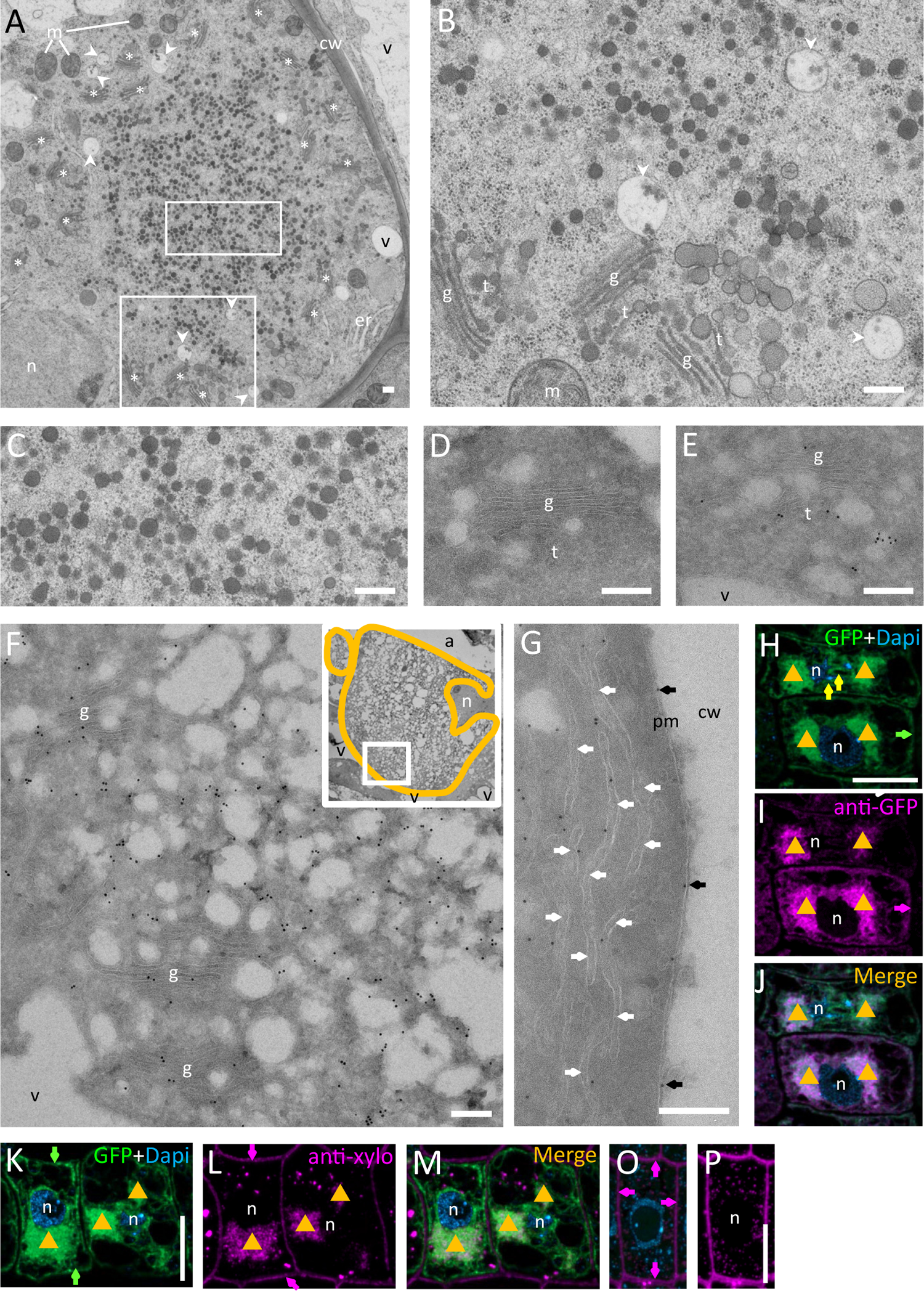
Ultrastructure and immunolocalization analysis of ultrathin transmission electron and light microscopy sections of Estradi-ol-induced GFP-Vamp721 Y57D seedling root tips. (A-C) Ultrastructural analysis of VAMP721Y57D expressing cells after cryofixation and resin embedding. (B, C) Enlarged details of boxed areas in (A). (D-G) Immunogold-labelling of VAMP721Y57D in thawed cryosections. (D) Cells that show no expression of VAMP721Y57D. (E) Cells with low levels of VAMP expression and no aggregate formation. (F-G) Cells with vesicle aggregates: (F) Inset shows the overview of the vesicle aggregate (orange outline) and the selected boxed (white) area enlarged. (G) Heavily labelled cells often show localization of GFP-VAMP721Y57D also at the endoplasmic reticulum and PM. (H-L) Immunolabeling of VAMP721Y57D in thawed 400 nm cryosections. H) GFP fluorescence of GFP-VAMP721Y57D I) immunolabelled GFP J) Merge. ([gl) Vesicle aggregates, ER (yellow arrow) and at the cell periphery (green and margenta arrow) in (H) and (I). K-P) Immunofluorescence labelling of xyloglucan after Estradiol induced expression of GFP-VAMP721Y57D. K-P) GFP fluorescence, xyloglucan labelling and overlay images of cells showing aggregates (K-M) and cells with almost no expression of GFP-VAMP721Y57D (O, overlay; P, enhanced xyloglucan signal). Green and magenta arrows indicate cell periphery. ([gl) vesicle aggregates.) TEM images: a, apoplast; cw, cell wall; er, endoplasmic reticulum; g, Golgi stack; m, mitochondrion; n, nucleus; t, trans-Golgi network; v, vacuole. Asterisk, Golgi stack/TGN; arrow head, multivesicular body; arrow, ER. Magenta: Immunolabeled I,J) GFP, L) L,M,O,P) xyloglucan; Green: H,J,K,,M,O) GFP fluorescence; Blue: H,J,K,M,O) Dapi stained nuclei and mitochondria. Scale bars in A-G = 250 nm, H-Q = 10 µm.

**Supplementary Figure 6:**
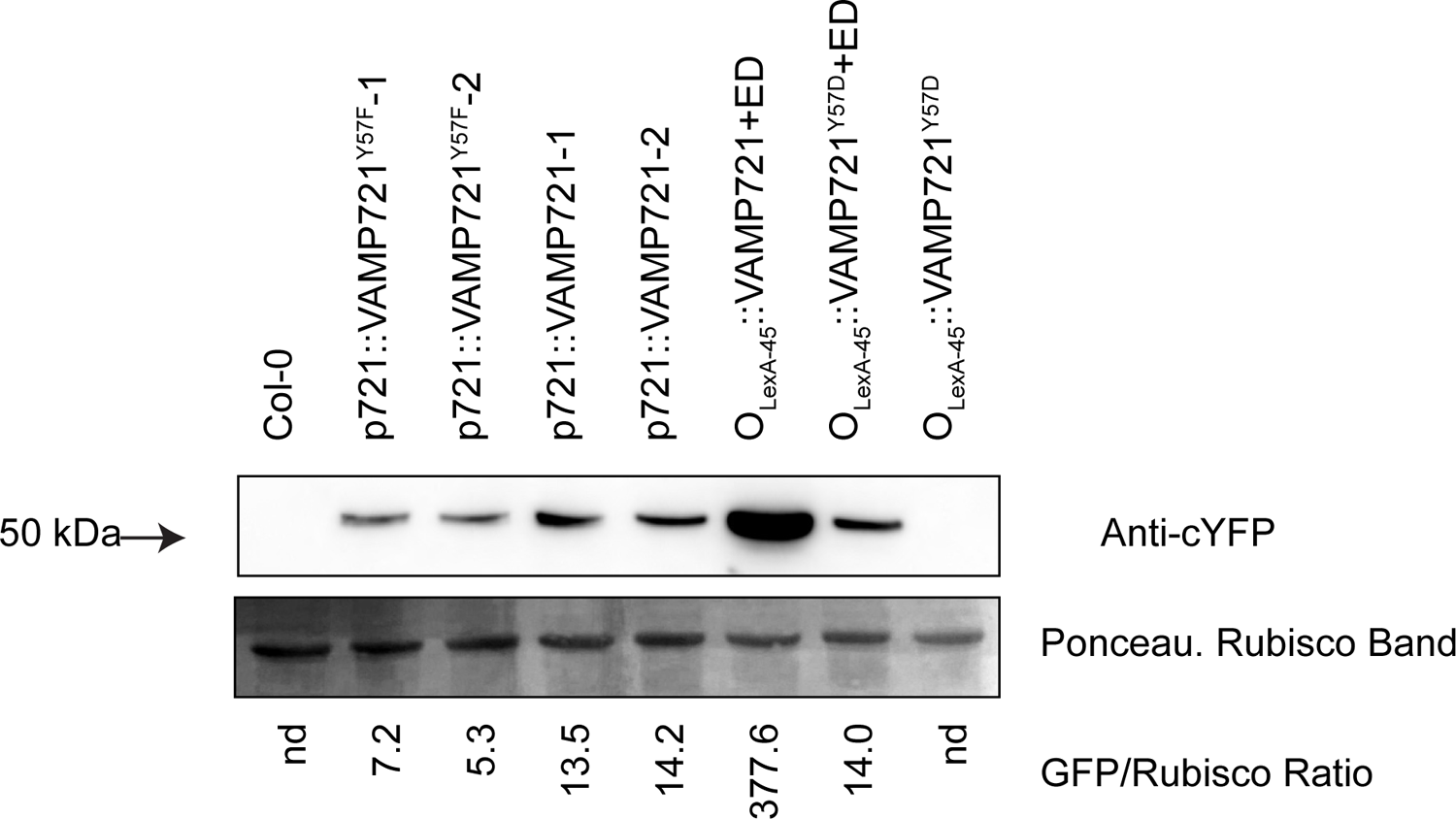
Protein levels in different VAMP721 lines. Anti-cEYFP Immunoblot of pVAMP721 driven GFP-VAMP721 or GFP-VAMP721Y57F and EDI GFP-VAMP721 or GFP-VAMP721Y57D as well as Col-0 and uninduced controls.

**Supplementary Figure 7:**
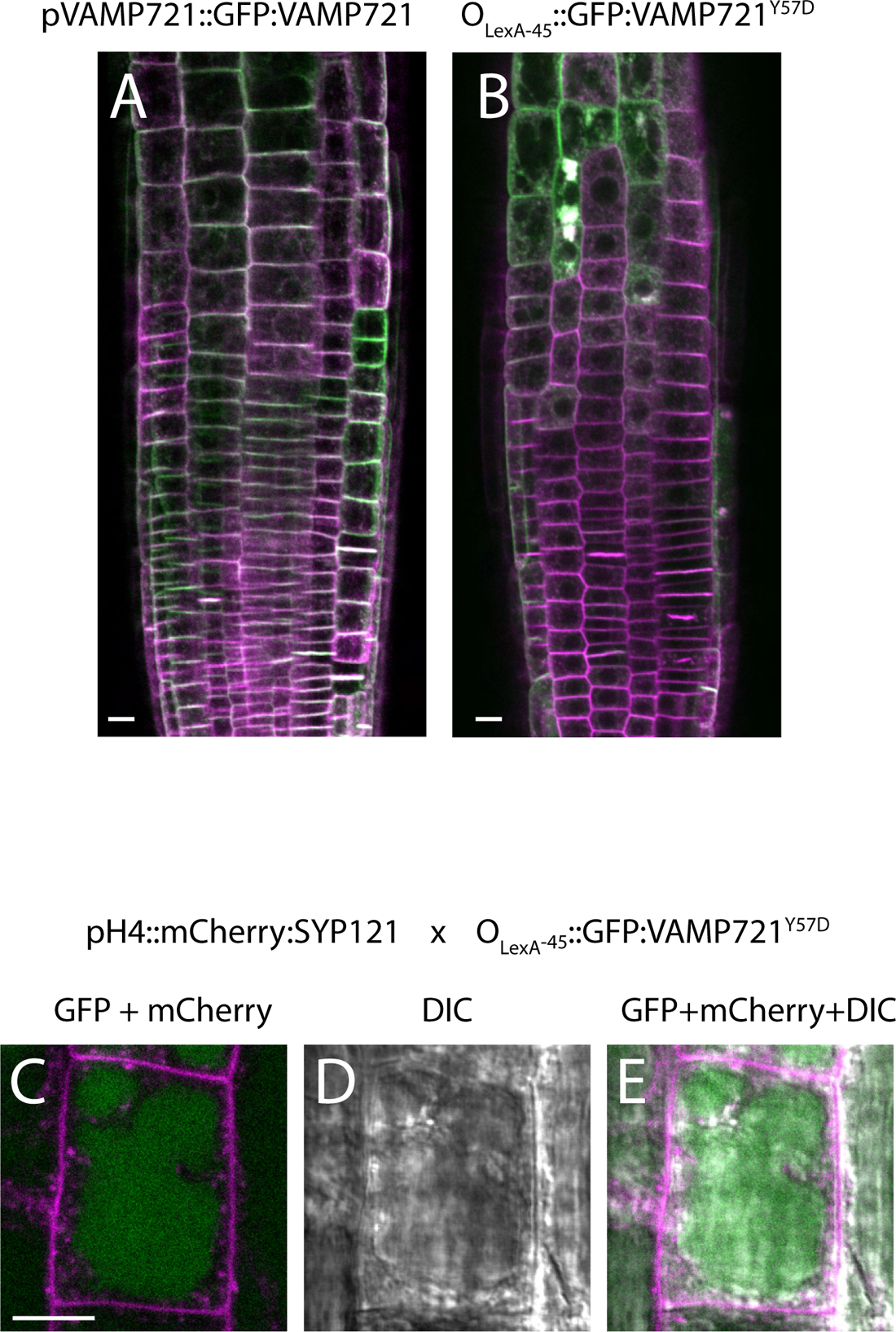
VAMP721 or VAMP721Y57D expression effects on SYP121. Representative confocal images of (A) pVAMP721::GFP:VAMP721 or (B) OlexA-45::GFP:VAMP721Y57D lines crossed with pH4:mcherry:SYP121. (C-E) Individual or merged images showing the lack of mCherry signal inside of the vacuole after dark treatment. GFP signal can readily be seen in (C, E). Scale bar = 10 µm.

**Supplementary Figure 8:**
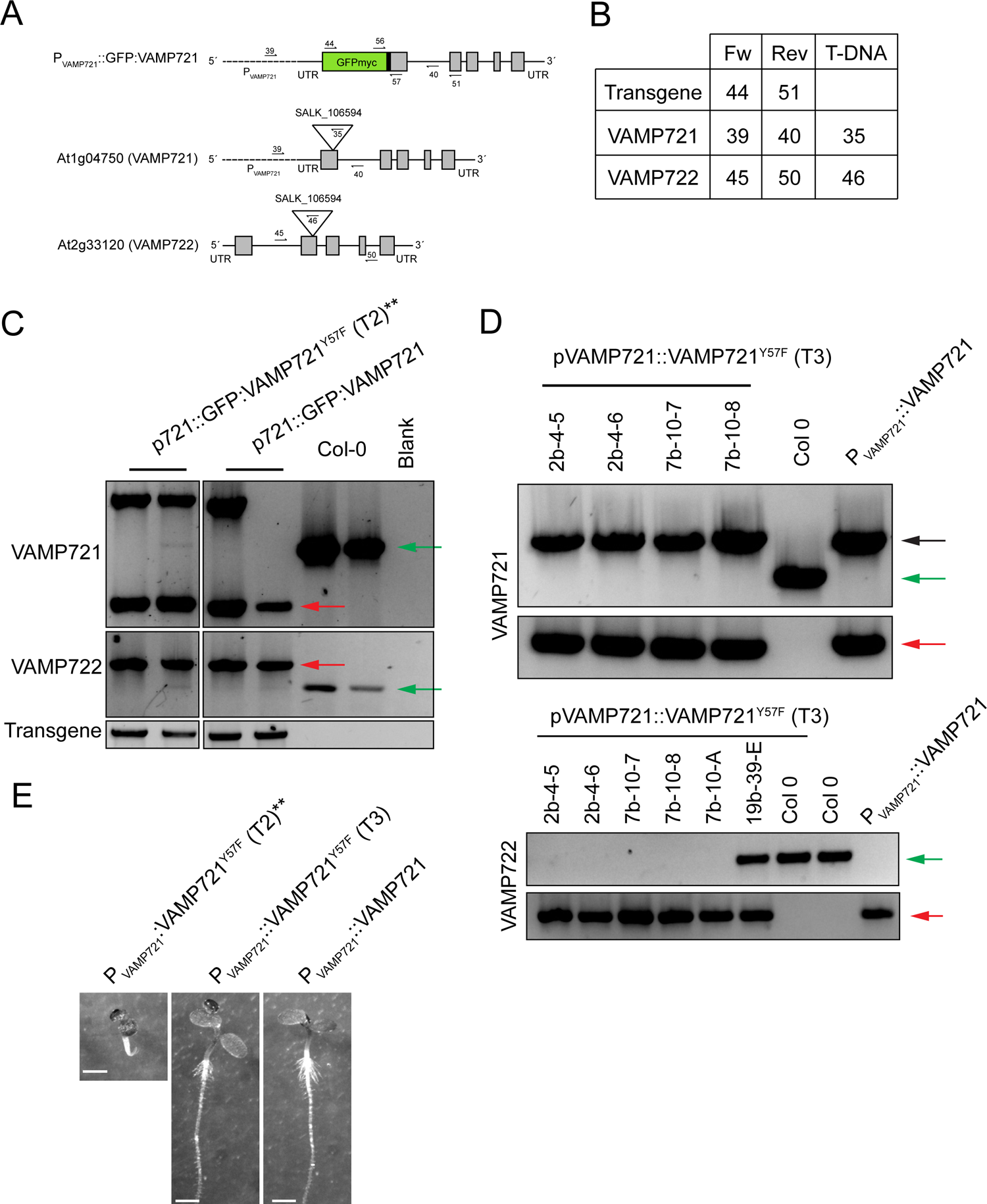
Phenotype rescue in VAMP721Y57F seedlings. (A) Primer binding sites within pVAMP721::GFP:VAMP721 constructs and genomic regions of VAMP721 and VAMP722 endogenous genes. (B) Primer combinations used to amplify transgenes, VAMP721 or VAMP722 regions during genotyping. (C-D) Full genotyping of (C) T2 or (D) T3 lines including Col-0 and VAMP721 rescue controls. Red, green and black arrows indicate the bands for detection of T-DNA, Wt or transgene for VAMP721 or VAMP722 locus as indicated. (E) Phenotype of individuals showing the relevant phenotypes of T2 and T3 lines VAMP721Y57F lines as well as VAMP721 control. ** Indicate mutant seedling phenotype.

