## Supplementary material for "A Tyrosine Phospho-switch within the Longin Domain of VAMP721 modulates SNARE functionality": Suplemental table 2

| Vector name | | Backbone | insert | Cloning method/Cite |
|---|---|---|---|---|
| D606 | pBB-basta-p721::gfp:myc:gVamp721 | V274 | GFP:MyC:gVAMP721 | Restriction-Ligation |
| D607 | pBB-basta-p721::gfp:myc:gVamp721$^{Y57D}$ | V274 | GFP:MyC:gVAMP721$^{Y57D}$ | Restriction-Ligation |
| D608 | pBB-basta-p723::gfp:myc:gVamp723 | V274 | GFP:MyC:gVAMP72e | Restriction-Ligation |
| D610 | pBB-basta-p721::gfp:myc:gVamp723 | V274 | GFP:MyC:gVAMP723 | Restriction-Ligation |
| D932 | pUBC-pMDC7- gfp:myc:gVamp721 | V414 | E443 (GFP:MyC:gVAMP721) | LR recombination |
| D933 | pUBC-pMDC7- gfp:myc:gVamp721$^{Y57D}$ | V414 | E444 (GFP:MyC:gVAMP721$^{Y57D}$) | LR recombination |
| D934 | pUBC-pMDC7- gfp:myc:gVamp723 | V414 | E445 (GFP:MyC:gVAMP723) | LR recombination |
| D1538 | pBB-basta-p723::gfp:myc:gVamp721 | D606 (NheI and XhoI) | D608 (NheI and XhoI) | Restriction-Ligation |
| D1681 | pBB-basta-p721::gfp:myc:gVamp721$^{Y57F}$ | D606 (SnaBI and AfeI) | GFP:MyC:gVAMP721$^{Y57F}$ | Gibson assembly |
| E443 | pDEST207: gfp:myc:gVamp721 | pDEST207 (Invitrogen) | GFP:MyC:gVAMP721 | BP recombination |
| E444 | pDEST207: gfp:myc:gVamp721$^{Y57D}$ | E443 | GFP:MyC:gVAMP721$^{Y57D}$ | SDM |
| E445 | pDEST207: gfp:myc:gVamp723 | pDEST207 (Invitrogen) | GFP:MyC:gVAMP723 | BP recombination |
| V217 | pBiFC-BB | | | Grefen and Blatt 2012 |
| V272 | pUC57-Backbone | | | Gene synthetized |
| V274 | pBBb | V272 (PmeI/EcoRV cut) | V217 (PmeI/SnaBI cut) | Restriction-Ligation |
| V414 | pMDC7-UB-XVE-Hyg | | | https://doi.org/10.7554/eLife.25327 |
