## Supplementary material for "A Tyrosine Phospho-switch within the Longin Domain of VAMP721 modulates SNARE functionality": Suplemental table 1

| Protein names | Gene names | Number of proteins | peptides | MAX iBAQ | VAMP/ Y57D | Max Coverage | Relevant annotated function |
|---|---|---|---|---|---|---|---|
| Novel plant SNARE 12 | NPSN12 | 2 | 7 | 1741045 | #DIV/0 ! | 27.5 | Cytokinesis SNARE |
| Vesicle transport v-SNARE 12 | VTI12 | 1 | 6 | 2748065 | #DIV/0 ! | 25.2 | Secretion / sorting SNARE |
| Syntaxin-61 | SYP61 | 1 | 7 | 4005850 | 20.85 | 40 | Secretion / sorting SNARE |
| Gamma-soluble NSF attachment protein | GSNAP | 1 | 7 | 3024770 | 12.28 | 29.6 | SNARE disassembly |
| Novel plant SNARE 11 | NPSN11 | 1 | 9 | 6730350 | 8.96 | 37 | Cytokinesis SNARE |
| SNAP25 homologous protein SNAP33 | SNAP33 | 1 | 11 | 3661885 | 5.58 | 40.7 | Very broad SNARE |
| SEC1 family transport protein SLY1 | SLY1 | 1 | 11 | 3946400 | 3.68 | 27.9 | Sec/Munc protein |
| Alpha-soluble NSF attachment protein 2 and 1 | ASNAP2; ASNAP1 | 2 | 10 | 3169595 | 3.05 | 38.8 | SNARE disassembly |
| Novel plant SNARE 13 | NPSN13 | 1 | 11 | 12238150 | 2.94 | 63 | Secretory SNARE |
| Aquaporin PIP1-3 | PIP1-3 | 1 | 3 | 2404025 | 2.74 | 10.5 | Aquaporin |
| Syntaxin-81 | SYP81 | 1 | 10 | 5339200 | 2.27 | 39.4 | ER-golgi transport |
| SEC20 | AtSec20 | 1 | 5 | 3971850 | 2.22 | 26.3 | ER-golgi transport |
| Syntaxin-122 | SYP122 | 1 | 7 | 5686150 | 1.82 | 22.3 | Secretory SNARE |
| USE1 | At3g5560 0 | 4 | 8 | 3853400 | 1.77 | 42.1 | ER-golgi transport |
| epsilon 14-3-3-like protein GF14 psi | GRF3 | 2 | 7 | 2569420 | 1.73 | 36.5 | 14-3-3 protein |

| | | | | | | | |
|---|---|---|---|---|---|---|---|
| Syntaxin-132 | SYP132 | 5 | 16 | 37014650 | 1.62 | 47.4 | Secretory SNARE |
| Vesicle transport v-SNARE 11 | VTI11 | 4 | 4 | 1667550 | 1.57 | 18.6 | intracellular transport SNARE |
| Syntaxin-71 | SYP71 | 1 | 20 | 71661500 | 1.54 | 66.5 | ER-golgi transport |
| Syntaxin-121 | SYP121 | 2 | 17 | 40461000 | 1.39 | 46.2 | Secretory SNARE |
| Aquaporin PIP2-1;Aquaporin PIP2-1, N-terminally processed | PIP2-1 | 1 | 5 | 11689750 | 1.38 | 17.4 | Aquaporin |
| Probable aquaporin PIP1-4 | PIP1.4; | 2 | 4 | 1548985 | 1.10 | 18.8 | Aquaporin |
| Pyrophosphate-energized vacuolar membrane proton pump 1 | AVP1 | 2 | 9 | 2669350 | 1.02 | 12.6 | Vacuole |
| Aquaporin PIP1-2 | PIP1B;PIP 1-2 | 3 | 4 | 22872500 | 0.98 | 17.5 | Aquaporin |
| Early response to dehidration / OSCA3.1 | ERD4 | 1 | 12 | 1145225 | 0.97 | 20.4 | Calcium |
| ATPase 2, plasma membrane-type | AHA2;HA 2 | 7 | 29 | 1395280 | 0.90 | 34.3 | PM ATPase |
| epsilon 14-3-3-like protein GF14 mu;14-3-3-like protein GF14 iota | GRF9;GR F12 | 2 | 4 | 1129630 | 0.80 | 14.5 | 14-3-3 protein |
| ABC transporter G family member 36;ABC transporter G family member 35 | ABCG36; ABCG35 | 10 | 23 | 2069750 | 0.79 | 16.8 | ABC transporter |

| | | | | | | | |
|---|---|---|---|---|---|---|---|
| Vesicle-associated membrane protein 721;Vesicle-associated membrane protein 722 | VAMP721;SAR1;VAMP722 | 13 | 12 | 4.24E+09 | 0.75 | 55.3 | Secretory/recycling R-SNARE |
| ATPase 11, plasma membrane-type;ATPase 4, plasma membrane-type | AHA11;AHA4 | 2 | 19 | 2044600 | 0.73 | 21.1 | PM ATPase |
| ATPase 1, plasma membrane-type | AHA1 | 2 | 30 | 19983000 | 0.66 | 33.1 | PM ATPase |
| Aquaporin PIP1-1 | PIP1-1 | 3 | 4 | 4597850 | 0.65 | 16.8 | Aquaporin |
| non-epsilon 14-3-3-like protein GF14 lambda;14-3-3-like protein GF14 kappa | GRF6;GRF8 | 7 | 7 | 4286000 | 0.64 | 32.7 | 14-3-3 protein |
| Aquaporin PIP2-2;Aquaporin PIP2-2, N-terminally processed | PIP2-2 | 2 | 3 | 1542750 | 0.62 | 9.1 | Aquaporin |
| Metacaspase-4;Metacaspase-4 subunit p20;Metacaspase-4 subunit p10 | AMC4 | 2 | 4 | 1373075 | 0.51 | 13.9 | IMMUNITY |
| Aquaporin PIP2-7;Aquaporin PIP2-7, N-terminally processed | PIP2-7 | 1 | 3 | 23481500 | 0.49 | 17.1 | Aquaporin |
| Serine hydroxymethyltransferase 4 | SHM4 | 1 | 10 | 3108650 | 0.47 | 28.9 | METABOLISM |

| | | | | | | | |
|---|---|---|---|---|---|---|---|
| SKP1-like protein 1A and SKP1B | SKP1A; SKP1B | 2 | 3 | 1886070 | 0.43 | 22.5 | Ubiquitination |
| Probable calcium-binding protein CML13 | CML13 | 1 | 6 | 1666250 | 0.41 | 44.6 | Calcium |
| ABC transporter B family member 4 | ABCB4 | 13 | 18 | 2255700 | 0.39 | 18.1 | ABC transporter |
| Annexin D4 | ANN4 | 2 | 4 | 1215930 | 0.38 | 13.9 | Secretion |
| Serine/threonine-protein phosphatase PP2A-1; PP2A-2; PP2A-3; PP2A-5; PERK10 | PP2A5;PP2A3;PP2A2;PP2A1;PERK10 | 10 | 3 | 1048005 | 0.37 | 10.5 | Phosphatase |
| Aldehyde dehydrogenase family 3 member F1 | ALDH3F1 | 1 | 12 | 2164000 | 0.37 | 29.1 | REDOX |
| Clathrin heavy chain 2 | CHC2 | 1 | 19 | 1480535 | 0.35 | 13.2 | CLATRIN |
| | HIT3 | 1 | 3 | 2167050 | 0.34 | 24.5 | METABOLISM |
| Cytochrome b5 isoform E | CYTB5-E | 1 | 3 | 2016950 | 0.33 | 29.9 | TA protein |
| Heat shock protein 90-2 | HSP90-2;HSP81-2 | 2 | 21 | 14987400 | 0.32 | 34 | chaperone |
| V-type proton ATPase subunit D | VHA-D | 1 | 5 | 1561750 | 0.29 | 20.7 | VACUOLE |
| | TLL1 | 3 | 8 | 3924600 | 0.26 | 19.6 | VACUOLE |
| ABC transporter G family member 11 | ABCG11 | 1 | 5 | 1207095 | 0.25 | 6.8 | IMMUNITY |

| | | | | | | | |
|---|---|---|---|---|---|---|---|
| 14-3-3-like protein GF14 epsilon | GRF10 | 3 | 6 | 5430850 | 0.22 | 29.9 | 14-3-3 |
| Trans-cinnamate 4-monooxygenase | CYP73A5 | 6 | 14 | 6087300 | 0.21 | 35.2 | IMMUNITY |
| Ubiquitin-40S ribosomal protein S27a-1,2,3;Ubiquitin;Polyubiquitin 4;Ubiquitin-60S ribosomal protein L40-2;Polyubiquitin 3,10,9 ;Ubiquitin-related 1 to 4;Ubiquitin-60S ribosomal protein L40-1;Ubiquitin;60S ribosomal protein L40-1;Polyubiquitin 14;Ubiquitin;Polyubiquitin 11;Ubiquitin;Polyubiquitin 8;Ubiquitin-related 1-8 | RPS27AA;RPS27AB;RPS27AC;UBQ11;UBQ4;UBQ13;RPL40B;UBQ3;UBQ10;UBQ9;RPL40A;UBQ14;UBQ8 | 18 | 6 | 1371000 | 0.20 | 36.5 | Ubiquitin |
| Tubulin alpha-3 chain;Tubulin alpha-5 chain | TUBA3;TUBA5 | 2 | 13 | 1206650 | 0.20 | 31.3 | Cytoesqueleton |
| ARL8b | K9I9.13 | 1 | 3 | 1533950 | 0.19 | 23.9 | VACUOLE |
| Probable phosphoglucomutase, cytoplasmic 2 | At1g70730 | 3 | 16 | 1220055 | 0.19 | 29 | SUGAR MET |
| Delta(24)-sterol reductase | DIM | 1 | 7 | 2525300 | 0.18 | 13.5 | VACUOLE |
| COP-I Subunit Delta | | 1 | 9 | 2414100 | 0.18 | 23.7 | COATOMER |

| | | | | | | | |
|---|---|---|---|---|---|---|---|
| 26S proteasome non-ATPase regulatory subunit 12 homolog A;26S proteasome non-ATPase regulatory subunit 12 homolog B | EMB2107;RPN5A;RPN5B | 6 | 6 | 1049175 | 0.18 | 13.6 | PROTEOSOME |
| V-type proton ATPase subunit B1;V-type proton ATPase subunit B3;V-type proton ATPase subunit B2 | VHA-B1;VHA-B3;At4g38510;VHA-B2 | 4 | 6 | 1218135 | 0.17 | 20.6 | VACUOLE |
| Patellin-1 | PATL1 | 3 | 12 | 2377150 | 0.16 | 23.6 | VACUOLE |
| V-type proton ATPase subunit H | VHA-H | 1 | 6 | 1603995 | 0.16 | 16.1 | VACUOLE |
| Heat shock 70 kDa protein 9, mitochondrial | HSP70-9 | 1 | 11 | 1288740 | 0.16 | 18.2 | chaperone |
| Eukaryotic initiation factor 4A-2 | TIF4A-2;EIF4A-2 | 2 | 18 | 1037400 | 0.15 | 41.3 | ERAD |
| COP-I Subunit Beta | | 2 | 16 | 2865400 | 0.15 | 21.8 | COATOMER |
| | MO1 | 2 | 8 | 2065000 | 0.14 | 29 | Signaling |
| COP-I Subunit Epsilon | | 1 | 6 | 3554350 | 0.13 | 26.7 | COATOMER |
| F5A18.5 | F5A18.5 | 3 | 7 | 1443675 | 0.13 | 12.8 | IMMUNITY |
| Heat shock protein 90-5 | CR88 | 2 | 18 | 3455750 | 0.12 | 31.9 | chaperone |
| Very-long-chain 3-oxoacyl-CoA reductase 1 | KCR1 | 1 | 8 | 5173900 | 0.12 | 30.5 | OTHER |

| | | | | | | | |
|---|---|---|---|---|---|---|---|
| 6-phosphogluconate dehydrogenase, decarboxylating 3 | | 1 | 18 | 4081200 | 0.12 | 49.6 | SUGAR MET |
| Cyclin-dependent kinase A-1 | CDKA-1 | 1 | 5 | 1140460 | 0.12 | 16.3 | KINASE |
| T-complex protein 1 subunit theta | CCT8 | 1 | 13 | 2069350 | 0.11 | 33.7 | chaperone |
| CLB1 | CLB1/SYT7 | 3 | 5 | 1148595 | 0.10 | 11.6 | ER-PM contact zone |
| MED24.18 | MED24.18 | 1 | 6 | 2389950 | 0.10 | 25.1 | OTHER |
| Probable phosphoglucomutase, cytoplasmic 1 | | 1 | 18 | 4234450 | 0.09 | 41 | SUGAR MET |
| ARM repeat superfamily protein | At3g62530 | 1 | 6 | 5379150 | 0.09 | 30.3 | OTHER |
| 26S protease regulatory subunit 6A homolog A;26S protease regulatory subunit 6A homolog B | RPT5A;RPT5B | 2 | 8 | 3540900 | 0.08 | 27.1 | PROTEOSOME |
| Synaptotagmin-1 | SYTA;SYT1 | 4 | 10 | 1481950 | 0.08 | 18.1 | ER-PM contact zone |
| 26S proteasome regulatory subunit 4 homolog A;26S proteasome regulatory subunit 4 homolog B | RPT2A;RPT2B | 2 | 11 | 5032450 | 0.08 | 33.6 | PROTEOSOME |

| | | | | | | | |
|---|---|---|---|---|---|---|---|
| Tubulin beta-3 chain;Tubulin beta-2 chain | TUBB3;TUBB2 | 2 | 15 | 4692950 | 0.07 | 38.9 | Cytoesqueleton |
| 26S proteasome non-ATPase regulatory subunit 11 homolog | RPN6 | 1 | 5 | 1252960 | 0.07 | 19.3 | PROTEOSOME |
| Mannose-1-phosphate guanylyltransferase 1 | CYT1 | 2 | 5 | 1474500 | 0.07 | 15 | IMMUNITY |
| 26S protease regulatory subunit 7 homolog A | RPT1A | 5 | 6 | 2104900 | 0.06 | 17.4 | PROTEOSOME |
| EF-P | At3g08740 | 1 | 3 | 1352940 | 0.06 | 17.8 | OTHER |
| Aspartate aminotransferase, cytoplasmic isozyme 1 | ASP2 | 4 | 16 | 5530900 | 0.06 | 39.5 | IMMUNITY |
| Chaperone protein dnaJ 3 | ATJ3 | 2 | 10 | 14426500 | 0.05 | 29.8 | Chaperone |
| | CYP706A1;T12H17.100 | 2 | 14 | 5033100 | 0.04 | 28.2 | OTHER |
| Pyrophosphate--fructose 6-phosphate 1-phosphotransferase subunit beta 1 | PFP-BETA1 | 1 | 7 | 1269055 | 0.04 | 18.9 | SUGAR MET |
| 26S proteasome non-ATPase regulatory subunit 4 homolog | RPN10 | 1 | 5 | 3155600 | 0.03 | 16.3 | PROTEOSOME |
| Trifunctional UDP-glucose 4,6-dehydratase/UDP-4-keto-6-deoxy-D-glucose 3,5- | RHM1 | 1 | 11 | 1743000 | 0.03 | 16.9 | CELL WALL |

| | | | | | | | |
|---|---|---|---|---|---|---|---|
| epimerase/UDP-4-keto-L-rhamnose-reductase RHM1;UDP-glucose 4,6-dehydratase;UDP-4-keto-6-deoxy-D-glucose 3,5-epimerase/UDP-4-keto-L-rhamnose 4-keto-reductase | | | | | | | |
| Probable leucine-rich repeat receptor-like serine/threonine-protein kinase At3g14840 | LIK1 | 2 | 11 | 1297430 | 0.03 | 12.3 | IMMUNITY |
| MEC18.18 | T21B14.13;At3g12050 | 2 | 7 | 2387050 | 0.03 | 35.8 | chaperone |
| Hexokinase-1 | HXK1 | 5 | 8 | 1572050 | 0.02 | 24.4 | VACUOLE |
| DnaJ protein ERDJ2A | ERDJ2A | 5 | 11 | 1736600 | 0.02 | 22.1 | chaperone |
| Heat shock 70 kDa protein 2 and 18 | HSP70-2;HSP70-18 | 2 | 31 | 8447400 | 0.00 | 57.3 | chaperone |
| Heat shock 70 kDa protein 4 | HSP70-4 | 1 | 37 | 50057000 | 0.00 | 58.8 | chaperone |
| UNIVERSAL STRESS PROTEIN 17 | F5K20_290;At3g53990 | 2 | 3 | 1222070 | 0.00 | 28.8 | chaperone |
| 26S proteasome non-ATPase regulatory subunit 6 homolog | RPN7 | 1 | 6 | 1677450 | 0.00 | 17.1 | PROTEOSOME |

| | | | | | | | |
|---|---|---|---|---|---|---|---|
| BAG6 | At5g42220 | 1 | 5 | 1326990 | 0.00 | 6.8 | chaperone |
| Chaperone protein dnaJ 2 | ATJ2 | 1 | 9 | 1083855 | 0.00 | 28.2 | chaperone |
| Cytochrome P450 705A5 | CYP705A5 | 12 | 7 | 1353760 | 0.00 | 18.8 | Cytochrome |
| DGR2, DUF642 L-GALL RESPONSIVE GENE 2 | At5g25460 | 2 | 4 | 1687275 | 0.00 | 14.1 | CELL WALL |
| DnaJ protein ERDJ3B | ERDJ3B | 1 | 3 | 1285865 | 0.00 | 11 | chaperone |
| LIM domain-containing protein WLIM1 | WLIM1 | 2 | 3 | 2913900 | 0.00 | 15.3 | Cytoesqueleton |
| Phosphoinositide phosphatase SAC7 | SAC7 | 2 | 8 | 1428920 | 0.00 | 17.6 | PI4P regulation |
| Protein SUPPRESSOR OF K(+) TRANSPORT GROWTH DEFECT 1 | SKD1 | 1 | 7 | 1337500 | 0.00 | 20.7 | VACUOLE |
| | T6H22.10 | 2 | 7 | 1824800 | 0.00 | 16.5 | OTHER |
