## Supplementary material for "A Tyrosine Phospho-switch within the Longin Domain of VAMP721 modulates SNARE functionality": Suplemental methods

**Supplemental Methods**

**Primer Details:**

| | | |
|---|---|---|
| 35 | CG-LBb1.3-SALK | ATTTTGCCGATTTCGGAAC |
| 39 | CG-SALK_106594_LP | CAGGTTCTAAACAATGCAGAGC |
| 40 | CG-SALK_106594_RP | ATCAACCAAAGCTACCACGG |
| 44 | NW-GFP-S | ATGGTGAGCAAGGGCGAG |
| 45 | Nw-F115-Vamp722-3-S | CTTATCAGCGATATATTGATGAATC |
| 46 | Nw-Ma126-LBA1-S | TGGTTCACGTAGTGGGCCATCG |
| 50 | NW-AS-722-geno | gtctctgtgtgggtctattgactttgA |
| 51 | Nw-RT-721-723-AS | AATCAACCGCAACAACACAATAGG |
| 56 | NW-Q-PCR-GFP-S | CATGGTCCTGCTGGAGTTCG |
| 57 | NW-Q-PCR-Myc-Vamp-AS | TGCGCCATGCCTAAGTCC |
| 58 | ACT2_qF | GCCATCCAAGCTGTTCTCTC |
| 59 | ACT2_qR | CAGTAAGGTCACGTCCAGCA |

**Detection of phosphorylation:**

**-Protein digest:** The agarose beads were washed 4 times with ice-cold 100mM NH4HCO3 buffer pH8 (200µl buffer; gentle mixing for 10 sec, centrifugation at 4°C and 4000 rpm). After careful removal of the buffer 10 µl Asp-N endoproteinase solution (0,04 µg/µl in water; Roche, sequencing grade) was added to the beads. After 5 min 20 µl 100 µM NH4HCO3 was layered on top. The subsequent digest was done at 37°C overnight. For improved phosphopeptide sensitivity the resulting 30 µl digest was mixed with 3 µl of 1M citric acid before UPLC-MS analysis.

**-UPLC-MS analysis:** For qualitative and quantitative determination of the phosphorylated and non-phosphorylated peptides, product ion scans (CID) and MRMs were developed using synthetic standards (Genescript). The analysis of the protein digest was performed using an Eksigent Micro-LC 200 and a QTRAP4000 mass spectrometer (ABSciex). Chromatographic separation was achieved at 35°C on a HaloFused $C_{18}$ column (150 x 0.5 mm, particle size 2.7 µm; Eksigent) applying the following binary gradient at a flow rate of 11 µL/min: 0 – 0.5 min isocratic 95% A (water, 0.1% aq. formic acid); 0.5 – 7.5 min, linear from 95% A to 5% A; 7.5 – 10 min, isocratic 95 % B (acetonitrile, 0.1% aq. formic acid). The injection volume was 2 µL. Analytes were ionized using an ESI TurboV ion source equipped with an Assy 65 µm ESI electrode in positive ion mode. The following instrument settings were applied: nebulizer and heater gas, zero grade air, 25 and 10 psi; curtain gas, nitrogen, 20 psi; collision gas, nitrogen, medium; source temperature, 200°C; ionspray voltage, 5000 V; entrance potential, 10 V; collision cell exit potential, 5V; dwell times for the MRMs were set to 200 msec. For the product ion scans a declustering potential (DP) of 60 Volt and a collision energy (CE) of 25 V was applied.

The transitions monitored for each analyte were:

| Q1 mass (Da) | Q3 mass (Da) | DP (volts) | CE (volts) | compound | quantifier ion |
|---|---|---|---|---|---|
| 598.0 | 247.0 | 60 | 25 | non-phospho peptide y2 | + |
| 598.0 | 148.0 | 60 | 25 | non-phospho peptide y1 | - |
| 598.0 | 598.0 | 60 | 10 | non-phospho peptide | - |
| 638.0 | 515.0 | 85 | 25 | phospho peptide b8 | + |
| 638.0 | 148.0 | 85 | 25 | phospho peptide y1 | - |
| 638.0 | 638.0 | 85 | 10 | phospho peptide | - |
| 638.0 | 216.0 | 60 | 55 | phospho peptide Y immonium ion | - |

The LC-MS data acquired were analyzed using the vendors Analyst 1.6.2, MultiQuant 3.0 and PeakView 2.1 software.

**-Results:** The affinity purified GFP fusion protein was cleaved with endoproteinase Asp-N to release the DGHTFNYLVE peptide. Due to very low concentrations and assumed even lower levels of a possibly phosphorylated tyrosine 57 version of the peptide (shot gun proteomics did fail to reveal a positive identification) a targeted LC-MS approach was chosen. Synthetic peptides for both variants were used to create database entries, MRM optimization and calibration curves. Fragments from the doubly charged precursor ions with the highest intensities (transitions: m/z 598/148, y1; m/z 598/247, y2; m/z 638/515, b8; m/z 638/148, y1) as well as the immonium ion of the phospho-tyrosine (transition m/z 638/216) were chosen for reliable quantification. The CID-analysis of the protein digest clearly identified a peak for the DGHTFNYLVE molecule (81% reverse fit, purity 73%). The phosphorylated DGHTFNpYLVE ion could also be seen, albeit in a significantly lower concentration (82% reverse fit, purity 29%). This is also reflected in the comparison of the MRM-XICs of both peptides. As the concentration of the phosphorylated ion was out of the range of the calibration curve (100 pM-10nM) to calculate an exact ratio of DGHTFNYLVE / DGHTFNpYLVE was not possible.

**Immunogold and immunofluorescence localization**

Seedling root tips from the identically treated seedling batch mentioned above were fixed with 4% formaldehyde (30 min) followed by 8% formaldehyde (90 min), embedded in 10% gelatin, frozen on a stub in LN2, and cryosectioned in a Leica UCT7/FC7 cryo-ultramicrotome at -115°C (100 nm sections for TEM) or -80°C (400 nm sections for fluorescence microscopy (FM)). Sections were mounted on pioloform and carbon coated grids (TEM) or coverslips (FM). After blocking unspecific binding sites with 0.5% milk powder and 0.2% gelatin in PBS, sections were incubated with rabbit anti-GFP antibodies (1:500; pabg1, Chromotek, Plannegg-Martinsried), or mouse monoclonal anti-xyloglucan IgG (1:10; mAb CCRC-M1, Carbosource Services, University of Georgia). After washing grids and coverslips 6 times with blocking buffer, sections on grids were incubated with goat anti-rabbit IgG coupled to 6 nm colloidal gold (1:30; Dianova, Hamburg; silver enhanced for 35 min with R-Gent, Aurion, Wageningen), sections on coverslips with goat anti-rabbit IgG coupled to Cy3 (1:400; Dianova, Hamburg). After washing the sections with blocking buffer (3x) and PBS (3x), grids were washed with bidistilled water and embedded in a thin layer of methyl cellulose, whereas coverslips were stained with Dapi (4´,6-diamidino-2-phenylindole; 1 µg/ml; Sigma) and embedded in Mowiol 4.88 (Calbiochem) supplemented with DABCO (1,4-diazabicyclo[2.2.2]octane; 25 mg/ml; anti-fading agent; Sigma). Coverslips were viewed in a Zeiss Axioimager M2 (63x/1.4 oil immersion objective) using filter sets for Cy3, GFP, and Dapi fluorescence. Some images were taken with an Apotome 1 attachment, deconvolution was set to 1 or 2. Images were taken with a sCMOS Orca-flash4.0 camera (Hamamatsu). Image contrast and brightness was adapted using Zeiss ZEN Blue and Adobe Photoshop CS5 software. Controls used: As GFP expressing seedling root tips also contain a large number of cells without showing GFP fluorescence and cells weakly expressing GFP (mainly in the meristematic zone), these parts of the root tip were used as an internal control (for TEM see Supplemental Figure 5D, E; for LM see Supplemental Figure 5O, P). The GFP-specific antibody turned out to be highly specific as FM shows nearly complete colocalization of GFP fluorescence and anti-GFP immunofluorescence (for LM see Supplemental Figure 5H-J). Xyloglucan labelling resulted in the well-known and often published labelling patterns (Zhang and Staehelin, 1992; Stierhof and El Kasmi, 2010).

```
FEATURES             Location/Qualifiers
     misc_feature    join(12036..12189,1..42)
                     /vntifkey="21"
                     /label="RB"
                     /ApEinfo_fwdcolor="pink"
                     /ApEinfo_revcolor="pink"
                     /ApEinfo_graphicformat="arrow_data {{0 1 2 0 0 -1} {}
                     0}width 5 offset 0"
     CDS             68..1400
                     /label="pVAMP721"
     misc_feature    68..1400
                     /label="Promotor Vamp721"
                     /ApEinfo_fwdcolor="cyan"
                     /ApEinfo_revcolor="green"
                     /ApEinfo_graphicformat="arrow_data {{0 1 2 0 0 -1} {}
                     0}width 5 offset 0"
     Site            join(745..754,749^750)
                     /site_type="restriction site"
                     /note="Name: BsaBI"
                     /note="Pattern: gatnnnnatc"
                     /note="Number_of_matches: 5"
                     /note="cut_0_on_positive_strand: 749^750"
                     /note="cut_0_on_negative_strand: 749^750"
                     /note="inhibited_by: "
                     /note="site_type: other"
                     /note="restriction site"
                     /label="BsaBI"
     5'UTR           1176..1400
                     /label="VAMP721 UTR"
     CDS             1407..1409
                     /label="MYC"
                     /ApEinfo_fwdcolor="pink"
                     /ApEinfo_revcolor="pink"
                     /ApEinfo_graphicformat="arrow_data {{0 1 2 0 0 -1} {}
                     0}width 5 offset 0"
     misc_feature    1410..2126
                     /label="XhoI-meGFP-SalI"
                     /ApEinfo_fwdcolor="#ff8000"
                     /ApEinfo_revcolor="#400040"
                     /ApEinfo_graphicformat="arrow_data {{0 1 2 0 0 -1} {}
                     0}width 5 offset 0"
     misc_feature    1672..1689
                     /label="RT-fwd"
                     /ApEinfo_fwdcolor="cyan"
                     /ApEinfo_revcolor="green"
                     /ApEinfo_graphicformat="arrow_data {{0 1 2 0 0 -1} {}
                     0}width 5 offset 0"
     CDS             2130..2159
                     /label="MYC(1)"
                     /ApEinfo_label="MYC"
                     /ApEinfo_fwdcolor="pink"
                     /ApEinfo_revcolor="pink"
                     /ApEinfo_graphicformat="arrow_data {{0 1 2 0 0 -1} {}
```

0}width 5 offset 0"
misc_feature 2163..4123
/label="gVamp721"
/ApEinfo_fwdcolor="cyan"
/ApEinfo_revcolor="green"
/ApEinfo_graphicformat="arrow_data {{0 1 2 0 0 -1} {}
0}width 5 offset 0"
mutation 2331..2333
/label="Y57F Change"
misc_feature 2859..2882
/label="RT-rev-1"
/ApEinfo_fwdcolor="#0000a0"
/ApEinfo_revcolor="green"
/ApEinfo_graphicformat="arrow_data {{0 1 2 0 0 -1} {}
0}width 5 offset 0"
misc_feature 3390..3409
/label="g721 seq primer"
/ApEinfo_fwdcolor="#d70428"
/ApEinfo_revcolor="green"
/ApEinfo_graphicformat="arrow_data {{0 1 2 0 0 -1} {}
0}width 5 offset 0"
misc_feature 3661..3723
/label="TMD"
/ApEinfo_fwdcolor="#ffff00"
/ApEinfo_revcolor="green"
/ApEinfo_graphicformat="arrow_data {{0 1 2 0 0 -1} {}
0}width 5 offset 0"
misc_feature 3686..3709
/label="RT-rev-2"
/ApEinfo_fwdcolor="#0000a0"
/ApEinfo_revcolor="green"
/ApEinfo_graphicformat="arrow_data {{0 1 2 0 0 -1} {}
0}width 5 offset 0"
Site join(3687..3696,3691^3692)
/site_type="restriction site"
/note="Name: BsaBI"
/note="Pattern: gatnnnnatc"
/note="Number_of_matches: 5"
/note="cut_0_on_positive_strand: 3691^3692"
/note="cut_0_on_negative_strand: 3691^3692"
/note="inhibited_by: "
/note="site_type: other"
/note="restriction site"
/label="BsaBI"
3'UTR 3733..4123
/label="VAMP721 UTR"
terminator 4149..4354
/label="T35S"
/ApEinfo_fwdcolor="pink"
/ApEinfo_revcolor="pink"
/ApEinfo_graphicformat="arrow_data {{0 1 2 0 0 -1} {}
0}width 5 offset 0"
promoter 4358..4660

/vntifkey="29"
/label="nosP"
/ApEinfo_fwdcolor="pink"
/ApEinfo_revcolor="pink"
/ApEinfo_graphicformat="arrow_data {{0 1 2 0 0 -1} {}
0}width 5 offset 0"
misc_feature 4360..4365
/label="BspHI site"
/ApEinfo_fwdcolor="cyan"
/ApEinfo_revcolor="green"
/ApEinfo_graphicformat="arrow_data {{0 1 2 0 0 -1} {}
0}width 5 offset 0"
CDS 4661..5212
/vntifkey="4"
/label="BarCO"
/ApEinfo_fwdcolor="pink"
/ApEinfo_revcolor="pink"
/ApEinfo_graphicformat="arrow_data {{0 1 2 0 0 -1} {}
0}width 5 offset 0"
Site join(5013..5022,5017^5018)
/site_type="restriction site"
/note="Name: BsaBI"
/note="Pattern: gatnnnnatc"
/note="Number_of_matches: 5"
/note="cut_0_on_positive_strand: 5036^5037"
/note="cut_0_on_negative_strand: 5036^5037"
/note="inhibited_by: "
/note="site_type: other"
/note="restriction site"
/label="BsaBI"
terminator 5213..5501
/vntifkey="43"
/label="nosT"
/ApEinfo_fwdcolor="pink"
/ApEinfo_revcolor="pink"
/ApEinfo_graphicformat="arrow_data {{0 1 2 0 0 -1} {}
0}width 5 offset 0"
Site join(5291..5300,5295^5296)
/site_type="restriction site"
/note="Name: BsaBI"
/note="Pattern: gatnnnnatc"
/note="Number_of_matches: 5"
/note="cut_0_on_positive_strand: 5314^5315"
/note="cut_0_on_negative_strand: 5314^5315"
/note="inhibited_by: "
/note="site_type: other"
/note="restriction site"
/label="BsaBI"
misc_feature complement(5522..5851)
/vntifkey="21"
/label="LB"
/ApEinfo_fwdcolor="pink"
/ApEinfo_revcolor="pink"

```
                     /ApEinfo_graphicformat="arrow_data {{0 1 2 0 0 -1} {}
                     0}width 5 offset 0"
     CDS             5857..7101
                     /vntifkey="4"
                     /label="SmR"
                     /ApEinfo_fwdcolor="pink"
                     /ApEinfo_revcolor="pink"
                     /ApEinfo_graphicformat="arrow_data {{0 1 2 0 0 -1} {}
                     0}width 5 offset 0"
     rep_origin      7375..8055
                     /vntifkey="33"
                     /label="pBR322 ori"
                     /ApEinfo_fwdcolor="pink"
                     /ApEinfo_revcolor="pink"
                     /ApEinfo_graphicformat="arrow_data {{0 1 2 0 0 -1} {}
                     0}width 5 offset 0"
     rep_origin      8465..11059
                     /vntifkey="33"
                     /label="pVS1 ori"
                     /ApEinfo_fwdcolor="pink"
                     /ApEinfo_revcolor="pink"
                     /ApEinfo_graphicformat="arrow_data {{0 1 2 0 0 -1} {}
                     0}width 5 offset 0"
     Site            join(10703..10712,10707^10708)
                     /site_type="restriction site"
                     /note="Name: BsaBI"
                     /note="Pattern: gatnnnnatc"
                     /note="Number_of_matches: 5"
                     /note="cut_0_on_positive_strand: 10726^10727"
                     /note="cut_0_on_negative_strand: 10726^10727"
                     /note="inhibited_by: "
                     /note="site_type: other"
                     /note="restriction site"
                     /label="BsaBI"
ORIGIN
        1 AAACTGAAGG CGGGAAACGA CAATCTGATC CAAGCTCAAG CTAAGCTAGG ACTAGCGCTC
       61 GCACGTGTCT TCCTTTTATC CAGATGAGAG AAAGATAAAA AAACGAAACC TAAGAACACC
      121 ACAAAAATCC GATCGAATTG ATGAGAGAGA TTAGTAAGCG ATGTAAAAAT ACACAAGGGA
      181 ACGAGGGGTA GCGATGGAGA AAGCAGGACG CAGAGATCGG AGAAGAGAAA CTATATGAGA
      241 GCGCTGGATT TGCGCGAGTA ACAGAGACAT TATTTTCCCT TTGGTAACGT TCAAATTTTG
      301 AATTTTTAAA TATAAAAAGT AAATAAAAAG AAAAATTCAT CATTTGTATA GTCGAAAATT
      361 GCCTTAATAA ACTTTATTTT GGTAAATAAT TCACGTCTAA TTAATAATAG TATCTTCATG
      421 TTTTTACACC ACGGTGGCTC GTTTTTCAAT TCAAATGTGA AATGTTTATC AATGAGCATT
      481 TTAAAAAACA AATATATATA TATATATATA TATTATATAT TGATTTTTTA TAAAGAAAAT
      541 TGCGATGTGT AAGGGATATT TGAGTTGTTA AATTTTCTAT AAATTATATG TTATGTTACT
      601 TAAAAGAACG TTCAATTGAT AATGTTGCAA ACCTTCCTAC CAAAAAAATT AAAATAATGT
      661 TGCAAACCTT TTCTTTCATT CCAATTCACT AACCTTCTAT TCTAACCACA ACCTAACTAC
      721 CTAAATTTCA ATTATTTGAT TGTTGATCAA AATCTTTAGG AACACGTTAA ACAAATTAAT
      781 TAGCAGGTTC TAAACAATGC AGAGCAATTG TAGATGACAA GAGATAACGG TAGAAGTTCG
      841 AATTTTGTTG CTGTAACAAG ATACTATGAC TATGAGCTCC CCCCGTCCAT TAAGAATTAA
      901 GAAATTTGGT TAATTACAAC ATTTGTTTTT ATGTTTAAGC CTTTTGTTCG TAACTGAGCT
      961 AATACTGGAA TCACTGATCA ATGATGAATT TGGAAAAAAA GATATTTTGA GGAAGACAGA
     1021 GACAGACTCT CTTCGACCAT GGGACCTGAA CTTGTCCTAA AGTTGGTAAT TACTAATTTG
```

1081 TGCCATCGGC AGAGTCCACG AGCTTTTTGG CTGGATCCGG GCGGGTCCAA CTTTCTCGAC
1141 ATTTGTTTGA TCTCTCCGTT TTGAAATTGT TTTTTTTTAG TCCTTAAATT CAAGGTTTTA
1201 TACTATTTTT GACCCAATCT GACTACCTTA AATGACAACT TTACCCTTGC CCGCTTTTCA
1261 ACTTTTTCTT CTCCCATGTC CACTTTAGTC CTCGTTTTGT TCTTTCATAG TCTTCTAGAT
1321 CTGTAGTTTT TTTCCGCGAC CGCCCAAAAA ATTCTCTGAT CAGACGGTCG ATTGGATCGG
1381 AGATTTAAGG TAAAGAAAAA ATGCTCGAGA TGGTGAGCAA GGGCGAGGAG CTGTTCACCG
1441 GGGTGGTGCC CATCCTGGTC GAGCTGGACG GCGACGTAAA CGGCCACAAG TTCAGCGTGT
1501 CCGGCGAGGG CGAGGGCGAT GCCACCTACG GCAAGCTGAC CCTGAAGTTC ATCTGCACCA
1561 CCGGCAAGCT GCCCGTGCCC TGGCCCACCC TCGTGACCAC CCTGACCTAC GGCGTGCAGT
1621 GCTTCAGCCG CTACCCCGAC CACATGAAGC AGCACGACTT CTTCAAGTCC GCCATGCCCG
1681 AAGGCTACGT CCAGGAGCGC ACCATCTTCT TCAAGGACGA CGGCAACTAC AAGACCCGCG
1741 CCGAGGTGAA GTTCGAGGGC GACACCCTGG TGAACCGCAT CGAGCTGAAG GGCATCGACT
1801 TCAAGGAGGA CGGCAACATC CTGGGGCACA AGCTGGAGTA CAACTACAAC AGCCACAACG
1861 TCTATATCAT GGCCGACAAG CAGAAGAACG GCATCAAGGT GAACTTCAAG ATCCGCCACA
1921 ACATCGAGGA CGGCAGCGTG CAGCTCGCCG ACCACTACCA GCAGAACACC CCCATCGGCG
1981 ACGGCCCCGT GCTGCTGCCC GACAACCACT ACCTGAGTAC TCAGTCCAAG CTGAGCAAAG
2041 ACCCCAACGA GAAGCGCGAT CACATGGTCC TGCTGGAGTT CGTGACCGCC GCCGGGATCA
2101 CTCTCGGCAT GGACGAGCTG TACAAGGTCG AGCAGAAGTT GATCTCAGAG GAGGACTTAG
2161 GCATGGCGCA ACAATCGTTG ATCTACAGTT TCGTAGCTCG CGGCACGGTG ATCCTCGTTG
2221 AGTTCACTGA TTTCAAAGGT AATTTCACCT CAATCGCTGC TCAGTGCCTC CAGAAGCTTC
2281 CGTCTTCGAA CAACAAGTTC ACCTACAACT GCGATGGCCA TACCTTCAAT TTCCTTGTCG
2341 AAGATGGATT CAGTAAGTCA CTTTTCGTTT GATCTATGCA TAGGTTTTAA TCTTACACCA
2401 TTCGGTGCTT ACCTCGATTT GTTTCTAGGT TTATTTGCCT AACCTATCGA TTCGTATGGA
2461 TTTGTGCATC CTGAGTAGTC TGATTTCACT GAGGAACCGC ATGAACCAAG CTTTTAGGGT
2521 TAGATCTATA TTTATTAAAC GCAATAAATC TCTGTAGAGT CTTACCTGTA TAATCACTCA
2581 GATTTGGACA GAATCCGTGG TAGCTTTGGT TGATACTATG TGGAAGAACA CCAACATTGA
2641 TCTGAGACAG GGTTTTTACT GGTTGCTTTA TTTGCGTATA GTTTCCTAGG TCATGGGTTT
2701 TGGCTGGCTT CATTTTTCAA TGAAAATCTG GGGTCTTATT GTAAGCTTTT GTGACTATCC
2761 TTTAGTTGTT GTAGCCATAT CAATTTCATA TTAAATCAAT GTCACCTGAT TATCCCGTAA
2821 GGGAAGACAA GGAGATTGAT TTTGTTCCTG TTGTACAGCC TATTGTGTTG TTGCGGTTGA
2881 TTCTGCTGGG AGGCAAATTC CCATGTCCTT TTTGGAAAGA GTAAAAGAAG ATTTTAACAA
2941 GCGATATGGT GGTGGAAAGG CTGCAACTGC TCAAGCAAAC AGCTTGAATA AGGAGTTTGG
3001 GTACTTTTTC ATTATCTCTT CTATTTGGAT GATCTTCTCT TTTATATTCG TGGACTGACT
3061 CTTTTTGGTA ACTTTGAAGC TCTAAACTGA AAGAGCATAT GCAGTATTGC ATGGATCATC
3121 CTGATGAGAT TAGCAAGCTT GCTAAGGTGA AGGCGCAAGT GTCAGAAGTT AAGGGTGTAA
3181 TGATGGAAAA CATTGAGAAG GTTTGAATCT GACCCTTTCT GTCCTCAATG TATATATTTA
3241 CATCTATGGT TGACCCATCT GAAAGAGCTC ATCTAGTCAT ACTAAGTTAC TGTGAAATCA
3301 ATTACTAATA ATTCAATGCA TTACATCTTA CGGGAAATGT CCAGTTTACC AATGTAACAC
3361 CGCTTACATA TGCGACTCAT TATTTGCAGG TTCTTGACCG TGGTGAGAAA ATTGAGCTTT
3421 TGGTGGACAA AACCGAAAAT CTTCGCTCAC AGGTTAGACT CTCTCATACC CTTATCTCCT
3481 GCATCTATTT GCCTACACTC ACACACAAAG ATGAACATGC TTTTTGAGTT CATAGCTGAC
3541 TCATCCTATT ACCATATCTT GTGTCTAACT GAACTCAAAC TGCCACAGGC ACAAGATTTC
3601 AGAACAACAG GAACGCAGAT GAGAAGAAAG ATGTGGCTTC AGAACATGAA GATAAAACTC
3661 ATAGTGCTCG CCATCATTAT CGCACTGATT CTCATCATCG TGCTCTCAGT TTGCCATGGG
3721 TTTAAGTGTT AAGCCCAGAA AATTCAAAAC TTCTGAGATC TTTCTCACTC GGTACATCGT
3781 AGTGCCTTCT CTTTTATTGT TCGATCTTCA ATCTAATGTT TCTGCTTGAA TTGGTTTTGT
3841 ATAGATAGAC ATATATGTAT GCTACTCATT TGCTATATTG CTTGCATCTG ATGACATTGA
3901 TCCCTTGCGA ACCAGTCAGT ACTTGACAAG AGATCCGAAA CTATTAAATG ATTTTGTTGA
3961 CTGAAGATTG CATTGTGGTG GTGATGATAA AAATTTGAGA TTGTATTGAC GATTATACGG
4021 CGTGATGTGT GCTGGTTACT TTAAATTTAG ATAAGAGTAG TGCCCTCTCT ATATCGTTAC
4081 TAGGGATGGT AGTACATATA TATACGTACG TCATATTCCG ACCGTTTTAA GCGGCCATGC
4141 TAGAGTCCGC AAAAATCACC AGTCTCTCTC TACAAATCTA TCTCTCTCTA TTTTTCTCCA
4201 GAATAATGTG TGAGTAGTTC CCAGATAAGG GAATTAGGGT TCTTATAGGG TTTCGCTCAT

4261 GTGTTGAGCA TATAAGAAAC CCTTAGTATG TATTTGTATT TGTAAAATAC TTCTATCAAT
4321 AAAATTTCTA ATTCCTAAAA CCAAAATCCA GTGAGTAGAT CATGAGCGGA GAATTAAGGG
4381 AGTCACGTTA TGACCCCCGC CGATGACGCG GGACAAGCCG TTTTACGTTT GGAACTGACA
4441 GAACCGCAAC GTTGAAGGAG CCACTGAGCG CGGGTTTCTG GAGTTTAATG AGCTAAGCAC
4501 ATACGTCAGA AACCATTATT GCGCGTTCAA AAGTCGCCTA AGGTCACTAT CATCTAGCAA
4561 ATATTTCTTG TCAAAAATGC TCCACTGACG TTCCATAAAT TCCCCTCGGT ATCCAATTAG
4621 AGTCTCATAT TCACTCTCAA CTCGATCGAG GGGATCTACC ATGAGTCCAG AAAGGAGACC
4681 AGCAGATATT AGGAGAGCAA CCGAAGCAGA TATGCCAGCA GTTTGCACCA TTGTGAACCA
4741 TTACATCGAG ACTTCTACAG TTAATTTTAG GACTGAACCT CAAGAGCCAC AGGAATGGAC
4801 AGATGATCTC GTGAGATTAA GGGAAAGATA CCCTTGGCTT GTTGCTGAGG TGGATGGAGA
4861 AGTTGCAGGT ATTGCTTACG CAGGACCTTG GAAGGCTAGA AACGCTTATG ATTGGACTGC
4921 TGAGTCTACC GTTTACGTGT CACCTAGACA TCAAAGAACC GGACTTGGTT CAACCTTGTA
4981 TACTCACCTT TTGAAGTCTC TTGAAGCTCA GGGATTCAAA TCTGTTGTGG CTGTTATCGG
5041 TTTGCCTAAT GATCCAAGTG TGAGAATGCA TGAAGCTCTC GGATACGCAC CAAGGGGTAT
5101 GTTAAGAGCT GCTGGATTTA AACATGGTAA CTGGCACGAT GTTGGTTTCT GGCAGTTAGA
5161 TTTCAGTTTA CCAGTTCCAC CAAGACCAGT GCTTCCAGTG ACCGAGATTT GACCCGGGCT
5221 AGAGTCAAGC AGATCGTTCA AACATTTGGC AATAAAGTTT CTTAATATTG AATCCTGTTG
5281 CCGGTCTTGC GATGATTATC ATATAATTTC TGTTGAATTA CGTTAAGCAT GTAATAATTA
5341 ACATGTAATG CATGACGTTA TTTATGAGAT GGGTTTTTAT GATTAGAGTC CCGCAATTAT
5401 ACATTTAATA CGCGATAGAA AACAAAATAT AGCGCGCAAA CTAGGATAAA TTATCGCGCG
5461 CGGTGTCATC TATGTTACTA GATCGACCGG CATGAAGCTG AGTTAACGAT GTAAGTCCTC
5521 AATTCGGCGT TAATTCAGTA CATTAAAAAC GTCCGCAATG TGTTATTAAG TTGTCTAAGC
5581 GTCAATTTGT TTACACCACA ATATATCCTG CCACCAGCCA GCCAACAGCT CCCCGACCGG
5641 CAGCTCGGCA CAAAATCACC ACTCGATACA GGCAGCCCAT CAGTCCGGGA CGGCGTCAGC
5701 GGGAGAGCCG TTGTAAGGCG GCAGACTTTG CTCATGTTAC CGATGCTATT CGGAAGAACG
5761 GCAACTAAGC TGCCGGGTTT GAAACACGGA TGATCTCGCG GAGGGTAGCA TGTTGATTGT
5821 AACGATGACA GAGCGTTGCT GCCTGTGATC AATTCGGGCA CGAACCCAGT GGACATAAGC
5881 CTGTTCGGTT CGTAAGCTGT AATGCAAGTA GCGTATGCGC TCACGCAACT GGTCCAGAAC
5941 CTTGACCGAA CGCAGCGGTG GTAACGGCGC AGTGGCGGTT TTCATGGCTT GTTATGACTG
6001 TTTTTTTGGG GTACAGTCTA TGCCTCGGGC ATCCAAGCAG CAAGCGCGTT ACGCCGTGGG
6061 TCGATGTTTG ATGTTATGGA GCAGCAACGA TGTTACGCAG CAGGGCAGTC GCCCTAAAAC
6121 AAAGTTAAAC ATCATGGGGG AAGCGGTGAT CGCCGAAGTA TCGACTCAAC TATCAGAGGT
6181 AGTTGGCGTC ATCGAGCGCC ATCTCGAACC GACGTTGCTG GCCGTACATT TGTACGGCTC
6241 CGCAGTGGAT GGCGGCCTGA AGCCACACAG TGATATTGAT TTGCTGGTTA CGGTGACCGT
6301 AAGGCTTGAT GAAACAACGC GGCGAGCTTT GATCAACGAC CTTTTGGAAA CTTCGGCTTC
6361 CCCTGGAGAG AGCGAGATTC TCCGCGCTGT AGAAGTCACC ATTGTTGTGC ACGACGACAT
6421 CATTCCGTGG CGTTATCCAG CTAAGCGCGA ACTGCAATTT GGAGAATGGC AGCGCAATGA
6481 CATTCTTGCA GGTATCTTCG AGCCAGCCAC GATCGACATT GATCTGGCTA TCTTGCTGAC
6541 AAAAGCAAGA GAACATAGCG TTGCCTTGGT AGGTCCAGCG GCGGAGGAAC TCTTTGATCC
6601 GGTTCCTGAA CAGGATCTAT TTGAGGCGCT AAATGAAACC TTAACGCTAT GGAACTCGCC
6661 GCCCGACTGG GCTGGCGATG AGCGAAATGT AGTGCTTACG TTGTCCCGCA TTTGGTACAG
6721 CGCAGTAACC GGCAAAATCG CGCCGAAGGA TGTCGCTGCC GACTGGGCAA TGGAGCGCCT
6781 GCCGGCCCAG TATCAGCCCG TCATACTTGA AGCTAGACAG GCTTATCTTG GACAAGAAGA
6841 AGATCGCTTG GCCTCGCGCG CAGATCAGTT GGAAGAATTT GTCCACTACG TGAAAGGCGA
6901 GATCACCAAG GTAGTCGGCA AATAATGTCT AGCTAGAAAT TCGTTCAAGC CGACGCCGCT
6961 TCGCGGCGCG GCTTAACTCA AGCGTTAGAT GCACTAAGCA CATAATTGCT CACAGCCAAA
7021 CTATCAGGTC AAGTCTGCTT TTATTATTTT TAAGCGTGCA TAATAAGCCC TACACAAATT
7081 GGGAGATATA TCATGCATGA CCAAAATCCC TTAACGTGAG TTTTCGTTCC ACTGAGCGTC
7141 AGACCCCGTA GAAAAGATCA AAGGATCTTC TTGAGATCCT TTTTTTCTGC GCGTAATCTG
7201 CTGCTTGCAA ACAAAAAAAC CACCGCTACC AGCGGTGGTT TGTTTGCCGG ATCAAGAGCT
7261 ACCAACTCTT TTTCCGAAGG TAACTGGCTT CAGCAGAGCG CAGATACCAA ATACTGTCCT
7321 TCTAGTGTAG CCGTAGTTAG GCCACCACTT CAAGAACTCT GTAGCACCGC CTACATACCT
7381 CGCTCTGCTA ATCCTGTTAC CAGTGGCTGC TGCCAGTGGC GATAAGTCGT GTCTTACCGG

7441 GTTGGACTCA AGACGATAGT TACCGGATAA GGCGCAGCGG TCGGGCTGAA CGGGGGGTTC
7501 GTGCACACAG CCCAGCTTGG AGCGAACGAC CTACACCGAA CTGAGATACC TACAGCGTGA
7561 GCTATGAGAA AGCGCCACGC TTCCCGAAGG GAGAAAGGCG GACAGGTATC CGGTAAGCGG
7621 CAGGGTCGGA ACAGGAGAGC GCACGAGGGA GCTTCCAGGG GGAAACGCCT GGTATCTTTA
7681 TAGTCCTGTC GGGTTTCGCC ACCTCTGACT TGAGCGTCGA TTTTTGTGAT GCTCGTCAGG
7741 GGGGCGGAGC CTATGGAAAA ACGCCAGCAA CGCGGCCTTT TTACGGTTCC TGGCCTTTTG
7801 CTGGCCTTTT GCTCACATGT TCTTTCCTGC GTTATCCCCT GATTCTGTGG ATAACCGTAT
7861 TACCGCCTTT GAGTGAGCTG ATACCGCTCG CCGCAGCCGA ACGACCGAGC GCAGCGAGTC
7921 AGTGAGCGAG GAAGCGGAAG AGCGCCTGAT GCGGTATTTT CTCCTTACGC ATCTGTGCGG
7981 TATTTCACAC CGCATATGGT GCACTCTCAG TACAATCTGC TCTGATGCCG CATAGTTAAG
8041 CCAGTATACA CTCCGCTATC GCTACGTGAC TGGGTCATGG CTGCGCCCCG ACACCCGCCA
8101 ACACCCGCTG ACGCGCCCTG ACGGGCTTGT CTGCTCCCGG CATCCGCTTA CAGACAAGCT
8161 GTGACCGTCT CCGGGAGCTG CATGTGTCAG AGGTTTTCAC CGTCATCACC GAAACGCGCG
8221 AGGCAGGGTG CCTTGATGTG GGCGCCGGCG GTCGAGTGGC GACGGCGCGG CTTGTCCGCG
8281 CCCTGGTAGA TTGCCTGGCC GTAGGCCAGC CATTTTTGAG CGGCCAGCGG CCGCGATAGG
8341 CCGACGCGAA GCGGCGGGGC GTAGGGAGCG CAGCGACCGA AGGGTAGGCG CTTTTTGCAG
8401 CTCTTCGGCT GTGCGCTGGC CAGACAGTTA TGCACAGGCC AGGCGGGTTT TAAGAGTTTT
8461 AATAAGTTTT AAAGAGTTTT AGGCGGAAAA ATCGCCTTTT TTCTCTTTTA TATCAGTCAC
8521 TTACATGTGT GACCGGTTCC CAATGTACGG CTTTGGGTTC CCAATGTACG GGTTCCGGTT
8581 CCCAATGTAC GGCTTTGGGT TCCCAATGTA CGTGCTATCC ACAGGAAAGA GACCTTTTCG
8641 ACCTTTTTCC CCTGCTAGGG CAATTTGCCC TAGCATCTGC TCCGTACATT AGGAACCGGC
8701 GGATGCTTCG CCCTCGATCA GGTTGCGGTA GCGCATGACT AGGATCGGGC CAGCCTGCCC
8761 CGCCTCCTCC TTCAAATCGT ACTCCGGCAG GTCATTTGAC CCGATCAGCT TGCGCACGGT
8821 GAAACAGAAC TTCTTGAACT CTCCGGCGCT GCCACTGCGT TCGTAGATCG TCTTGAACAA
8881 CCATCTGGCT TCTGCCTTGC CTGCGGCGCG GCGTGCCAGG CGGTAGAGAA AACGGCCGAT
8941 GCCGGGATCG ATCAAAAAGT AATCGGGGTG AACCGTCAGC ACGTCCGGGT TCTTGCCTTC
9001 TGTGATCTCG CGGTACATCC AATCAGCTAG CTCGATCTCG ATGTACTCCG GCCGCCCGGT
9061 TTCGCTCTTT ACGATCTTGT AGCGGCTAAT CAAGGCTTCA CCCTCGGATA CCGTCACCAG
9121 GCGGCCGTTC TTGGCCTTCT TCGTACGCTG CATGGCAACG TGCGTGGTGT TTAACCGAAT
9181 GCAGGTTTCT ACCAGGTCGT CTTTCTGCTT TCCGCCATCG GCTCGCCGGC AGAACTTGAG
9241 TACGTCCGCA ACGTGTGGAC GGAACACGCG GCCGGGCTTG TCTCCCTTCC CTTCCCGGTA
9301 TCGGTTCATG GATTCGGTTA GATGGGAAAC CGCCATCAGT ACCAGGTCGT AATCCCACAC
9361 ACTGGCCATG CCGGCCGGCC CTGCGGAAAC CTCTACGTGC CCGTCTGGAA GCTCGTAGCG
9421 GATCACCTCG CCAGCTCGTC GGTCACGCTT CGACAGACGG AAAACGGCCA CGTCCATGAT
9481 GCTGCGACTA TCGCGGGTGC CCACGTCATA GAGCATCGGA ACGAAAAAAT CTGGTTGCTC
9541 GTCGCCCTTG GGCGGCTTCC TAATCGACGG CGCACCGGCT GCCGGCGGTT GCCGGGATTC
9601 TTTGCGGATT CGATCAGCGG CCGCTTGCCA CGATTCACCG GGCGTGCTT CTGCCTCGAT
9661 GCGTTGCCGC TGGGCGGCCT GCGCGGCCTT CAACTTCTCC ACCAGGTCAT CACCCAGCGC
9721 CGCGCCGATT TGTACCGGGC CGGATGGTTT GCGACCGCTC ACGCCGATTC CTCGGGCTTG
9781 GGGGTTCCAG TGCCATTGCA GGGCCGGCAG ACAACCCAGC CGCTTACGCC TGGCCAACCG
9841 CCCGTTCCTC CACACATGGG GCATTCCACG GCGTCGGTGC CTGGTTGTTC TTGATTTTCC
9901 ATGCCGCCTC CTTTAGCCGC TAAAATTCAT CTACTCATTT ATTCATTTGC TCATTTACTC
9961 TGGTAGCTGC GCGATGTATT CAGATAGCAG CTCGGTAATG GTCTTGCCTT GGCGTACCGC
10021 GTACATCTTC AGCTTGGTGT GATCCTCCGC CGGCAACTGA AAGTTGACCC GCTTCATGGC
10081 TGGCGTGTCT GCCAGGCTGG CCAACGTTGC AGCCTTGCTG CTGCGTGCGC TCGGACGGCC
10141 GGCACTTAGC GTGTTTGTGC TTTTGCTCAT TTTCTCTTTA CCTCATTAAC TCAAATGAGT
10201 TTTGATTTAA TTTCAGCGGC CAGCGCCTGG ACCTCGCGGG CAGCGTCGCC CTCGGGTTCT
10261 GATTCAAGAA CGGTTGTGCC GGCGGCGGCA GTGCCTGGGT AGCTCACGCG CTGCGTGATA
10321 CGGGACTCAA GAATGGGCAG CTCGTACCCG GCCAGCGCCT CGGCAACCTC ACCGCCGATG
10381 CGCGTGCCTT TGATCGCCCG CGACACGACA AAGGCCGCTT GTAGCCTTCC ATCCGTGACC
10441 TCAATGCGCT GCTTAACCAG CTCCACCAGG TCGGCGGTGG CCCATATGTC GTAAGGGCTT
10501 GGCTGCACCG GAATCAGCAC GAAGTCGGCT GCCTTGATCG CGGACACAGC CAAGTCCGCC
10561 GCCTGGGGCG CTCCGTCGAT CACTACGAAG TCGCGCCGGC CGATGGCCTT CACGTCGCGG

10621 TCAATCGTCG GGCGGTCGAT GCCGACAACG GTTAGCGGTT GATCTTCCCG CACGGCCGCC
10681 CAATCGCGGG CACTGCCCTG GGGATCGGAA TCGACTAACA GAACATCGGC CCCGGCGAGT
10741 TGCAGGGCGC GGGCTAGATG GGTTGCGATG GTCGTCTTGC CTGACCCGCC TTTCTGGTTA
10801 AGTACAGCGA TAACCTTCAT GCGTTCCCCT TGCGTATTTG TTTATTTACT CATCGCATCA
10861 TATACGCAGC GACCGCATGA CGCAAGCTGT TTTACTCAAA TACACATCAC CTTTTTAGAC
10921 GGCGGCGCTC GGTTTCTTCA GCGGCCAAGC TGGCCGGCCA GGCCGCCAGC TTGGCATCAG
10981 ACAAACCGGC CAGGATTTCA TGCAGCCGCA CGGTTGAGAC GTGCGCGGGC GGCTCGAACA
11041 CGTACCCGGC CGCGATCATC TCCGCCTCGA TCTCTTCGGT AATGAAAAAC GGTTCGTCCT
11101 GGCCGTCCTG GTGCGGTTTC ATGCTTGTTC CTCTTGGCGT TCATTCTCGG CGGCCGCCAG
11161 GGCGTCGGCC TCGGTCAATG CGTCCTCACG GAAGGCACCG CGCCGCCTGG CCTCGGTGGG
11221 CGTCACTTCC TCGCTGCGCT CAAGTGCGCG GTACAGGGTC GAGCGATGCA CGCCAAGCAG
11281 TGCAGCCGCC TCTTTCACGG TGCGGCCTTC CTGGTCGATC AGCTCGCGGG CGTGCGCGAT
11341 CTGTGCCGGG GTGAGGGTAG GGCGGGGGCC AAACTTCACG CCTCGGGCCT TGGCGGCCTC
11401 GCGCCCGCTC CGGGTGCGGT CGATGATTAG GGAACGCTCG AACTCGGCAA TGCCGGCGAA
11461 CACGGTCAAC ACCATGCGGC CGGCCGGCGT GGTGGTGTCG GCCCACGGCT CTGCCAGGCT
11521 ACGCAGGCCC GCGCCGGCCT CCTGGATGCG CTCGGCAATG TCCAGTAGGT CGCGGGTGCT
11581 GCGGGCCAGG CGGTCTAGCC TGGTCACTGT CACAACGTCG CCAGGGCGTA GGTGGTCAAG
11641 CATCCTGGCC AGCTCCGGGC GGTCGCGCCT GGTGCCGGTG ATCTTCTCGG AAAACAGCTT
11701 GGTGCAGCCG GCCGCGTGCA GTTCGGCCCG TTGGTTGGTC AAGTCCTGGT CGTCGGTGCT
11761 GACGCGGGCA TAGCCCAGCA GGCCAGCGGC GGCGCTCTTG TTCATGGCGT AATGTCTCCG
11821 GTTCTAGTCG CAAGTATTCT ACTTTATGCG ACTAAAACAC GCGACAAGAA AACGCCAGGA
11881 AAAGGGCAGG GCGGCAGCCT GTCGCGTAAC TTAGGACTTG TGCGACATGT CGTTTTCAGA
11941 AGACGGCTGC ACTGAACGTC AGAAGCCGAC TGCACTATAG CAGCGGAGGG GTTGGATCAA
12001 AGTACTTTAA AGTACTTTAA AGTACTTTAA AGTACTTTGA TCCCGAGGGG AACCCTGTGG
12061 TTGGCATGCA CATACAAATG GACGAACGGA TAAACCTTTT CACGCCCTTT TAAATATCCG
12121 ATTATTCTAA TAAACGCTCT TTTCTCTTAG GTTTACCCGC CAATATATCC TGTCAAACAC
12181 TGATAGTTT
//
LOCUS D606_pVAMP721 12208 bp DNA circular UNA 21-DEC-2016
COMMENT
FEATURES Location/Qualifiers
misc_feature join(12055..12208,1..42)
/vntifkey="21"
/label="RB"
/ApEinfo_fwdcolor="pink"
/ApEinfo_revcolor="pink"
/ApEinfo_graphicformat="arrow_data {{0 1 2 0 0 -1} {}
0}width 5 offset 0"
CDS 68..1400
/label="pVAMP721"
misc_feature 68..1400
/label="Promotor Vamp721"
/ApEinfo_fwdcolor="cyan"
/ApEinfo_revcolor="green"
/ApEinfo_graphicformat="arrow_data {{0 1 2 0 0 -1} {}
0}width 5 offset 0"
Site join(745..754,749^750)
/site_type="restriction site"
/note="Name: BsaBI"
/note="Pattern: gatnnnnatc"
/note="Number_of_matches: 5"
/note="cut_0_on_positive_strand: 749^750"
/note="cut_0_on_negative_strand: 749^750"

/note="inhibited_by: "
/note="site_type: other"
/note="restriction site"
/label="BsaBI"
5'UTR 1176..1400
/label="VAMP721 UTR"
CDS 1407..1409
/label="MYC"
/ApEinfo_fwdcolor="pink"
/ApEinfo_revcolor="pink"
/ApEinfo_graphicformat="arrow_data {{0 1 2 0 0 -1} {}
0}width 5 offset 0"
misc_feature 1410..2126
/label="XhoI-meGFP-SalI"
/ApEinfo_fwdcolor="#ff8000"
/ApEinfo_revcolor="#400040"
/ApEinfo_graphicformat="arrow_data {{0 1 2 0 0 -1} {}
0}width 5 offset 0"
misc_feature 1672..1689
/label="RT-fwd"
/ApEinfo_fwdcolor="cyan"
/ApEinfo_revcolor="green"
/ApEinfo_graphicformat="arrow_data {{0 1 2 0 0 -1} {}
0}width 5 offset 0"
CDS 2130..2159
/label="MYC(1)"
/ApEinfo_label="MYC"
/ApEinfo_fwdcolor="pink"
/ApEinfo_revcolor="pink"
/ApEinfo_graphicformat="arrow_data {{0 1 2 0 0 -1} {}
0}width 5 offset 0"
exon 2163..2352
/label="Exon-1"
misc_feature 2163..4123
/label="gVamp721"
/ApEinfo_fwdcolor="cyan"
/ApEinfo_revcolor="green"
/ApEinfo_graphicformat="arrow_data {{0 1 2 0 0 -1} {}
0}width 5 offset 0"
exon 2859..3000
/label="Exon-2"
misc_feature 2859..2882
/label="RT-rev-1"
/ApEinfo_fwdcolor="#0000a0"
/ApEinfo_revcolor="green"
/ApEinfo_graphicformat="arrow_data {{0 1 2 0 0 -1} {}
0}width 5 offset 0"
exon 3080..3203
/label="Exon-3"
misc_feature 3390..3409
/label="g721 seq primer"
/ApEinfo_fwdcolor="#d70428"
/ApEinfo_revcolor="green"

/ApEinfo_graphicformat="arrow_data {{0 1 2 0 0 -1} {}
0}width 5 offset 0"
exon 3393..3453
/label="Exon-4"
exon 3590..3732
/label="Exon-5"
misc_feature 3661..3723
/label="TMD"
/ApEinfo_fwdcolor="#ffff00"
/ApEinfo_revcolor="green"
/ApEinfo_graphicformat="arrow_data {{0 1 2 0 0 -1} {}
0}width 5 offset 0"
misc_feature 3686..3709
/label="RT-rev-2"
/ApEinfo_fwdcolor="#0000a0"
/ApEinfo_revcolor="green"
/ApEinfo_graphicformat="arrow_data {{0 1 2 0 0 -1} {}
0}width 5 offset 0"
Site join(3687..3696,3691^3692)
/site_type="restriction site"
/note="Name: BsaBI"
/note="Pattern: gatnnnnatc"
/note="Number_of_matches: 5"
/note="cut_0_on_positive_strand: 3691^3692"
/note="cut_0_on_negative_strand: 3691^3692"
/note="inhibited_by: "
/note="site_type: other"
/note="restriction site"
/label="BsaBI"
3'UTR 3733..4123
/label="VAMP721 UTR"
terminator 4149..4354
/label="T35S"
/ApEinfo_fwdcolor="pink"
/ApEinfo_revcolor="pink"
/ApEinfo_graphicformat="arrow_data {{0 1 2 0 0 -1} {}
0}width 5 offset 0"
promoter 4377..4679
/vntifkey="29"
/label="nosP"
/ApEinfo_fwdcolor="pink"
/ApEinfo_revcolor="pink"
/ApEinfo_graphicformat="arrow_data {{0 1 2 0 0 -1} {}
0}width 5 offset 0"
misc_feature 4379..4384
/label="BspHI site"
/ApEinfo_fwdcolor="cyan"
/ApEinfo_revcolor="green"
/ApEinfo_graphicformat="arrow_data {{0 1 2 0 0 -1} {}
0}width 5 offset 0"
CDS 4680..5231
/vntifkey="4"
/label="BarCO"

```
                     /ApEinfo_fwdcolor="pink"
                     /ApEinfo_revcolor="pink"
                     /ApEinfo_graphicformat="arrow_data {{0 1 2 0 0 -1} {}
                     0}width 5 offset 0"
     Site            join(5032..5041,5036^5037)
                     /site_type="restriction site"
                     /note="Name: BsaBI"
                     /note="Pattern: gatnnnnatc"
                     /note="Number_of_matches: 5"
                     /note="cut_0_on_positive_strand: 5036^5037"
                     /note="cut_0_on_negative_strand: 5036^5037"
                     /note="inhibited_by: "
                     /note="site_type: other"
                     /note="restriction site"
                     /label="BsaBI"
     terminator      5232..5520
                     /vntifkey="43"
                     /label="nosT"
                     /ApEinfo_fwdcolor="pink"
                     /ApEinfo_revcolor="pink"
                     /ApEinfo_graphicformat="arrow_data {{0 1 2 0 0 -1} {}
                     0}width 5 offset 0"
     Site            join(5310..5319,5314^5315)
                     /site_type="restriction site"
                     /note="Name: BsaBI"
                     /note="Pattern: gatnnnnatc"
                     /note="Number_of_matches: 5"
                     /note="cut_0_on_positive_strand: 5314^5315"
                     /note="cut_0_on_negative_strand: 5314^5315"
                     /note="inhibited_by: "
                     /note="site_type: other"
                     /note="restriction site"
                     /label="BsaBI"
     misc_feature    complement(5541..5870)
                     /vntifkey="21"
                     /label="LB"
                     /ApEinfo_fwdcolor="pink"
                     /ApEinfo_revcolor="pink"
                     /ApEinfo_graphicformat="arrow_data {{0 1 2 0 0 -1} {}
                     0}width 5 offset 0"
     CDS             5876..7120
                     /vntifkey="4"
                     /label="SmR"
                     /ApEinfo_fwdcolor="pink"
                     /ApEinfo_revcolor="pink"
                     /ApEinfo_graphicformat="arrow_data {{0 1 2 0 0 -1} {}
                     0}width 5 offset 0"
     rep_origin      7394..8074
                     /vntifkey="33"
                     /label="pBR322 ori"
                     /ApEinfo_fwdcolor="pink"
                     /ApEinfo_revcolor="pink"
                     /ApEinfo_graphicformat="arrow_data {{0 1 2 0 0 -1} {}
```

0}width 5 offset 0"
rep_origin 8484..11078
/vntifkey="33"
/label="pVS1 ori"
/ApEinfo_fwdcolor="pink"
/ApEinfo_revcolor="pink"
/ApEinfo_graphicformat="arrow_data {{0 1 2 0 0 -1} {}
0}width 5 offset 0"
Site join(10722..10731,10726^10727)
/site_type="restriction site"
/note="Name: BsaBI"
/note="Pattern: gatnnnnatc"
/note="Number_of_matches: 5"
/note="cut_0_on_positive_strand: 10726^10727"
/note="cut_0_on_negative_strand: 10726^10727"
/note="inhibited_by: "
/note="site_type: other"
/note="restriction site"
/label="BsaBI"
ORIGIN
1 AAACTGAAGG CGGGAAACGA CAATCTGATC CAAGCTCAAG CTAAGCTAGG ACTAGCGCTC
61 GCACGTGTCT TCCTTTTATC CAGATGAGAG AAAGATAAAA AAACGAAACC TAAGAACACC
121 ACAAAAATCC GATCGAATTG ATGAGAGAGA TTAGTAAGCG ATGTAAAAAT ACACAAGGGA
181 ACGAGGGGTA GCGATGGAGA AAGCAGGACG CAGAGATCGG AGAAGAGAAA CTATATGAGA
241 GCGCTGGATT TGCGCGAGTA ACAGAGACAT TATTTTCCCT TTGGTAACGT TCAAATTTTG
301 AATTTTTAAA TATAAAAAGT AAATAAAAAG AAAAATTCAT CATTTGTATA GTCGAAAATT
361 GCCTTAATAA ACTTTATTTT GGTAAATAAT TCACGTCTAA TTAATAATAG TATCTTCATG
421 TTTTTACACC ACGGTGGCTC GTTTTTCAAT TCAAATGTGA AATGTTTATC AATGAGCATT
481 TTAAAAAACA AATATATATA TATATATATA TATTATATAT TGATTTTTTA TAAAGAAAAT
541 TGCGATGTGT AAGGGATATT TGAGTTGTTA AATTTTCTAT AAATTATATG TTATGTTACT
601 TAAAAGAACG TTCAATTGAT AATGTTGCAA ACCTTCCTAC CAAAAAAATT AAAATAATGT
661 TGCAAACCTT TTCTTTCATT CCAATTCACT AACCTTCTAT TCTAACCACA ACCTAACTAC
721 CTAAATTTCA ATTATTTGAT TGTTGATCAA AATCTTTAGG AACACGTTAA ACAAATTAAT
781 TAGCAGGTTC TAAACAATGC AGAGCAATTG TAGATGACAA GAGATAACGG TAGAAGTTCG
841 AATTTTGTTG CTGTAACAAG ATACTATGAC TATGAGCTCC CCCCGTCCAT TAAGAATTAA
901 GAAATTTGGT TAATTACAAC ATTTGTTTTT ATGTTTAAGC CTTTTGTTCG TAACTGAGCT
961 AATACTGGAA TCACTGATCA ATGATGAATT TGGAAAAAAA GATATTTTGA GGAAGACAGA
1021 GACAGACTCT CTTCGACCAT GGGACCTGAA CTTGTCCTAA AGTTGGTAAT TACTAATTTG
1081 TGCCATCGGC AGAGTCCACG AGCTTTTTGG CTGGATCCGG GCGGGTCCAA CTTTCTCGAC
1141 ATTTGTTTGA TCTCTCCGTT TTGAAATTGT TTTTTTTTAG TCCTTAAATT CAAGGTTTTA
1201 TACTATTTTT GACCCAATCT GACTACCTTA AATGACAACT TTACCCTTGC CCGCTTTTCA
1261 ACTTTTTCTT CTCCCATGTC CACTTTAGTC CTCGTTTTGT TCTTTCATAG TCTTCTAGAT
1321 CTGTAGTTTT TTTCCGCGAC CGCCCAAAAA ATTCTCTGAT CAGACGGTCG ATTGGATCGG
1381 AGATTTAAGG TAAAGAAAAA ATGCTCGAGA TGGTGAGCAA GGGCGAGGAG CTGTTCACCG
1441 GGGTGGTGCC CATCCTGGTC GAGCTGGACG GCGACGTAAA CGGCCACAAG TTCAGCGTGT
1501 CCGGCGAGGG CGAGGGCGAT GCCACCTACG GCAAGCTGAC CCTGAAGTTC ATCTGCACCA
1561 CCGGCAAGCT GCCCGTGCCC TGGCCCACCC TCGTGACCAC CCTGACCTAC GGCGTGCAGT
1621 GCTTCAGCCG CTACCCCGAC CACATGAAGC AGCACGACTT CTTCAAGTCC GCCATGCCCG
1681 AAGGCTACGT CCAGGAGCGC ACCATCTTCT TCAAGGACGA CGGCAACTAC AAGACCCGCG
1741 CCGAGGTGAA GTTCGAGGGC GACACCCTGG TGAACCGCAT CGAGCTGAAG GGCATCGACT
1801 TCAAGGAGGA CGGCAACATC CTGGGGCACA AGCTGGAGTA CAACTACAAC AGCCACAACG
1861 TCTATATCAT GGCCGACAAG CAGAAGAACG GCATCAAGGT GAACTTCAAG ATCCGCCACA
1921 ACATCGAGGA CGGCAGCGTG CAGCTCGCCG ACCACTACCA GCAGAACACC CCCATCGGCG

1981 ACGGCCCCGT GCTGCTGCCC GACAACCACT ACCTGAGTAC TCAGTCCAAG CTGAGCAAAG
2041 ACCCCAACGA GAAGCGCGAT CACATGGTCC TGCTGGAGTT CGTGACCGCC GCCGGGATCA
2101 CTCTCGGCAT GGACGAGCTG TACAAGGTCG AGCAGAAGTT GATCTCAGAG GAGGACTTAG
2161 GCATGGCGCA ACAATCGTTG ATCTACAGTT TCGTAGCTCG CGGCACGGTG ATCCTCGTTG
2221 AGTTCACTGA TTTCAAAGGT AATTTCACCT CAATCGCTGC TCAGTGCCTC CAGAAGCTTC
2281 CGTCTTCGAA CAACAAGTTC ACCTACAACT GCGACGGTCA TACCTTCAAT TACCTTGTCG
2341 AAGATGGATT CAGTAAGTCA CTTTTCGTTT GATCTATGCA TAGGTTTTAA TCTTACACCA
2401 TTCGGTGCTT ACCTCGATTT GTTTCTAGGT TTATTTGCCT AACCTATCGA TTCGTATGGA
2461 TTTGTGCATC CTGAGTAGTC TGATTTCACT GAGGAACCGC ATGAACCAAG CTTTTAGGGT
2521 TAGATCTATA TTTATTAAAC GCAATAAATC TCTGTAGAGT CTTACCTGTA TAATCACTCA
2581 GATTTGGACA GAATCCGTGG TAGCTTTGGT TGATACTATG TGGAAGAACA CCAACATTGA
2641 TCTGAGACAG GGTTTTTACT GGTTGCTTTA TTTGCGTATA GTTTCCTAGG TCATGGGTTT
2701 TGGCTGGCTT CATTTTTCAA TGAAAATCTG GGGTCTTATT GTAAGCTTTT GTGACTATCC
2761 TTTAGTTGTT GTAGCCATAT CAATTTCATA TTAAATCAAT GTCACCTGAT TATCCCGTAA
2821 GGGAAGACAA GGAGATTGAT TTTGTTCCTG TTGTACAGCC TATTGTGTTG TTGCGGTTGA
2881 TTCTGCTGGG AGGCAAATTC CCATGTCCTT TTTGGAAAGA GTAAAAGAAG ATTTTAACAA
2941 GCGATATGGT GGTGGAAAGG CTGCAACTGC TCAAGCAAAC AGCTTGAATA AGGAGTTTGG
3001 GTACTTTTTC ATTATCTCTT CTATTTGGAT GATCTTCTCT TTTATATTCG TGGACTGACT
3061 CTTTTTGGTA ACTTTGAAGC TCTAAACTGA AAGAGCATAT GCAGTATTGC ATGGATCATC
3121 CTGATGAGAT TAGCAAGCTT GCTAAGGTGA AGGCGCAAGT GTCAGAAGTT AAGGGTGTAA
3181 TGATGGAAAA CATTGAGAAG GTTTGAATCT GACCCTTTCT GTCCTCAATG TATATATTTA
3241 CATCTATGGT TGACCCATCT GAAAGAGCTC ATCTAGTCAT ACTAAGTTAC TGTGAAATCA
3301 ATTACTAATA ATTCAATGCA TTACATCTTA CGGGAAATGT CCAGTTTACC AATGTAACAC
3361 CGCTTACATA TGCGACTCAT TATTTGCAGG TTCTTGACCG TGGTGAGAAA ATTGAGCTTT
3421 TGGTGGACAA AACCGAAAAT CTTCGCTCAC AGGTTAGACT CTCTCATACC CTTATCTCCT
3481 GCATCTATTT GCCTACACTC ACACACAAAG ATGAACATGC TTTTTGAGTT CATAGCTGAC
3541 TCATCCTATT ACCATATCTT GTGTCTAACT GAACTCAAAC TGCCACAGGC ACAAGATTTC
3601 AGAACAACAG GAACGCAGAT GAGAAGAAAG ATGTGGCTTC AGAACATGAA GATAAAACTC
3661 ATAGTGCTCG CCATCATTAT CGCACTGATT CTCATCATCG TGCTCTCAGT TTGCCATGGG
3721 TTTAAGTGTT AAGCCCAGAA AATTCAAAAC TTCTGAGATC TTTCTCACTC GGTACATCGT
3781 AGTGCCTTCT CTTTTATTGT TCGATCTTCA ATCTAATGTT TCTGCTTGAA TTGGTTTTGT
3841 ATAGATAGAC ATATATGTAT GCTACTCATT TGCTATATTG CTTGCATCTG ATGACATTGA
3901 TCCCTTGCGA ACCAGTCAGT ACTTGACAAG AGATCCGAAA CTATTAAATG ATTTTGTTGA
3961 CTGAAGATTG CATTGTGGTG GTGATGATAA AAATTTGAGA TTGTATTGAC GATTATACGG
4021 CGTGATGTGT GCTGGTTACT TTAAATTTAG ATAAGAGTAG TGCCCTCTCT ATATCGTTAC
4081 TAGGGATGGT AGTACATATA TATACGTACG TCATATTCCG ACCGTTTTAA GCGGCCATGC
4141 TAGAGTCCGC AAAAATCACC AGTCTCTCTC TACAAATCTA TCTCTCTCTA TTTTTCTCCA
4201 GAATAATGTG TGAGTAGTTC CCAGATAAGG GAATTAGGGT TCTTATAGGG TTTCGCTCAT
4261 GTGTTGAGCA TATAAGAAAC CCTTAGTATG TATTTGTATT TGTAAAATAC TTCTATCAAT
4321 AAAATTTCTA ATTCCTAAAA CCAAAATCCA GTGACACGTG CGGAGCTCCG TACGTAGATC
4381 ATGAGCGGAG AATTAAGGGA GTCACGTTAT GACCCCCGCC GATGACGCGG GACAAGCCGT
4441 TTTACGTTTG GAACTGACAG AACCGCAACG TTGAAGGAGC CACTGAGCGC GGGTTTCTGG
4501 AGTTTAATGA GCTAAGCACA TACGTCAGAA ACCATTATTG CGCGTTCAAA AGTCGCCTAA
4561 GGTCACTATC ATCTAGCAAA TATTTCTTGT CAAAAATGCT CCACTGACGT TCCATAAATT
4621 CCCCTCGGTA TCCAATTAGA GTCTCATATT CACTCTCAAC TCGATCGAGG GGATCTACCA
4681 TGAGTCCAGA AAGGAGACCA GCAGATATTA GGAGAGCAAC CGAAGCAGAT ATGCCAGCAG
4741 TTTGCACCAT TGTGAACCAT TACATCGAGA CTTCTACAGT TAATTTTAGG ACTGAACCTC
4801 AAGAGCCACA GGAATGGACA GATGATCTCG TGAGATTAAG GGAAAGATAC CCTTGGCTTG
4861 TTGCTGAGGT GGATGGAGAA GTTGCAGGTA TTGCTTACGC AGGACCTTGG AAGGCTAGAA
4921 ACGCTTATGA TTGGACTGCT GAGTCTACCG TTTACGTGTC ACCTAGACAT CAAAGAACCG
4981 GACTTGGTTC AACCTTGTAT ACTCACCTTT TGAAGTCTCT TGAAGCTCAG GGATTCAAAT
5041 CTGTTGTGGC TGTTATCGGT TTGCCTAATG ATCCAAGTGT GAGAATGCAT GAAGCTCTCG
5101 GATACGCACC AAGGGGTATG TTAAGAGCTG CTGGATTTAA ACATGGTAAC TGGCACGATG

5161 TTGGTTTCTG GCAGTTAGAT TTCAGTTTAC CAGTTCCACC AAGACCAGTG CTTCCAGTGA
5221 CCGAGATTTG ACCCGGGCTA GAGTCAAGCA GATCGTTCAA ACATTTGGCA ATAAAGTTTC
5281 TTAATATTGA ATCCTGTTGC CGGTCTTGCG ATGATTATCA TATAATTTCT GTTGAATTAC
5341 GTTAAGCATG TAATAATTAA CATGTAATGC ATGACGTTAT TTATGAGATG GGTTTTTATG
5401 ATTAGAGTCC CGCAATTATA CATTTAATAC GCGATAGAAA ACAAAATATA GCGCGCAAAC
5461 TAGGATAAAT TATCGCGCGC GGTGTCATCT ATGTTACTAG ATCGACCGGC ATGAAGCTGA
5521 GTTAACGATG TAAGTCCTCA ATTCGGCGTT AATTCAGTAC ATTAAAAACG TCCGCAATGT
5581 GTTATTAAGT TGTCTAAGCG TCAATTTGTT TACACCACAA TATATCCTGC CACCAGCCAG
5641 CCAACAGCTC CCCGACCGGC AGCTCGGCAC AAAATCACCA CTCGATACAG GCAGCCCATC
5701 AGTCCGGGAC GGCGTCAGCG GGAGAGCCGT TGTAAGGCGG CAGACTTTGC TCATGTTACC
5761 GATGCTATTC GGAAGAACGG CAACTAAGCT GCCGGGTTTG AAACACGGAT GATCTCGCGG
5821 AGGGTAGCAT GTTGATTGTA ACGATGACAG AGCGTTGCTG CCTGTGATCA ATTCGGGCAC
5881 GAACCCAGTG GACATAAGCC TGTTCGGTTC GTAAGCTGTA ATGCAAGTAG CGTATGCGCT
5941 CACGCAACTG GTCCAGAACC TTGACCGAAC GCAGCGGTGG TAACGGCGCA GTGGCGGTTT
6001 TCATGGCTTG TTATGACTGT TTTTTTGGGG TACAGTCTAT GCCTCGGGCA TCCAAGCAGC
6061 AAGCGCGTTA CGCCGTGGGT CGATGTTTGA TGTTATGGAG CAGCAACGAT GTTACGCAGC
6121 AGGGCAGTCG CCCTAAAACA AAGTTAAACA TCATGGGGGA AGCGGTGATC GCCGAAGTAT
6181 CGACTCAACT ATCAGAGGTA GTTGGCGTCA TCGAGCGCCA TCTCGAACCG ACGTTGCTGG
6241 CCGTACATTT GTACGGCTCC GCAGTGGATG GCGGCCTGAA GCCACACAGT GATATTGATT
6301 TGCTGGTTAC GGTGACCGTA AGGCTTGATG AAACAACGCG GCGAGCTTTG ATCAACGACC
6361 TTTTGGAAAC TTCGGCTTCC CCTGGAGAGA GCGAGATTCT CCGCGCTGTA GAAGTCACCA
6421 TTGTTGTGCA CGACGACATC ATTCCGTGGC GTTATCCAGC TAAGCGCGAA CTGCAATTTG
6481 GAGAATGGCA GCGCAATGAC ATTCTTGCAG GTATCTTCGA GCCAGCCACG ATCGACATTG
6541 ATCTGGCTAT CTTGCTGACA AAAGCAAGAG AACATAGCGT TGCCTTGGTA GGTCCAGCGG
6601 CGGAGGAACT CTTTGATCCG GTTCCTGAAC AGGATCTATT TGAGGCGCTA AATGAAACCT
6661 TAACGCTATG GAACTCGCCG CCCGACTGGG CTGGCGATGA GCGAAATGTA GTGCTTACGT
6721 TGTCCCGCAT TTGGTACAGC GCAGTAACCG GCAAAATCGC GCCGAAGGAT GTCGCTGCCG
6781 ACTGGGCAAT GGAGCGCCTG CCGGCCCAGT ATCAGCCCGT CATACTTGAA GCTAGACAGG
6841 CTTATCTTGG ACAAGAAGAA GATCGCTTGG CCTCGCGCGC AGATCAGTTG GAAGAATTTG
6901 TCCACTACGT GAAAGGCGAG ATCACCAAGG TAGTCGGCAA ATAATGTCTA GCTAGAAATT
6961 CGTTCAAGCC GACGCCGCTT CGCGGCGCGG CTTAACTCAA GCGTTAGATG CACTAAGCAC
7021 ATAATTGCTC ACAGCCAAAC TATCAGGTCA AGTCTGCTTT TATTATTTTT AAGCGTGCAT
7081 AATAAGCCCT ACACAAATTG GGAGATATAT CATGCATGAC CAAAATCCCT TAACGTGAGT
7141 TTTCGTTCCA CTGAGCGTCA GACCCCGTAG AAAAGATCAA AGGATCTTCT TGAGATCCTT
7201 TTTTTCTGCG CGTAATCTGC TGCTTGCAAA CAAAAAAACC ACCGCTACCA GCGGTGGTTT
7261 GTTTGCCGGA TCAAGAGCTA CCAACTCTTT TTCCGAAGGT AACTGGCTTC AGCAGAGCGC
7321 AGATACCAAA TACTGTCCTT CTAGTGTAGC CGTAGTTAGG CCACCACTTC AAGAACTCTG
7381 TAGCACCGCC TACATACCTC GCTCTGCTAA TCCTGTTACC AGTGGCTGCT GCCAGTGGCG
7441 ATAAGTCGTG TCTTACCGGG TTGGACTCAA GACGATAGTT ACCGGATAAG GCGCAGCGGT
7501 CGGGCTGAAC GGGGGGTTCG TGCACACAGC CCAGCTTGGA GCGAACGACC TACACCGAAC
7561 TGAGATACCT ACAGCGTGAG CTATGAGAAA GCGCCACGCT TCCCGAAGGG AGAAAGGCGG
7621 ACAGGTATCC GGTAAGCGGC AGGGTCGGAA CAGGAGAGCG CACGAGGGAG CTTCCAGGGG
7681 GAAACGCCTG GTATCTTTAT AGTCCTGTCG GGTTTCGCCA CCTCTGACTT GAGCGTCGAT
7741 TTTTGTGATG CTCGTCAGGG GGGCGGAGCC TATGGAAAAA CGCCAGCAAC GCGGCCTTTT
7801 TACGGTTCCT GGCCTTTTGC TGGCCTTTTG CTCACATGTT CTTTCCTGCG TTATCCCCTG
7861 ATTCTGTGGA TAACCGTATT ACCGCCTTTG AGTGAGCTGA TACCGCTCGC CGCAGCCGAA
7921 CGACCGAGCG CAGCGAGTCA GTGAGCGAGG AAGCGGAAGA GCGCCTGATG CGGTATTTTC
7981 TCCTTACGCA TCTGTGCGGT ATTTCACACC GCATATGGTG CACTCTCAGT ACAATCTGCT
8041 CTGATGCCGC ATAGTTAAGC CAGTATACAC TCCGCTATCG CTACGTGACT GGGTCATGGC
8101 TGCGCCCCGA CACCCGCCAA CACCCGCTGA CGCGCCCTGA CGGGCTTGTC TGCTCCCGGC
8161 ATCCGCTTAC AGACAAGCTG TGACCGTCTC CGGGAGCTGC ATGTGTCAGA GGTTTTCACC
8221 GTCATCACCG AAACGCGCGA GGCAGGGTGC CTTGATGTGG GCGCCGGCGG TCGAGTGGCG
8281 ACGGCGCGGC TTGTCCGCGC CCTGGTAGAT TGCCTGGCCG TAGGCCAGCC ATTTTTGAGC

8341 GGCCAGCGGC CGCGATAGGC CGACGCGAAG CGGCGGGGCG TAGGGAGCGC AGCGACCGAA
8401 GGGTAGGCGC TTTTTGCAGC TCTTCGGCTG TGCGCTGGCC AGACAGTTAT GCACAGGCCA
8461 GGCGGGTTTT AAGAGTTTTA ATAAGTTTTA AAGAGTTTTA GGCGGAAAAA TCGCCTTTTT
8521 TCTCTTTTAT ATCAGTCACT TACATGTGTG ACCGGTTCCC AATGTACGGC TTTGGGTTCC
8581 CAATGTACGG GTTCCGGTTC CCAATGTACG GCTTTGGGTT CCCAATGTAC GTGCTATCCA
8641 CAGGAAAGAG ACCTTTTCGA CCTTTTTCCC CTGCTAGGGC AATTTGCCCT AGCATCTGCT
8701 CCGTACATTA GGAACCGGCG GATGCTTCGC CCTCGATCAG GTTGCGGTAG CGCATGACTA
8761 GGATCGGGCC AGCCTGCCCC GCCTCCTCCT TCAAATCGTA CTCCGGCAGG TCATTTGACC
8821 CGATCAGCTT GCGCACGGTG AAACAGAACT TCTTGAACTC TCCGGCGCTG CCACTGCGTT
8881 CGTAGATCGT CTTGAACAAC CATCTGGCTT CTGCCTTGCC TGCGGCGCGG CGTGCCAGGC
8941 GGTAGAGAAA ACGGCCGATG CCGGGATCGA TCAAAAAGTA ATCGGGGTGA ACCGTCAGCA
9001 CGTCCGGGTT CTTGCCTTCT GTGATCTCGC GGTACATCCA ATCAGCTAGC TCGATCTCGA
9061 TGTACTCCGG CCGCCCGGTT TCGCTCTTTA CGATCTTGTA GCGGCTAATC AAGGCTTCAC
9121 CCTCGGATAC CGTCACCAGG CGGCCGTTCT TGGCCTTCTT CGTACGCTGC ATGGCAACGT
9181 GCGTGGTGTT TAACCGAATG CAGGTTTCTA CCAGGTCGTC TTTCTGCTTT CCGCCATCGG
9241 CTCGCCGGCA GAACTTGAGT ACGTCCGCAA CGTGTGGACG GAACACGCGG CCGGGCTTGT
9301 CTCCCTTCCC TTCCCGGTAT CGGTTCATGG ATTCGGTTAG ATGGGAAACC GCCATCAGTA
9361 CCAGGTCGTA ATCCCACACA CTGGCCATGC CGGCCGGCCC TGCGGAAACC TCTACGTGCC
9421 CGTCTGGAAG CTCGTAGCGG ATCACCTCGC CAGCTCGTCG GTCACGCTTC GACAGACGGA
9481 AAACGGCCAC GTCCATGATG CTGCGACTAT CGCGGGTGCC CACGTCATAG AGCATCGGAA
9541 CGAAAAAATC TGGTTGCTCG TCGCCCTTGG GCGGCTTCCT AATCGACGGC GCACCGGCTG
9601 CCGGCGGTTG CCGGGATTCT TTGCGGATTC GATCAGCGGC CGCTTGCCAC GATTCACCGG
9661 GGCGTGCTTC TGCCTCGATG CGTTGCCGCT GGGCGGCCTG CGCGGCCTTC AACTTCTCCA
9721 CCAGGTCATC ACCCAGCGCC GCGCCGATTT GTACCGGGCC GGATGGTTTG CGACCGCTCA
9781 CGCCGATTCC TCGGGCTTGG GGGTTCCAGT GCCATTGCAG GGCCGGCAGA CAACCCAGCC
9841 GCTTACGCCT GGCCAACCGC CCGTTCCTCC ACACATGGGG CATTCCACGG CGTCGGTGCC
9901 TGGTTGTTCT TGATTTTCCA TGCCGCCTCC TTTAGCCGCT AAAATTCATC TACTCATTTA
9961 TTCATTTGCT CATTTACTCT GGTAGCTGCG CGATGTATTC AGATAGCAGC TCGGTAATGG
10021 TCTTGCCTTG GCGTACCGCG TACATCTTCA GCTTGGTGTG ATCCTCCGCC GGCAACTGAA
10081 AGTTGACCCG CTTCATGGCT GGCGTGTCTG CCAGGCTGGC CAACGTTGCA GCCTTGCTGC
10141 TGCGTGCGCT CGGACGGCCG GCACTTAGCG TGTTTGTGCT TTTGCTCATT TTCTCTTTAC
10201 CTCATTAACT CAAATGAGTT TTGATTTAAT TTCAGCGGCC AGCGCCTGGA CCTCGCGGGC
10261 AGCGTCGCCC TCGGGTTCTG ATTCAAGAAC GGTTGTGCCG GCGGCGGCAG TGCCTGGGTA
10321 GCTCACGCGC TGCGTGATAC GGGACTCAAG AATGGGCAGC TCGTACCCGG CCAGCGCCTC
10381 GGCAACCTCA CCGCCGATGC GCGTGCCTTT GATCGCCCGC GACACGACAA AGGCCGCTTG
10441 TAGCCTTCCA TCCGTGACCT CAATGCGCTG CTTAACCAGC TCCACCAGGT CGGCGGTGGC
10501 CCATATGTCG TAAGGGCTTG GCTGCACCGG AATCAGCACG AAGTCGGCTG CCTTGATCGC
10561 GGACACAGCC AAGTCCGCCG CCTGGGGCGC TCCGTCGATC ACTACGAAGT CGCGCCGGCC
10621 GATGGCCTTC ACGTCGCGGT CAATCGTCGG GCGGTCGATG CCGACAACGG TTAGCGGTTG
10681 ATCTTCCCGC ACGGCCGCCC AATCGCGGGC ACTGCCCTGG GGATCGGAAT CGACTAACAG
10741 AACATCGGCC CCGGCGAGTT GCAGGGCGCG GGCTAGATGG GTTGCGATGG TCGTCTTGCC
10801 TGACCCGCCT TTCTGGTTAA GTACAGCGAT AACCTTCATG CGTTCCCCTT GCGTATTTGT
10861 TTATTTACTC ATCGCATCAT ATACGCAGCG ACCGCATGAC GCAAGCTGTT TTACTCAAAT
10921 ACACATCACC TTTTTAGACG GCGGCGCTCG GTTTCTTCAG CGGCCAAGCT GGCCGGCCAG
10981 GCCGCCAGCT TGGCATCAGA CAAACCGGCC AGGATTTCAT GCAGCCGCAC GGTTGAGACG
11041 TGCGCGGGCG GCTCGAACAC GTACCCGGCC GCGATCATCT CCGCCTCGAT CTCTTCGGTA
11101 ATGAAAAACG GTTCGTCCTG GCCGTCCTGG TGCGGTTTCA TGCTTGTTCC TCTTGGCGTT
11161 CATTCTCGGC GGCCGCCAGG GCGTCGGCCT CGGTCAATGC GTCCTCACGG AAGGCACCGC
11221 GCCGCCTGGC CTCGGTGGGC GTCACTTCCT CGCTGCGCTC AAGTGCGCGG TACAGGGTCG
11281 AGCGATGCAC GCCAAGCAGT GCAGCCGCCT CTTTCACGGT GCGGCCTTCC TGGTCGATCA
11341 GCTCGCGGGC GTGCGCGATC TGTGCCGGGG TGAGGGTAGG GCGGGGGCCA AACTTCACGC
11401 CTCGGGCCTT GGCGGCCTCG CGCCCGCTCC GGGTGCGGTC GATGATTAGG GAACGCTCGA
11461 ACTCGGCAAT GCCGGCGAAC ACGGTCAACA CCATGCGGCC GGCCGGCGTG GTGGTGTCGG

11521 CCCACGGCTC TGCCAGGCTA CGCAGGCCCG CGCCGGCCTC CTGGATGCGC TCGGCAATGT
11581 CCAGTAGGTC GCGGGTGCTG CGGGCCAGGC GGTCTAGCCT GGTCACTGTC ACAACGTCGC
11641 CAGGGCGTAG GTGGTCAAGC ATCCTGGCCA GCTCCGGGCG GTCGCGCCTG GTGCCGGTGA
11701 TCTTCTCGGA AAACAGCTTG GTGCAGCCGG CCGCGTGCAG TTCGGCCCGT TGGTTGGTCA
11761 AGTCCTGGTC GTCGGTGCTG ACGCGGGCAT AGCCCAGCAG GCCAGCGGCG GCGCTCTTGT
11821 TCATGGCGTA ATGTCTCCGG TTCTAGTCGC AAGTATTCTA CTTTATGCGA CTAAAACACG
11881 CGACAAGAAA ACGCCAGGAA AAGGGCAGGG CGGCAGCCTG TCGCGTAACT TAGGACTTGT
11941 GCGACATGTC GTTTTCAGAA GACGGCTGCA CTGAACGTCA GAAGCCGACT GCACTATAGC
12001 AGCGGAGGGG TTGGATCAAA GTACTTTAAA GTACTTTAAA GTACTTTAAA GTACTTTGAT
12061 CCCGAGGGGA ACCCTGTGGT TGGCATGCAC ATACAAATGG ACGAACGGAT AAACCTTTTC
12121 ACGCCCTTTT AAATATCCGA TTATTCTAAT AAACGCTCTT TTCTCTTAGG TTTACCCGCC
12181 AATATATCCT GTCAAACACT GATAGTTT
//
LOCUS D607_pVAMP721Y57D 12208 bp DNA circular UNA 16-AUG-2019
COMMENT ApEinfo:methylated:1
FEATURES Location/Qualifiers
misc_feature join(12055..12208,1..42)
/vntifkey="21"
/locus_tag="RB"
misc_feature 68..1400
/locus_tag="Promotor Vamp721"
CDS 1407..1409
/locus_tag="MYC(1)"
misc_feature 1410..2126
/locus_tag="XhoI-meGFP-SalI"
CDS 2130..2159
/locus_tag="MYC"
misc_feature join(2163..2306,2347..4123)
/locus_tag="gVamp721"
misc_feature 2307..2346
/locus_tag="mutated sequence Y57D + MscI site"
misc_feature 3390..3409
/locus_tag="g721 seq primer"
terminator 4149..4354
/locus_tag="T35S"
promoter 4377..4679
/vntifkey="29"
/locus_tag="nosP"
misc_feature 4379..4384
/locus_tag="BspHI site"
CDS 4680..5231
/vntifkey="4"
/locus_tag="BarCO"
terminator 5232..5520
/vntifkey="43"
/locus_tag="nosT"
misc_feature complement(5541..5870)
/vntifkey="21"
/locus_tag="LB"
CDS 5876..7120
/vntifkey="4"
/locus_tag="SmR"
rep_origin 7394..8074

/vntifkey="33"
/locus_tag="pBR322 ori"
rep_origin 8484..11078
/vntifkey="33"
/locus_tag="pVS1 ori"
ORIGIN
1 AAACTGAAGG CGGGAAACGA CAATCTGATC CAAGCTCAAG CTAAGCTAGG ACTAGCGCTC
61 GCACGTGTCT TCCTTTTATC CAGATGAGAG AAAGATAAAA AAACGAAACC TAAGAACACC
121 ACAAAAATCC GATCGAATTG ATGAGAGAGA TTAGTAAGCG ATGTAAAAAT ACACAAGGGA
181 ACGAGGGGTA GCGATGGAGA AAGCAGGACG CAGAGATCGG AGAAGAGAAA CTATATGAGA
241 GCGCTGGATT TGCGCGAGTA ACAGAGACAT TATTTTCCCT TTGGTAACGT TCAAATTTTG
301 AATTTTTAAA TATAAAAAGT AAATAAAAAG AAAAATTCAT CATTTGTATA GTCGAAAATT
361 GCCTTAATAA ACTTTATTTT GGTAAATAAT TCACGTCTAA TTAATAATAG TATCTTCATG
421 TTTTTACACC ACGGTGGCTC GTTTTTCAAT TCAAATGTGA AATGTTTATC AATGAGCATT
481 TTAAAAAACA AATATATATA TATATATATA TATTATATAT TGATTTTTTA TAAAGAAAAT
541 TGCGATGTGT AAGGGATATT TGAGTTGTTA AATTTTCTAT AAATTATATG TTATGTTACT
601 TAAAAGAACG TTCAATTGAT AATGTTGCAA ACCTTCCTAC CAAAAAAATT AAAATAATGT
661 TGCAAACCTT TTCTTTCATT CCAATTCACT AACCTTCTAT TCTAACCACA ACCTAACTAC
721 CTAAATTTCA ATTATTTGAT TGTTGATCAA AATCTTTAGG AACACGTTAA ACAAATTAAT
781 TAGCAGGTTC TAAACAATGC AGAGCAATTG TAGATGACAA GAGATAACGG TAGAAGTTCG
841 AATTTTGTTG CTGTAACAAG ATACTATGAC TATGAGCTCC CCCCGTCCAT TAAGAATTAA
901 GAAATTTGGT TAATTACAAC ATTTGTTTTT ATGTTTAAGC CTTTTGTTCG TAACTGAGCT
961 AATACTGGAA TCACTGATCA ATGATGAATT TGGAAAAAAA GATATTTTGA GGAAGACAGA
1021 GACAGACTCT CTTCGACCAT GGGACCTGAA CTTGTCCTAA AGTTGGTAAT TACTAATTTG
1081 TGCCATCGGC AGAGTCCACG AGCTTTTTGG CTGGATCCGG GCGGGTCCAA CTTTCTCGAC
1141 ATTTGTTTGA TCTCTCCGTT TTGAAATTGT TTTTTTTTAG TCCTTAAATT CAAGGTTTTA
1201 TACTATTTTT GACCCAATCT GACTACCTTA AATGACAACT TTACCCTTGC CCGCTTTTCA
1261 ACTTTTTCTT CTCCCATGTC CACTTTAGTC CTCGTTTTGT TCTTTCATAG TCTTCTAGAT
1321 CTGTAGTTTT TTTCCGCGAC CGCCCAAAAA ATTCTCTGAT CAGACGGTCG ATTGGATCGG
1381 AGATTTAAGG TAAAGAAAAA ATGCTCGAGA TGGTGAGCAA GGGCGAGGAG CTGTTCACCG
1441 GGGTGGTGCC CATCCTGGTC GAGCTGGACG GCGACGTAAA CGGCCACAAG TTCAGCGTGT
1501 CCGGCGAGGG CGAGGGCGAT GCCACCTACG GCAAGCTGAC CCTGAAGTTC ATCTGCACCA
1561 CCGGCAAGCT GCCCGTGCCC TGGCCCACCC TCGTGACCAC CCTGACCTAC GGCGTGCAGT
1621 GCTTCAGCCG CTACCCCGAC CACATGAAGC AGCACGACTT CTTCAAGTCC GCCATGCCCG
1681 AAGGCTACGT CCAGGAGCGC ACCATCTTCT TCAAGGACGA CGGCAACTAC AAGACCCGCG
1741 CCGAGGTGAA GTTCGAGGGC GACACCCTGG TGAACCGCAT CGAGCTGAAG GGCATCGACT
1801 TCAAGGAGGA CGGCAACATC CTGGGGCACA AGCTGGAGTA CAACTACAAC AGCCACAACG
1861 TCTATATCAT GGCCGACAAG CAGAAGAACG GCATCAAGGT GAACTTCAAG ATCCGCCACA
1921 ACATCGAGGA CGGCAGCGTG CAGCTCGCCG ACCACTACCA GCAGAACACC CCCATCGGCG
1981 ACGGCCCCGT GCTGCTGCCC GACAACCACT ACCTGAGTAC TCAGTCCAAG CTGAGCAAAG
2041 ACCCCAACGA GAAGCGCGAT CACATGGTCC TGCTGGAGTT CGTGACCGCC GCCGGGATCA
2101 CTCTCGGCAT GGACGAGCTG TACAAGGTCG AGCAGAAGTT GATCTCAGAG GAGGACTTAG
2161 GCATGGCGCA ACAATCGTTG ATCTACAGTT TCGTAGCTCG CGGCACGGTG ATCCTCGTTG
2221 AGTTCACTGA TTTCAAAGGT AATTTCACCT CAATCGCTGC TCAGTGCCTC CAGAAGCTTC
2281 CGTCTTCGAA CAACAAGTTC ACCTACAACT GCGATGGCCA TACCTTCAAT GACCTTGTCG
2341 AAGATGGATT CAGTAAGTCA CTTTTCGTTT GATCTATGCA TAGGTTTTAA TCTTACACCA
2401 TTCGGTGCTT ACCTCGATTT GTTTCTAGGT TTATTTGCCT AACCTATCGA TTCGTATGGA
2461 TTTGTGCATC CTGAGTAGTC TGATTTCACT GAGGAACCGC ATGAACCAAG CTTTTAGGGT
2521 TAGATCTATA TTTATTAAAC GCAATAAATC TCTGTAGAGT CTTACCTGTA TAATCACTCA
2581 GATTTGGACA GAATCCGTGG TAGCTTTGGT TGATACTATG TGGAAGAACA CCAACATTGA
2641 TCTGAGACAG GGTTTTTACT GGTTGCTTTA TTTGCGTATA GTTTCCTAGG TCATGGGTTT
2701 TGGCTGGCTT CATTTTTCAA TGAAAATCTG GGGTCTTATT GTAAGCTTTT GTGACTATCC
2761 TTTAGTTGTT GTAGCCATAT CAATTTCATA TTAAATCAAT GTCACCTGAT TATCCCGTAA

2821 GGGAAGACAA GGAGATTGAT TTTGTTCCTG TTGTACAGCC TATTGTGTTG TTGCGGTTGA
2881 TTCTGCTGGG AGGCAAATTC CCATGTCCTT TTTGGAAAGA GTAAAAGAAG ATTTTAACAA
2941 GCGATATGGT GGTGGAAAGG CTGCAACTGC TCAAGCAAAC AGCTTGAATA AGGAGTTTGG
3001 GTACTTTTTC ATTATCTCTT CTATTTGGAT GATCTTCTCT TTTATATTCG TGGACTGACT
3061 CTTTTTGGTA ACTTTGAAGC TCTAAACTGA AAGAGCATAT GCAGTATTGC ATGGATCATC
3121 CTGATGAGAT TAGCAAGCTT GCTAAGGTGA AGGCGCAAGT GTCAGAAGTT AAGGGTGTAA
3181 TGATGGAAAA CATTGAGAAG GTTTGAATCT GACCCTTTCT GTCCTCAATG TATATATTTA
3241 CATCTATGGT TGACCCATCT GAAAGAGCTC ATCTAGTCAT ACTAAGTTAC TGTGAAATCA
3301 ATTACTAATA ATTCAATGCA TTACATCTTA CGGGAAATGT CCAGTTTACC AATGTAACAC
3361 CGCTTACATA TGCGACTCAT TATTTGCAGG TTCTTGACCG TGGTGAGAAA ATTGAGCTTT
3421 TGGTGGACAA AACCGAAAAT CTTCGCTCAC AGGTTAGACT CTCTCATACC CTTATCTCCT
3481 GCATCTATTT GCCTACACTC ACACACAAAG ATGAACATGC TTTTTGAGTT CATAGCTGAC
3541 TCATCCTATT ACCATATCTT GTGTCTAACT GAACTCAAAC TGCCACAGGC ACAAGATTTC
3601 AGAACAACAG GAACGCAGAT GAGAAGAAAG ATGTGGCTTC AGAACATGAA GATAAAACTC
3661 ATAGTGCTCG CCATCATTAT CGCACTGATT CTCATCATCG TGCTCTCAGT TTGCCATGGG
3721 TTTAAGTGTT AAGCCCAGAA AATTCAAAAC TTCTGAGATC TTTCTCACTC GGTACATCGT
3781 AGTGCCTTCT CTTTTATTGT TCGATCTTCA ATCTAATGTT TCTGCTTGAA TTGGTTTTGT
3841 ATAGATAGAC ATATATGTAT GCTACTCATT TGCTATATTG CTTGCATCTG ATGACATTGA
3901 TCCCTTGCGA ACCAGTCAGT ACTTGACAAG AGATCCGAAA CTATTAAATG ATTTTGTTGA
3961 CTGAAGATTG CATTGTGGTG GTGATGATAA AAATTTGAGA TTGTATTGAC GATTATACGG
4021 CGTGATGTGT GCTGGTTACT TTAAATTTAG ATAAGAGTAG TGCCCTCTCT ATATCGTTAC
4081 TAGGGATGGT AGTACATATA TATACGTACG TCATATTCCG ACCGTTTTAA GCGGCCATGC
4141 TAGAGTCCGC AAAAATCACC AGTCTCTCTC TACAAATCTA TCTCTCTCTA TTTTTCTCCA
4201 GAATAATGTG TGAGTAGTTC CCAGATAAGG GAATTAGGGT TCTTATAGGG TTTCGCTCAT
4261 GTGTTGAGCA TATAAGAAAC CCTTAGTATG TATTTGTATT TGTAAAATAC TTCTATCAAT
4321 AAAATTTCTA ATTCCTAAAA CCAAAATCCA GTGACACGTG CGGAGCTCCG TACGTAGATC
4381 ATGAGCGGAG AATTAAGGGA GTCACGTTAT GACCCCCGCC GATGACGCGG GACAAGCCGT
4441 TTTACGTTTG GAACTGACAG AACCGCAACG TTGAAGGAGC CACTGAGCGC GGGTTTCTGG
4501 AGTTTAATGA GCTAAGCACA TACGTCAGAA ACCATTATTG CGCGTTCAAA AGTCGCCTAA
4561 GGTCACTATC ATCTAGCAAA TATTTCTTGT CAAAAATGCT CCACTGACGT TCCATAAATT
4621 CCCCTCGGTA TCCAATTAGA GTCTCATATT CACTCTCAAC TCGATCGAGG GGATCTACCA
4681 TGAGTCCAGA AAGGAGACCA GCAGATATTA GGAGAGCAAC CGAAGCAGAT ATGCCAGCAG
4741 TTTGCACCAT TGTGAACCAT TACATCGAGA CTTCTACAGT TAATTTTAGG ACTGAACCTC
4801 AAGAGCCACA GGAATGGACA GATGATCTCG TGAGATTAAG GGAAAGATAC CCTTGGCTTG
4861 TTGCTGAGGT GGATGGAGAA GTTGCAGGTA TTGCTTACGC AGGACCTTGG AAGGCTAGAA
4921 ACGCTTATGA TTGGACTGCT GAGTCTACCG TTTACGTGTC ACCTAGACAT CAAAGAACCG
4981 GACTTGGTTC AACCTTGTAT ACTCACCTTT TGAAGTCTCT TGAAGCTCAG GGATTCAAAT
5041 CTGTTGTGGC TGTTATCGGT TTGCCTAATG ATCCAAGTGT GAGAATGCAT GAAGCTCTCG
5101 GATACGCACC AAGGGGTATG TTAAGAGCTG CTGGATTTAA ACATGGTAAC TGGCACGATG
5161 TTGGTTTCTG GCAGTTAGAT TTCAGTTTAC CAGTTCCACC AAGACCAGTG CTTCCAGTGA
5221 CCGAGATTTG ACCCGGGCTA GAGTCAAGCA GATCGTTCAA ACATTTGGCA ATAAAGTTTC
5281 TTAATATTGA ATCCTGTTGC CGGTCTTGCG ATGATTATCA TATAATTTCT GTTGAATTAC
5341 GTTAAGCATG TAATAATTAA CATGTAATGC ATGACGTTAT TTATGAGATG GGTTTTTATG
5401 ATTAGAGTCC CGCAATTATA CATTTAATAC GCGATAGAAA ACAAAATATA GCGCGCAAAC
5461 TAGGATAAAT TATCGCGCGC GGTGTCATCT ATGTTACTAG ATCGACCGGC ATGAAGCTGA
5521 GTTAACGATG TAAGTCCTCA ATTCGGCGTT AATTCAGTAC ATTAAAAACG TCCGCAATGT
5581 GTTATTAAGT TGTCTAAGCG TCAATTTGTT TACACCACAA TATATCCTGC CACCAGCCAG
5641 CCAACAGCTC CCCGACCGGC AGCTCGGCAC AAAATCACCA CTCGATACAG GCAGCCCATC
5701 AGTCCGGGAC GGCGTCAGCG GGAGAGCCGT TGTAAGGCGG CAGACTTTGC TCATGTTACC
5761 GATGCTATTC GGAAGAACGG CAACTAAGCT GCCGGGTTTG AAACACGGAT GATCTCGCGG
5821 AGGGTAGCAT GTTGATTGTA ACGATGACAG AGCGTTGCTG CCTGTGATCA ATTCGGGCAC
5881 GAACCCAGTG GACATAAGCC TGTTCGGTTC GTAAGCTGTA ATGCAAGTAG CGTATGCGCT
5941 CACGCAACTG GTCCAGAACC TTGACCGAAC GCAGCGGTGG TAACGGCGCA GTGGCGGTTT

6001 TCATGGCTTG TTATGACTGT TTTTTTGGGG TACAGTCTAT GCCTCGGGCA TCCAAGCAGC
6061 AAGCGCGTTA CGCCGTGGGT CGATGTTTGA TGTTATGGAG CAGCAACGAT GTTACGCAGC
6121 AGGGCAGTCG CCCTAAAACA AAGTTAAACA TCATGGGGGA AGCGGTGATC GCCGAAGTAT
6181 CGACTCAACT ATCAGAGGTA GTTGGCGTCA TCGAGCGCCA TCTCGAACCG ACGTTGCTGG
6241 CCGTACATTT GTACGGCTCC GCAGTGGATG GCGGCCTGAA GCCACACAGT GATATTGATT
6301 TGCTGGTTAC GGTGACCGTA AGGCTTGATG AAACAACGCG GCGAGCTTTG ATCAACGACC
6361 TTTTGGAAAC TTCGGCTTCC CCTGGAGAGA GCGAGATTCT CCGCGCTGTA GAAGTCACCA
6421 TTGTTGTGCA CGACGACATC ATTCCGTGGC GTTATCCAGC TAAGCGCGAA CTGCAATTTG
6481 GAGAATGGCA GCGCAATGAC ATTCTTGCAG GTATCTTCGA GCCAGCCACG ATCGACATTG
6541 ATCTGGCTAT CTTGCTGACA AAAGCAAGAG AACATAGCGT TGCCTTGGTA GGTCCAGCGG
6601 CGGAGGAACT CTTTGATCCG GTTCCTGAAC AGGATCTATT TGAGGCGCTA AATGAAACCT
6661 TAACGCTATG GAACTCGCCG CCCGACTGGG CTGGCGATGA GCGAAATGTA GTGCTTACGT
6721 TGTCCCGCAT TTGGTACAGC GCAGTAACCG GCAAAATCGC GCCGAAGGAT GTCGCTGCCG
6781 ACTGGGCAAT GGAGCGCCTG CCGGCCCAGT ATCAGCCCGT CATACTTGAA GCTAGACAGG
6841 CTTATCTTGG ACAAGAAGAA GATCGCTTGG CCTCGCGCGC AGATCAGTTG GAAGAATTTG
6901 TCCACTACGT GAAAGGCGAG ATCACCAAGG TAGTCGGCAA ATAATGTCTA GCTAGAAATT
6961 CGTTCAAGCC GACGCCGCTT CGCGGCGCGG CTTAACTCAA GCGTTAGATG CACTAAGCAC
7021 ATAATTGCTC ACAGCCAAAC TATCAGGTCA AGTCTGCTTT TATTATTTTT AAGCGTGCAT
7081 AATAAGCCCT ACACAAATTG GGAGATATAT CATGCATGAC CAAAATCCCT TAACGTGAGT
7141 TTTCGTTCCA CTGAGCGTCA GACCCCGTAG AAAAGATCAA AGGATCTTCT TGAGATCCTT
7201 TTTTTCTGCG CGTAATCTGC TGCTTGCAAA CAAAAAAACC ACCGCTACCA GCGGTGGTTT
7261 GTTTGCCGGA TCAAGAGCTA CCAACTCTTT TTCCGAAGGT AACTGGCTTC AGCAGAGCGC
7321 AGATACCAAA TACTGTCCTT CTAGTGTAGC CGTAGTTAGG CCACCACTTC AAGAACTCTG
7381 TAGCACCGCC TACATACCTC GCTCTGCTAA TCCTGTTACC AGTGGCTGCT GCCAGTGGCG
7441 ATAAGTCGTG TCTTACCGGG TTGGACTCAA GACGATAGTT ACCGGATAAG GCGCAGCGGT
7501 CGGGCTGAAC GGGGGGTTCG TGCACACAGC CCAGCTTGGA GCGAACGACC TACACCGAAC
7561 TGAGATACCT ACAGCGTGAG CTATGAGAAA GCGCCACGCT TCCCGAAGGG AGAAAGGCGG
7621 ACAGGTATCC GGTAAGCGGC AGGGTCGGAA CAGGAGAGCG CACGAGGGAG CTTCCAGGGG
7681 GAAACGCCTG GTATCTTTAT AGTCCTGTCG GGTTTCGCCA CCTCTGACTT GAGCGTCGAT
7741 TTTTGTGATG CTCGTCAGGG GGGCGGAGCC TATGGAAAAA CGCCAGCAAC GCGGCCTTTT
7801 TACGGTTCCT GGCCTTTTGC TGGCCTTTTG CTCACATGTT CTTTCCTGCG TTATCCCCTG
7861 ATTCTGTGGA TAACCGTATT ACCGCCTTTG AGTGAGCTGA TACCGCTCGC CGCAGCCGAA
7921 CGACCGAGCG CAGCGAGTCA GTGAGCGAGG AAGCGGAAGA GCGCCTGATG CGGTATTTTC
7981 TCCTTACGCA TCTGTGCGGT ATTTCACACC GCATATGGTG CACTCTCAGT ACAATCTGCT
8041 CTGATGCCGC ATAGTTAAGC CAGTATACAC TCCGCTATCG CTACGTGACT GGGTCATGGC
8101 TGCGCCCCGA CACCCGCCAA CACCCGCTGA CGCGCCCTGA CGGGCTTGTC TGCTCCCGGC
8161 ATCCGCTTAC AGACAAGCTG TGACCGTCTC CGGGAGCTGC ATGTGTCAGA GGTTTTCACC
8221 GTCATCACCG AAACGCGCGA GGCAGGGTGC CTTGATGTGG GCGCCGGCGG TCGAGTGGCG
8281 ACGGCGCGGC TTGTCCGCGC CCTGGTAGAT TGCCTGGCCG TAGGCCAGCC ATTTTTGAGC
8341 GGCCAGCGGC CGCGATAGGC CGACGCGAAG CGGCGGGGCG TAGGGAGCGC AGCGACCGAA
8401 GGGTAGGCGC TTTTTGCAGC TCTTCGGCTG TGCGCTGGCC AGACAGTTAT GCACAGGCCA
8461 GGCGGGTTTT AAGAGTTTTA ATAAGTTTTA AAGAGTTTTA GGCGGAAAAA TCGCCTTTTT
8521 TCTCTTTTAT ATCAGTCACT TACATGTGTG ACCGGTTCCC AATGTACGGC TTTGGGTTCC
8581 CAATGTACGG GTTCCGGTTC CCAATGTACG GCTTTGGGTT CCCAATGTAC GTGCTATCCA
8641 CAGGAAAGAG ACCTTTTCGA CCTTTTTCCC CTGCTAGGGC AATTTGCCCT AGCATCTGCT
8701 CCGTACATTA GGAACCGGCG GATGCTTCGC CCTCGATCAG GTTGCGGTAG CGCATGACTA
8761 GGATCGGGCC AGCCTGCCCC GCCTCCTCCT TCAAATCGTA CTCCGGCAGG TCATTTGACC
8821 CGATCAGCTT GCGCACGGTG AAACAGAACT TCTTGAACTC TCCGGCGCTG CCACTGCGTT
8881 CGTAGATCGT CTTGAACAAC CATCTGGCTT CTGCCTTGCC TGCGGCGCGG CGTGCCAGGC
8941 GGTAGAGAAA ACGGCCGATG CCGGGATCGA TCAAAAAGTA ATCGGGGTGA ACCGTCAGCA
9001 CGTCCGGGTT CTTGCCTTCT GTGATCTCGC GGTACATCCA ATCAGCTAGC TCGATCTCGA
9061 TGTACTCCGG CCGCCCGGTT TCGCTCTTTA CGATCTTGTA GCGGCTAATC AAGGCTTCAC
9121 CCTCGGATAC CGTCACCAGG CGGCCGTTCT TGGCCTTCTT CGTACGCTGC ATGGCAACGT

9181 GCGTGGTGTT TAACCGAATG CAGGTTTCTA CCAGGTCGTC TTTCTGCTTT CCGCCATCGG
9241 CTCGCCGGCA GAACTTGAGT ACGTCCGCAA CGTGTGGACG GAACACGCGG CCGGGCTTGT
9301 CTCCCTTCCC TTCCCGGTAT CGGTTCATGG ATTCGGTTAG ATGGGAAACC GCCATCAGTA
9361 CCAGGTCGTA ATCCCACACA CTGGCCATGC CGGCCGGCCC TGCGGAAACC TCTACGTGCC
9421 CGTCTGGAAG CTCGTAGCGG ATCACCTCGC CAGCTCGTCG GTCACGCTTC GACAGACGGA
9481 AAACGGCCAC GTCCATGATG CTGCGACTAT CGCGGGTGCC CACGTCATAG AGCATCGGAA
9541 CGAAAAAATC TGGTTGCTCG TCGCCCTTGG GCGGCTTCCT AATCGACGGC GCACCGGCTG
9601 CCGGCGGTTG CCGGGATTCT TTGCGGATTC GATCAGCGGC CGCTTGCCAC GATTCACCGG
9661 GGCGTGCTTC TGCCTCGATG CGTTGCCGCT GGGCGGCCTG CGCGGCCTTC AACTTCTCCA
9721 CCAGGTCATC ACCCAGCGCC GCGCCGATTT GTACCGGGCC GGATGGTTTG CGACCGCTCA
9781 CGCCGATTCC TCGGGCTTGG GGGTTCCAGT GCCATTGCAG GGCCGGCAGA CAACCCAGCC
9841 GCTTACGCCT GGCCAACCGC CCGTTCCTCC ACACATGGGG CATTCCACGG CGTCGGTGCC
9901 TGGTTGTTCT TGATTTTCCA TGCCGCCTCC TTTAGCCGCT AAAATTCATC TACTCATTTA
9961 TTCATTTGCT CATTTACTCT GGTAGCTGCG CGATGTATTC AGATAGCAGC TCGGTAATGG
10021 TCTTGCCTTG GCGTACCGCG TACATCTTCA GCTTGGTGTG ATCCTCCGCC GGCAACTGAA
10081 AGTTGACCCG CTTCATGGCT GGCGTGTCTG CCAGGCTGGC CAACGTTGCA GCCTTGCTGC
10141 TGCGTGCGCT CGGACGGCCG GCACTTAGCG TGTTTGTGCT TTTGCTCATT TTCTCTTTAC
10201 CTCATTAACT CAAATGAGTT TTGATTTAAT TTCAGCGGCC AGCGCCTGGA CCTCGCGGGC
10261 AGCGTCGCCC TCGGGTTCTG ATTCAAGAAC GGTTGTGCCG GCGGCGGCAG TGCCTGGGTA
10321 GCTCACGCGC TGCGTGATAC GGGACTCAAG AATGGGCAGC TCGTACCCGG CCAGCGCCTC
10381 GGCAACCTCA CCGCCGATGC GCGTGCCTTT GATCGCCCGC GACACGACAA AGGCCGCTTG
10441 TAGCCTTCCA TCCGTGACCT CAATGCGCTG CTTAACCAGC TCCACCAGGT CGGCGGTGGC
10501 CCATATGTCG TAAGGGCTTG GCTGCACCGG AATCAGCACG AAGTCGGCTG CCTTGATCGC
10561 GGACACAGCC AAGTCCGCCG CCTGGGGCGC TCCGTCGATC ACTACGAAGT CGCGCCGGCC
10621 GATGGCCTTC ACGTCGCGGT CAATCGTCGG GCGGTCGATG CCGACAACGG TTAGCGGTTG
10681 ATCTTCCCGC ACGGCCGCCC AATCGCGGGC ACTGCCCTGG GGATCGGAAT CGACTAACAG
10741 AACATCGGCC CCGGCGAGTT GCAGGGCGCG GGCTAGATGG GTTGCGATGG TCGTCTTGCC
10801 TGACCCGCCT TTCTGGTTAA GTACAGCGAT AACCTTCATG CGTTCCCCTT GCGTATTTGT
10861 TTATTTACTC ATCGCATCAT ATACGCAGCG ACCGCATGAC GCAAGCTGTT TTACTCAAAT
10921 ACACATCACC TTTTTAGACG GCGGCGCTCG GTTTCTTCAG CGGCCAAGCT GGCCGGCCAG
10981 GCCGCCAGCT TGGCATCAGA CAAACCGGCC AGGATTTCAT GCAGCCGCAC GGTTGAGACG
11041 TGCGCGGGCG GCTCGAACAC GTACCCGGCC GCGATCATCT CCGCCTCGAT CTCTTCGGTA
11101 ATGAAAAACG GTTCGTCCTG GCCGTCCTGG TGCGGTTTCA TGCTTGTTCC TCTTGGCGTT
11161 CATTCTCGGC GGCCGCCAGG GCGTCGGCCT CGGTCAATGC GTCCTCACGG AAGGCACCGC
11221 GCCGCCTGGC CTCGGTGGGC GTCACTTCCT CGCTGCGCTC AAGTGCGCGG TACAGGGTCG
11281 AGCGATGCAC GCCAAGCAGT GCAGCCGCCT CTTTCACGGT GCGGCCTTCC TGGTCGATCA
11341 GCTCGCGGGC GTGCGCGATC TGTGCCGGGG TGAGGGTAGG GCGGGGGCCA AACTTCACGC
11401 CTCGGGCCTT GGCGGCCTCG CGCCCGCTCC GGGTGCGGTC GATGATTAGG GAACGCTCGA
11461 ACTCGGCAAT GCCGGCGAAC ACGGTCAACA CCATGCGGCC GGCCGGCGTG GTGGTGTCGG
11521 CCCACGGCTC TGCCAGGCTA CGCAGGCCCG CGCCGGCCTC CTGGATGCGC TCGGCAATGT
11581 CCAGTAGGTC GCGGGTGCTG CGGGCCAGGC GGTCTAGCCT GGTCACTGTC ACAACGTCGC
11641 CAGGGCGTAG GTGGTCAAGC ATCCTGGCCA GCTCCGGGCG GTCGCGCCTG GTGCCGGTGA
11701 TCTTCTCGGA AAACAGCTTG GTGCAGCCGG CCGCGTGCAG TTCGGCCCGT TGGTTGGTCA
11761 AGTCCTGGTC GTCGGTGCTG ACGCGGGCAT AGCCCAGCAG GCCAGCGGCG GCGCTCTTGT
11821 TCATGGCGTA ATGTCTCCGG TTCTAGTCGC AAGTATTCTA CTTTATGCGA CTAAAACACG
11881 CGACAAGAAA ACGCCAGGAA AAGGGCAGGG CGGCAGCCTG TCGCGTAACT TAGGACTTGT
11941 GCGACATGTC GTTTTCAGAA GACGGCTGCA CTGAACGTCA GAAGCCGACT GCACTATAGC
12001 AGCGGAGGGG TTGGATCAAA GTACTTTAAA GTACTTTAAA GTACTTTAAA GTACTTTGAT
12061 CCCGAGGGGA ACCCTGTGGT TGGCATGCAC ATACAAATGG ACGAACGGAT AAACCTTTTC
12121 ACGCCCTTTT AAATATCCGA TTATTCTAAT AAACGCTCTT TTCTCTTAGG TTTACCCGCC
12181 AATATATCCT GTCAAACACT GATAGTTT
//
LOCUS D610_pVAMP721_gVAMP723 12102 bp DNA circular UNA 02-JAN-2019

```
COMMENT     ApEinfo:methylated:1
FEATURES             Location/Qualifiers
     misc_feature    join(11949..12102,1..42)
                     /vntifkey="21"
                     /locus_tag="RB"
     promoter        68..1400
                     /locus_tag="Promotor Vamp721"
     CDS             1407..1409
                     /locus_tag="MYC(1)"
     CDS             1410..2126
                     /locus_tag=""
                     /label="XhoI-meGFP-SalI"
     CDS             2130..2159
                     /locus_tag="MYC"
     exon            2163..2352
                     /label="Exon 1"
     gene            2163..4017
                     /locus_tag="gVamp723"
     mutation        2740
                     /label="C for T change"
     exon            2888..3031
                     /label="Exon 2"
     exon            3118..3234
                     /label="Exon 3"
     misc_feature    3230..3251
                     /locus_tag="g723 seq primer"
     exon            3434..3480
                     /label="Exon 4"
     exon            3603..3758
                     /label="Exon 5"
     terminator      4043..4248
                     /locus_tag="T35S"
     promoter        4271..4573
                     /vntifkey="29"
                     /locus_tag="nosP"
     misc_feature    4273..4278
                     /locus_tag="BspHI site"
     CDS             4574..5125
                     /vntifkey="4"
                     /locus_tag="BarCO"
     terminator      5126..5414
                     /vntifkey="43"
                     /locus_tag="nosT"
     misc_feature    complement(5435..5764)
                     /vntifkey="21"
                     /locus_tag="LB"
     CDS             5770..7014
                     /vntifkey="4"
                     /locus_tag="SmR"
     rep_origin      7288..7968
                     /vntifkey="33"
                     /locus_tag="pBR322 ori"
     rep_origin      8378..10972
```

/vntifkey="33"
/locus_tag="pVS1 ori"
ORIGIN
1 AAACTGAAGG CGGGAAACGA CAATCTGATC CAAGCTCAAG CTAAGCTAGG ACTAGCGCTC
61 GCACGTGTCT TCCTTTTATC CAGATGAGAG AAAGATAAAA AAACGAAACC TAAGAACACC
121 ACAAAAATCC GATCGAATTG ATGAGAGAGA TTAGTAAGCG ATGTAAAAAT ACACAAGGGA
181 ACGAGGGGTA GCGATGGAGA AAGCAGGACG CAGAGATCGG AGAAGAGAAA CTATATGAGA
241 GCGCTGGATT TGCGCGAGTA ACAGAGACAT TATTTTCCCT TTGGTAACGT TCAAATTTTG
301 AATTTTTAAA TATAAAAAGT AAATAAAAAG AAAAATTCAT CATTTGTATA GTCGAAAATT
361 GCCTTAATAA ACTTTATTTT GGTAAATAAT TCACGTCTAA TTAATAATAG TATCTTCATG
421 TTTTTACACC ACGGTGGCTC GTTTTTCAAT TCAAATGTGA AATGTTTATC AATGAGCATT
481 TTAAAAAACA AATATATATA TATATATATA TATTATATAT TGATTTTTTA TAAAGAAAAT
541 TGCGATGTGT AAGGGATATT TGAGTTGTTA AATTTTCTAT AAATTATATG TTATGTTACT
601 TAAAAGAACG TTCAATTGAT AATGTTGCAA ACCTTCCTAC CAAAAAAATT AAAATAATGT
661 TGCAAACCTT TTCTTTCATT CCAATTCACT AACCTTCTAT TCTAACCACA ACCTAACTAC
721 CTAAATTTCA ATTATTTGAT TGTTGATCAA AATCTTTAGG AACACGTTAA ACAAATTAAT
781 TAGCAGGTTC TAAACAATGC AGAGCAATTG TAGATGACAA GAGATAACGG TAGAAGTTCG
841 AATTTTGTTG CTGTAACAAG ATACTATGAC TATGAGCTCC CCCCGTCCAT TAAGAATTAA
901 GAAATTTGGT TAATTACAAC ATTTGTTTTT ATGTTTAAGC CTTTTGTTCG TAACTGAGCT
961 AATACTGGAA TCACTGATCA ATGATGAATT TGGAAAAAAA GATATTTTGA GGAAGACAGA
1021 GACAGACTCT CTTCGACCAT GGGACCTGAA CTTGTCCTAA AGTTGGTAAT TACTAATTTG
1081 TGCCATCGGC AGAGTCCACG AGCTTTTTGG CTGGATCCGG GCGGGTCCAA CTTTCTCGAC
1141 ATTTGTTTGA TCTCTCCGTT TTGAAATTGT TTTTTTTTAG TCCTTAAATT CAAGGTTTTA
1201 TACTATTTTT GACCCAATCT GACTACCTTA AATGACAACT TTACCCTTGC CCGCTTTTCA
1261 ACTTTTTCTT CTCCCATGTC CACTTTAGTC CTCGTTTTGT TCTTTCATAG TCTTCTAGAT
1321 CTGTAGTTTT TTTCCGCGAC CGCCCAAAAA ATTCTCTGAT CAGACGGTCG ATTGGATCGG
1381 AGATTTAAGG TAAAGAAAAA ATGCTCGAGA TGGTGAGCAA GGGCGAGGAG CTGTTCACCG
1441 GGGTGGTGCC CATCCTGGTC GAGCTGGACG GCGACGTAAA CGGCCACAAG TTCAGCGTGT
1501 CCGGCGAGGG CGAGGGCGAT GCCACCTACG GCAAGCTGAC CCTGAAGTTC ATCTGCACCA
1561 CCGGCAAGCT GCCCGTGCCC TGGCCCACCC TCGTGACCAC CCTGACCTAC GGCGTGCAGT
1621 GCTTCAGCCG CTACCCCGAC CACATGAAGC AGCACGACTT CTTCAAGTCC GCCATGCCCG
1681 AAGGCTACGT CCAGGAGCGC ACCATCTTCT TCAAGGACGA CGGCAACTAC AAGACCCGCG
1741 CCGAGGTGAA GTTCGAGGGC GACACCCTGG TGAACCGCAT CGAGCTGAAG GGCATCGACT
1801 TCAAGGAGGA CGGCAACATC CTGGGGCACA AGCTGGAGTA CAACTACAAC AGCCACAACG
1861 TCTATATCAT GGCCGACAAG CAGAAGAACG GCATCAAGGT GAACTTCAAG ATCCGCCACA
1921 ACATCGAGGA CGGCAGCGTG CAGCTCGCCG ACCACTACCA GCAGAACACC CCCATCGGCG
1981 ACGGCCCCGT GCTGCTGCCC GACAACCACT ACCTGAGTAC TCAGTCCAAG CTGAGCAAAG
2041 ACCCCAACGA GAAGCGCGAT CACATGGTCC TGCTGGAGTT CGTGACCGCC GCCGGGATCA
2101 CTCTCGGCAT GGACGAGCTG TACAAGGTCG AGCAGAAGTT GATCTCAGAG GAGGACTTAG
2161 GCATGGCGCA ACAATCGTTG TTCTACAGTT TCATCGCTCG CGGCACCGTA ATCCTCGTCG
2221 AGTTCACAGA TTTCAAAGGC AATTTCACAT CTGTCGCTGC TCAGTACCTT GAGAATCTTC
2281 CTTCCTCGAA CAACAAGTTT ACCTACAACT GCGATGGTCA TACGTTCAAC GACCTCGTCG
2341 AAAATGGATT CAGTGAGTCA AAATATTGCT CGTGATGTGT TTGTGATTGT GTTTTCGATT
2401 AGTCGATTTA TTCATTGTTT GGATTAGATT TCTTTTTACC TAATTGAGCA TTTGAGAATC
2461 GAGTTCTTAT GTTTGGTATA TCTCTATGAC TTATCTGAGT CGAGTTCTTA TGTGTTTACC
2521 GTTTGGTATA ATTTCAGGAA TAGTTATGTA TTGGGTTTGA TAATTAATTG ATGTTAGATC
2581 TATAATCAGT GATATGTTTG GTGGATCTAT TTAAGCATCT ATGCAATTCT TATTATGAAT
2641 CTATGATTCT TGGCTTCTTT GCGTGTAGCT TTTGTTTTCA TGAATTTGGC TGGTTTTTCA
2701 AATGAGAGTT GTTTGGGAAA TTTGTGGATA TATATCGCTT GAGTAAATTG CATTGGTGTA
2761 TTGTGTTTAT TATATTGCTC TGTTTTAATT AAACTTAGTT GGGTTTTGAT GACGTTGGAT
2821 ATGGTCCATA TCATATGGAC TCTTTATTAT ATTTTTTAAG TTTCTTATGA GGGATATGTT
2881 TGTGCAGCCT ATTGTGTTGT TGCAGTTGAT TCTGCTGGGA GGGAGATTCC TATGGCTTTC
2941 TTGGAACGCG TGAAGGAGGA TTTTTATAAG AGATATGGTG GTGAAAAGGC TGCAACTGAT

3001 CAAGCAAATA GCTTGAATAA AGAATTTGGG TATGTGTTTT TTGATCTATA TGATTGGTAT
3061 TGAGAAGTTT GTATTATTCA TGTCAATGAC AGACTCCTTT TTGGCTGCTC TAAAGGTCGA
3121 ATCTGAAAGA GCACATGCAG TATTGCATGG ATCATCCTGA TGAGATTAGC AACCTTGCTA
3181 AAGCTAAAGC TCAAGTGTCT GAAGTTAAAA GTTTAATGAT GGAAAACATT GAGAAGGTTC
3241 GATTCTGACC CTTTTTCTG TTAATGGTAT AATTTTATGT TTTAATCTAT TTCCAGTAGA
3301 GTTAAGCTCG TATGAGTTAA GTGTCATGAT CTTTTGGTTC CTTAACCTTT AAATTGATGA
3361 GGGGAATGTC CAGTTCTGGG TTGTATAGGC ATGTATGCTG ACCGTTTTTA CATTGTGGAA
3421 CACATTCATT TGCAGGTTCT TGCCCGTGGT GTGATATGTG AGATGCTGGG TAGTTCAGAG
3481 GTTAGTTCTC TTAGGCATAT GGTTCTAAGT TCCATACACA TAGATATGCT TGAGTTGCCA
3541 GTTCTCTCTT ACTAATGTTG TCTTGGGTTT CCAACTTTTA ATTGAACTCG AAACTGTCAC
3601 AGTCACAGCC GCAAGCTTTC TATATAAAAA GAACTCAAAT GAAAAGGAAG AAGTGGTTTC
3661 AGAACATGAA GATAAAACTC ATTGTCCTTG CAATTATCAT TGCCTTGATT CTCATCATCA
3721 TCCTCTCGGT TTGTGGGGGA TTCAACTGCG GTAAATAAGT CTGGAACATT TCTTCCCGGC
3781 GTTATCGACT GCGCTCTGTG CTTCCCAAGA TCTCTGAGAA TATCTTCATT CAGTTCGTTG
3841 TGCTGCTTTC TTTTTGTGGT TTCTAATTTC TAATTTCTGC TATGTCCTTT CTACTTAGGT
3901 TGTACATAAA TATTATACAT ATATGTATGT CCTTTCTTCT TTGAAATTGG TTTTGTATAG
3961 GCCATATATA TGTATGAATG TTTTTGCTAT TACTGGTACT GACACTGATT CTTGCGAGTT
4021 TTAAGCGGCC ATGCTAGAGT CCGCAAAAAT CACCAGTCTC TCTCTACAAA TCTATCTCTC
4081 TCTATTTTTC TCCAGAATAA TGTGTGAGTA GTTCCCAGAT AAGGGAATTA GGGTTCTTAT
4141 AGGGTTTCGC TCATGTGTTG AGCATATAAG AAACCCTTAG TATGTATTTG TATTTGTAAA
4201 ATACTTCTAT CAATAAAATT TCTAATTCCT AAAACCAAAA TCCAGTGACA CGTGCGGAGC
4261 TCCGTACGTA GATCATGAGC GGAGAATTAA GGGAGTCACG TTATGACCCC CGCCGATGAC
4321 GCGGGACAAG CCGTTTTACG TTTGGAACTG ACAGAACCGC AACGTTGAAG GAGCCACTGA
4381 GCGCGGGTTT CTGGAGTTTA ATGAGCTAAG CACATACGTC AGAAACCATT ATTGCGCGTT
4441 CAAAAGTCGC CTAAGGTCAC TATCATCTAG CAAATATTTC TTGTCAAAAA TGCTCCACTG
4501 ACGTTCCATA AATTCCCCTC GGTATCCAAT TAGAGTCTCA TATTCACTCT CAACTCGATC
4561 GAGGGGATCT ACCATGAGTC CAGAAAGGAG ACCAGCAGAT ATTAGGAGAG CAACCGAAGC
4621 AGATATGCCA GCAGTTTGCA CCATTGTGAA CCATTACATC GAGACTTCTA CAGTTAATTT
4681 TAGGACTGAA CCTCAAGAGC CACAGGAATG GACAGATGAT CTCGTGAGAT TAAGGGAAAG
4741 ATACCCTTGG CTTGTTGCTG AGGTGGATGG AGAAGTTGCA GGTATTGCTT ACGCAGGACC
4801 TTGGAAGGCT AGAAACGCTT ATGATTGGAC TGCTGAGTCT ACCGTTTACG TGTCACCTAG
4861 ACATCAAAGA ACCGGACTTG GTTCAACCTT GTATACTCAC CTTTTGAAGT CTCTTGAAGC
4921 TCAGGGATTC AAATCTGTTG TGGCTGTTAT CGGTTTGCCT AATGATCCAA GTGTGAGAAT
4981 GCATGAAGCT CTCGGATACG CACCAAGGGG TATGTTAAGA GCTGCTGGAT TTAAACATGG
5041 TAACTGGCAC GATGTTGGTT TCTGGCAGTT AGATTTCAGT TTACCAGTTC CACCAAGACC
5101 AGTGCTTCCA GTGACCGAGA TTTGACCCGG GCTAGAGTCA AGCAGATCGT TCAAACATTT
5161 GGCAATAAAG TTTCTTAATA TTGAATCCTG TTGCCGGTCT TGCGATGATT ATCATATAAT
5221 TTCTGTTGAA TTACGTTAAG CATGTAATAA TTAACATGTA ATGCATGACG TTATTTATGA
5281 GATGGGTTTT TATGATTAGA GTCCCGCAAT TATACATTTA ATACGCGATA GAAAACAAAA
5341 TATAGCGCGC AAACTAGGAT AAATTATCGC GCGCGGTGTC ATCTATGTTA CTAGATCGAC
5401 CGGCATGAAG CTGAGTTAAC GATGTAAGTC CTCAATTCGG CGTTAATTCA GTACATTAAA
5461 AACGTCCGCA ATGTGTTATT AAGTTGTCTA AGCGTCAATT TGTTTACACC ACAATATATC
5521 CTGCCACCAG CCAGCCAACA GCTCCCCGAC CGGCAGCTCG GCACAAAATC ACCACTCGAT
5581 ACAGGCAGCC CATCAGTCCG GGACGGCGTC AGCGGGAGAG CCGTTGTAAG GCGGCAGACT
5641 TTGCTCATGT TACCGATGCT ATTCGGAAGA ACGGCAACTA AGCTGCCGGG TTTGAAACAC
5701 GGATGATCTC GCGGAGGGTA GCATGTTGAT TGTAACGATG ACAGAGCGTT GCTGCCTGTG
5761 ATCAATTCGG GCACGAACCC AGTGGACATA AGCCTGTTCG GTTCGTAAGC TGTAATGCAA
5821 GTAGCGTATG CGCTCACGCA ACTGGTCCAG AACCTTGACC GAACGCAGCG GTGGTAACGG
5881 CGCAGTGGCG GTTTTCATGG CTTGTTATGA CTGTTTTTTT GGGGTACAGT CTATGCCTCG
5941 GGCATCCAAG CAGCAAGCGC GTTACGCCGT GGGTCGATGT TTGATGTTAT GGAGCAGCAA
6001 CGATGTTACG CAGCAGGGCA GTCGCCCTAA AACAAAGTTA AACATCATGG GGGAAGCGGT
6061 GATCGCCGAA GTATCGACTC AACTATCAGA GGTAGTTGGC GTCATCGAGC GCCATCTCGA
6121 ACCGACGTTG CTGGCCGTAC ATTTGTACGG CTCCGCAGTG GATGGCGGCC TGAAGCCACA

6181 CAGTGATATT GATTTGCTGG TTACGGTGAC CGTAAGGCTT GATGAAACAA CGCGGCGAGC
6241 TTTGATCAAC GACCTTTTGG AAACTTCGGC TTCCCCTGGA GAGAGCGAGA TTCTCCGCGC
6301 TGTAGAAGTC ACCATTGTTG TGCACGACGA CATCATTCCG TGGCGTTATC CAGCTAAGCG
6361 CGAACTGCAA TTTGGAGAAT GGCAGCGCAA TGACATTCTT GCAGGTATCT TCGAGCCAGC
6421 CACGATCGAC ATTGATCTGG CTATCTTGCT GACAAAAGCA AGAGAACATA GCGTTGCCTT
6481 GGTAGGTCCA GCGGCGGAGG AACTCTTTGA TCCGGTTCCT GAACAGGATC TATTTGAGGC
6541 GCTAAATGAA ACCTTAACGC TATGGAACTC GCCGCCCGAC TGGGCTGGCG ATGAGCGAAA
6601 TGTAGTGCTT ACGTTGTCCC GCATTTGGTA CAGCGCAGTA ACCGGCAAAA TCGCGCCGAA
6661 GGATGTCGCT GCCGACTGGG CAATGGAGCG CCTGCCGGCC CAGTATCAGC CCGTCATACT
6721 TGAAGCTAGA CAGGCTTATC TTGGACAAGA AGAAGATCGC TTGGCCTCGC GCGCAGATCA
6781 GTTGGAAGAA TTTGTCCACT ACGTGAAAGG CGAGATCACC AAGGTAGTCG GCAAATAATG
6841 TCTAGCTAGA AATTCGTTCA AGCCGACGCC GCTTCGCGGC GCGGCTTAAC TCAAGCGTTA
6901 GATGCACTAA GCACATAATT GCTCACAGCC AAACTATCAG GTCAAGTCTG CTTTTATTAT
6961 TTTTAAGCGT GCATAATAAG CCCTACACAA ATTGGGAGAT ATATCATGCA TGACCAAAAT
7021 CCCTTAACGT GAGTTTTCGT TCCACTGAGC GTCAGACCCC GTAGAAAAGA TCAAAGGATC
7081 TTCTTGAGAT CCTTTTTTTC TGCGCGTAAT CTGCTGCTTG CAAACAAAAA AACCACCGCT
7141 ACCAGCGGTG GTTTGTTTGC CGGATCAAGA GCTACCAACT CTTTTTCCGA AGGTAACTGG
7201 CTTCAGCAGA GCGCAGATAC CAAATACTGT CCTTCTAGTG TAGCCGTAGT TAGGCCACCA
7261 CTTCAAGAAC TCTGTAGCAC CGCCTACATA CCTCGCTCTG CTAATCCTGT TACCAGTGGC
7321 TGCTGCCAGT GGCGATAAGT CGTGTCTTAC CGGGTTGGAC TCAAGACGAT AGTTACCGGA
7381 TAAGGCGCAG CGGTCGGGCT GAACGGGGGG TTCGTGCACA CAGCCCAGCT TGGAGCGAAC
7441 GACCTACACC GAACTGAGAT ACCTACAGCG TGAGCTATGA GAAAGCGCCA CGCTTCCCGA
7501 AGGGAGAAAG GCGGACAGGT ATCCGGTAAG CGGCAGGGTC GGAACAGGAG AGCGCACGAG
7561 GGAGCTTCCA GGGGGAAACG CCTGGTATCT TTATAGTCCT GTCGGGTTTC GCCACCTCTG
7621 ACTTGAGCGT CGATTTTTGT GATGCTCGTC AGGGGGGCGG AGCCTATGGA AAAACGCCAG
7681 CAACGCGGCC TTTTTACGGT TCCTGGCCTT TTGCTGGCCT TTTGCTCACA TGTTCTTTCC
7741 TGCGTTATCC CCTGATTCTG TGGATAACCG TATTACCGCC TTTGAGTGAG CTGATACCGC
7801 TCGCCGCAGC CGAACGACCG AGCGCAGCGA GTCAGTGAGC GAGGAAGCGG AAGAGCGCCT
7861 GATGCGGTAT TTTCTCCTTA CGCATCTGTG CGGTATTTCA CACCGCATAT GGTGCACTCT
7921 CAGTACAATC TGCTCTGATG CCGCATAGTT AAGCCAGTAT ACACTCCGCT ATCGCTACGT
7981 GACTGGGTCA TGGCTGCGCC CCGACACCCG CCAACACCCG CTGACGCGCC CTGACGGGCT
8041 TGTCTGCTCC CGGCATCCGC TTACAGACAA GCTGTGACCG TCTCCGGGAG CTGCATGTGT
8101 CAGAGGTTTT CACCGTCATC ACCGAAACGC GCGAGGCAGG GTGCCTTGAT GTGGGCGCCG
8161 GCGGTCGAGT GGCGACGGCG CGGCTTGTCC GCGCCCTGGT AGATTGCCTG GCCGTAGGCC
8221 AGCCATTTTT GAGCGGCCAG CGGCCGCGAT AGGCCGACGC GAAGCGGCGG GGCGTAGGGA
8281 GCGCAGCGAC CGAAGGGTAG GCGCTTTTTG CAGCTCTTCG GCTGTGCGCT GGCCAGACAG
8341 TTATGCACAG GCCAGGCGGG TTTTAAGAGT TTTAATAAGT TTTAAAGAGT TTTAGGCGGA
8401 AAAATCGCCT TTTTTCTCTT TTATATCAGT CACTTACATG TGTGACCGGT TCCCAATGTA
8461 CGGCTTTGGG TTCCCAATGT ACGGGTTCCG GTTCCCAATG TACGGCTTTG GGTTCCCAAT
8521 GTACGTGCTA TCCACAGGAA AGAGACCTTT TCGACCTTTT TCCCCTGCTA GGGCAATTTG
8581 CCCTAGCATC TGCTCCGTAC ATTAGGAACC GGCGGATGCT TCGCCCTCGA TCAGGTTGCG
8641 GTAGCGCATG ACTAGGATCG GGCCAGCCTG CCCCGCCTCC TCCTTCAAAT CGTACTCCGG
8701 CAGGTCATTT GACCCGATCA GCTTGCGCAC GGTGAAACAG AACTTCTTGA ACTCTCCGGC
8761 GCTGCCACTG CGTTCGTAGA TCGTCTTGAA CAACCATCTG GCTTCTGCCT TGCCTGCGGC
8821 GCGGCGTGCC AGGCGGTAGA GAAAACGGCC GATGCCGGGA TCGATCAAAA AGTAATCGGG
8881 GTGAACCGTC AGCACGTCCG GGTTCTTGCC TTCTGTGATC TCGCGGTACA TCCAATCAGC
8941 TAGCTCGATC TCGATGTACT CCGGCCGCCC GGTTTCGCTC TTTACGATCT TGTAGCGGCT
9001 AATCAAGGCT TCACCCTCGG ATACCGTCAC CAGGCGGCCG TTCTTGGCCT TCTTCGTACG
9061 CTGCATGGCA ACGTGCGTGG TGTTTAACCG AATGCAGGTT TCTACCAGGT CGTCTTTCTG
9121 CTTTCCGCCA TCGGCTCGCC GGCAGAACTT GAGTACGTCC GCAACGTGTG GACGGAACAC
9181 GCGGCCGGGC TTGTCTCCCT TCCCTTCCCG GTATCGGTTC ATGGATTCGG TTAGATGGGA
9241 AACCGCCATC AGTACCAGGT CGTAATCCCA CACACTGGCC ATGCCGGCCG GCCCTGCGGA
9301 AACCTCTACG TGCCCGTCTG GAAGCTCGTA GCGGATCACC TCGCCAGCTC GTCGGTCACG

9361 CTTCGACAGA CGGAAAACGG CCACGTCCAT GATGCTGCGA CTATCGCGGG TGCCCACGTC
9421 ATAGAGCATC GGAACGAAAA AATCTGGTTG CTCGTCGCCC TTGGGCGGCT TCCTAATCGA
9481 CGGCGCACCG GCTGCCGGCG GTTGCCGGGA TTCTTTGCGG ATTCGATCAG CGGCCGCTTG
9541 CCACGATTCA CCGGGGCGTG CTTCTGCCTC GATGCGTTGC CGCTGGGCGG CCTGCGCGGC
9601 CTTCAACTTC TCCACCAGGT CATCACCCAG CGCCGCGCCG ATTTGTACCG GGCCGGATGG
9661 TTTGCGACCG CTCACGCCGA TTCCTCGGGC TTGGGGGTTC CAGTGCCATT GCAGGGCCGG
9721 CAGACAACCC AGCCGCTTAC GCCTGGCCAA CCGCCCGTTC CTCCACACAT GGGGCATTCC
9781 ACGGCGTCGG TGCCTGGTTG TTCTTGATTT TCCATGCCGC CTCCTTTAGC CGCTAAAATT
9841 CATCTACTCA TTTATTCATT TGCTCATTTA CTCTGGTAGC TGCGCGATGT ATTCAGATAG
9901 CAGCTCGGTA ATGGTCTTGC CTTGGCGTAC CGCGTACATC TTCAGCTTGG TGTGATCCTC
9961 CGCCGGCAAC TGAAAGTTGA CCCGCTTCAT GGCTGGCGTG TCTGCCAGGC TGGCCAACGT
10021 TGCAGCCTTG CTGCTGCGTG CGCTCGGACG GCCGGCACTT AGCGTGTTTG TGCTTTTGCT
10081 CATTTTCTCT TTACCTCATT AACTCAAATG AGTTTTGATT TAATTTCAGC GGCCAGCGCC
10141 TGGACCTCGC GGGCAGCGTC GCCCTCGGGT TCTGATTCAA GAACGGTTGT GCCGGCGGCG
10201 GCAGTGCCTG GGTAGCTCAC GCGCTGCGTG ATACGGGACT CAAGAATGGG CAGCTCGTAC
10261 CCGGCCAGCG CCTCGGCAAC CTCACCGCCG ATGCGCGTGC CTTTGATCGC CCGCGACACG
10321 ACAAAGGCCG CTTGTAGCCT TCCATCCGTG ACCTCAATGC GCTGCTTAAC CAGCTCCACC
10381 AGGTCGGCGG TGGCCCATAT GTCGTAAGGG CTTGGCTGCA CCGGAATCAG CACGAAGTCG
10441 GCTGCCTTGA TCGCGGACAC AGCCAAGTCC GCCGCCTGGG GCGCTCCGTC GATCACTACG
10501 AAGTCGCGCC GGCCGATGGC CTTCACGTCG CGGTCAATCG TCGGGCGGTC GATGCCGACA
10561 ACGGTTAGCG GTTGATCTTC CCGCACGGCC GCCCAATCGC GGGCACTGCC CTGGGGATCG
10621 GAATCGACTA ACAGAACATC GGCCCCGGCG AGTTGCAGGG CGCGGGCTAG ATGGGTTGCG
10681 ATGGTCGTCT TGCCTGACCC GCCTTTCTGG TTAAGTACAG CGATAACCTT CATGCGTTCC
10741 CCTTGCGTAT TTGTTTATTT ACTCATCGCA TCATATACGC AGCGACCGCA TGACGCAAGC
10801 TGTTTTACTC AAATACACAT CACCTTTTTA GACGGCGGCG CTCGGTTTCT TCAGCGGCCA
10861 AGCTGGCCGG CCAGGCCGCC AGCTTGGCAT CAGACAAACC GGCCAGGATT TCATGCAGCC
10921 GCACGGTTGA GACGTGCGCG GGCGGCTCGA ACACGTACCC GGCCGCGATC ATCTCCGCCT
10981 CGATCTCTTC GGTAATGAAA AACGGTTCGT CCTGGCCGTC CTGGTGCGGT TTCATGCTTG
11041 TTCCTCTTGG CGTTCATTCT CGGCGGCCGC CAGGGCGTCG GCCTCGGTCA ATGCGTCCTC
11101 ACGGAAGGCA CCGCGCCGCC TGGCCTCGGT GGGCGTCACT TCCTCGCTGC GCTCAAGTGC
11161 GCGGTACAGG GTCGAGCGAT GCACGCCAAG CAGTGCAGCC GCCTCTTTCA CGGTGCGGCC
11221 TTCCTGGTCG ATCAGCTCGC GGGCGTGCGC GATCTGTGCC GGGGTGAGGG TAGGGCGGGG
11281 GCCAAACTTC ACGCCTCGGG CCTTGGCGGC CTCGCGCCCG CTCCGGGTGC GGTCGATGAT
11341 TAGGGAACGC TCGAACTCGG CAATGCCGGC GAACACGGTC AACACCATGC GGCCGGCCGG
11401 CGTGGTGGTG TCGGCCCACG GCTCTGCCAG GCTACGCAGG CCCGCGCCGG CCTCCTGGAT
11461 GCGCTCGGCA ATGTCCAGTA GGTCGCGGGT GCTGCGGGCC AGGCGGTCTA GCCTGGTCAC
11521 TGTCACAACG TCGCCAGGGC GTAGGTGGTC AAGCATCCTG GCCAGCTCCG GGCGGTCGCG
11581 CCTGGTGCCG GTGATCTTCT CGGAAAACAG CTTGGTGCAG CCGGCCGCGT GCAGTTCGGC
11641 CCGTTGGTTG GTCAAGTCCT GGTCGTCGGT GCTGACGCGG GCATAGCCCA GCAGGCCAGC
11701 GGCGGCGCTC TTGTTCATGG CGTAATGTCT CCGGTTCTAG TCGCAAGTAT TCTACTTTAT
11761 GCGACTAAAA CACGCGACAA GAAAACGCCA GGAAAAGGGC AGGGCGGCAG CCTGTCGCGT
11821 AACTTAGGAC TTGTGCGACA TGTCGTTTTC AGAAGACGGC TGCACTGAAC GTCAGAAGCC
11881 GACTGCACTA TAGCAGCGGA GGGGTTGGAT CAAAGTACTT TAAAGTACTT TAAAGTACTT
11941 TAAAGTACTT TGATCCCGAG GGGAACCCTG TGGTTGGCAT GCACATACAA ATGGACGAAC
12001 GGATAAACCT TTTCACGCCC TTTTAAATAT CCGATTATTC TAATAAACGC TCTTTTCTCT
12061 TAGGTTTACC CGCCAATATA TCCTGTCAAA CACTGATAGT TT
//
LOCUS D932 15649 bp DNA circular UNA 10-DEC-2018
COMMENT Insert from E443_pDONR207_GFP_myc_gVamp721 414 to 2772
FEATURES Location/Qualifiers
misc_feature 1..26
/vntifkey="21"
/locus_tag="RB"

promoter 44..1640
/vntifkey="29"
/locus_tag="5' Promoter-UBQ10"
5'UTR 1530..2029
/vntifkey="52"
/locus_tag="5'UTR"
misc_feature 1641..1725
/vntifkey="21"
/locus_tag="Exon 1.1"
intron 1726..2029
/vntifkey="15"
/locus_tag="Intron 1.1"
CDS 2037..3921
/vntifkey="4"
/locus_tag="XVE"
promoter 3937..4269
/vntifkey="29"
/locus_tag="NOS promoter"
CDS 3938..5573
/vntifkey="4"
/locus_tag="hygromycin resistance"
promoter 5576..5853
/vntifkey="29"
/locus_tag="lexA -46 35S promoter"
repeat_region 5961..5973
/vntifkey="34"
/locus_tag="attR1"
CDS 5997..5999
/locus_tag="MYC"
misc_feature 6000..6716
/locus_tag="XhoI-meGFP-SalI"
misc_feature 6262..6279
/locus_tag="gfp_seq_geno"
CDS 6720..6749
/locus_tag="MYC(1)"
misc_feature 6753..8322
/locus_tag="gVamp721"
misc_feature 7980..7999
/locus_tag="g721 seq primer"
misc_feature 8251..8313
/locus_tag="TMD"
misc_feature 8333..8418
/vntifkey="21"
/locus_tag="confirmed sequence"
repeat_region 8333..8347
/vntifkey="34"
/locus_tag="attR2"
terminator 8375..8854
/vntifkey="43"
/locus_tag="T3A"
misc_feature 9115..9139
/vntifkey="21"
/locus_tag="LB"

ORIGIN
1 GTTTACCCGC CAATATATCC TGTCAAACAC TGATAGTTTA AACAGTCTAG CTCAACAGAG
61 CTTTTAACCC AAATTGGTAC AATAGAATAC AACTTTAGAT CATAATTCTC AAAAGAAAGA
121 GATTCCTTAG CTATTCTATC TGCCACTCCA TTTCCTTCTC GGCTTGTATG CACAAGCATA
181 AAATCCTCAA ACTTGCTAAG TAGATACTTT ATGTCTTGGA TAATTGGATT GAGACTTGAC
241 AAGCATAACT TTCATGTAAC CAAAGACACA AGTTGCTGAG AATCCACCTC AAAAATGATC
301 TTCCTATAAT TGAATCGGGA TAATGACAGC ACAGCCCATC TAAGAGCCTC CACTTCTACT
361 TCCAGCACGC TTCTTACTTT TACCACAGCT CTTGCACCTA ACCATAACAC CTTCCCTGTA
421 TGATCGCGAA GCACCCACCC TAAGCCACAT TTTAATCCTT CTGTTGGCCA TGCCCCATCA
481 AAGTTGCACT TAACCCAAGA TTGTGGTGGA GCTTCCCATG TTTCTCGTCT GTCCCGACGG
541 TGTTGTGGTT GGTGCTTTCC TTACATTCTG AGCCTCTTTC CTTCTAATCC ACTCATCTGC
601 ATCTTCTTGT GTCCTTACTA ATACCTCATT GGTTCCAAAT TCCCTCCCTT TAAGCACCAG
661 CTCGTTTCTG TTCTTCCACA GCCTCCCAAG TATCCAAGGG ACTAAAGCCT CCACATTCTT
721 CAGATCAGGA TATTCTTGTT TAAGATGTTG AACTCTATGG AGGTTTGTAT GAACTGATGA
781 TCTAGGACCG GATAAGTTCC CTTCTTCATA GCGAACTTAT TCAAAGAATG TTTTGTGTAT
841 CATTCTTGTT ACATTGTTAT TAATGAAAAA ATATTATTGG TCATTGGACT GAACACGAGT
901 GTTAAATATG GACCAGGCCC CAAATAAGAT CCATTGATAT ATGAATTAAA TAACAAGAAT
961 AAATCGAGTC ACCAAACCAC TTGCCTTTTT TAACGAGACT TGTTCACCAA CTTGATACAA
1021 AAGTCATTAT CCTATGCAAA TCAATAATCA TACAAAAATA TCCAATAACA CTAAAAAATT
1081 AAAAGAAATG GATAATTTCA CAATATGTTA TACGATAAAG AAGTTACTTT TCCAAGAAAT
1141 TCACTGATTT TATAAGCCCA CTTGCATTAG ATAAATGGCA AAAAAAAACA AAAAGGAAAA
1201 GAAATAAAGC ACGAAGAATT CTAGAAAATA CGAAATACGC TTCAATGCAG TGGGACCCAC
1261 GGTTCAATTA TTGCCAATTT TCAGCTCCAC CGTATATTTA AAAAATAAAA CGATAATGCT
1321 AAAAAAATAT AAATCGTAAC GATCGTTAAA TCTCAACGGC TGGATCTTAT GACGACCGTT
1381 AGAAATTGTG GTTGTCGACG AGTCAGTAAT AAACGGCGTC AAAGTGGTTG CAGCCGGCAC
1441 ACACGAGTCG TGTTTATCAA CTCAAAGCAC AAATACTTTT CCTCAACCTA AAAATAAGGC
1501 AATTAGCCAA AAACAACTTT GCGTGTAAAC AACGCTCAAT ACACGTGTCA TTTTATTATT
1561 AGCTATTGCT TCACCGCCTT AGCTTTCTCG TGACCTAGTC GTCCTCGTCT TTTCTTCTTC
1621 TTCTTCTATA AAACAATACC CAAAGAGCTC TTCTTCTTCA CAATTCAGAT TTCAATTTCT
1681 CAAAATCTTA AAAACTTTCT CTCAATTCTC TCTACCGTGA TCAAGGTAAA TTTCTGTGTT
1741 CCTTATTCTC TCAAAATCTT CGATTTTGTT TTCGTTCGAT CCCAATTTCG TATATGTTCT
1801 TTGGTTTAGA TTCTGTTAAT CTTAGATCGA AGACGATTTT CTGGGTTTGA TCGTTAGATA
1861 TCATCTTAAT TCTCGATTAG GGTTTCATAG ATATCATCCG ATTTGTTCAA ATAATTTGAG
1921 TTTTGTCGAA TAATTACTCT TCGATTTGTG ATTTCTATCT AGATCTGGTG TTAGTTTCTA
1981 GTTTGTGCGA TCGAATTTGT CGATTAATCT GAGTTTTTCT GATTAACAGT TCGAAATGAA
2041 AGCGTTAACG GCCAGGCAAC AAGAGGTGTT TGATCTCATC CGTGATCACA TCAGCCAGAC
2101 AGGTATGCCG CCGACGCGTG CGGAAATCGC GCAGCGTTTG GGGTTCCGTT CCCCAAACGC
2161 GGCTGAAGAA CATCTGAAGG CGCTGGCACG CAAAGGCGTT ATTGAAATTG TTTCCGGCGC
2221 ATCACGCGGG ATTCGTCTGT TGCAGGAAGA GGAAGAAGGG TTGCCGCTGG TAGGTCGTGT
2281 GGCTGCCGGT GAACCGTCGA GCGCCCCCCC GACCGATGTC AGCCTGGGGG ACGAGCTCCA
2341 CTTAGACGGC GAGGACGTGG CGATGGCGCA TGCCGACGCG CTAGACGATT TCGATCTGGA
2401 CATGTTGGGG GACGGGGATT CCCCGGGTCC GGGATTTACC CCCCACGACT CCGCCCCCTA
2461 CGGCGCTCTG GATATGGCCG ACTTCGAGTT TGAGCAGATG TTTACCGATG CCCTTGGAAT
2521 TGACGAGTAC GGTGGGGATC CGTCTGCTGG AGACATGAGA GCTGCCAACC TTTGGCCAAG
2581 CCCGCTCATG ATCAAACGCT CTAAGAAGAA CAGCCTGGCC TTGTCCCTGA CGGCCGACCA
2641 GATGGTCAGT GCCTTGTTGG ATGCTGAGCC CCCCATACTC TATTCCGAGT ATGATCCTAC
2701 CAGACCCTTC AGTGAAGCTT CGATGATGGG CTTACTGACC AACCTGGCAG ACAGGGAGCT
2761 GGTTCACATG ATCAACTGGG CGAAGAGGGT GCCAGGCTTT GTGGATTTGA CCCTCCATGA
2821 TCAGGTCCAC CTTCTAGAAT GTGCCTGGCT AGAGATCCTG ATGATTGGTC TCGTCTGGCG
2881 CTCCATGGAG CACCCAGTGA AGCTACTGTT TGCTCCTAAC TTGCTCTTGG ACAGGAACCA
2941 GGGAAAATGT GTAGAGGGCA TGGTGGAGAT CTTCGACATG CTGCTGGCTA CATCATCTCG
3001 GTTCCGCATG ATGAATCTGC AGGGAGAGGA GTTTGTGTGC CTCAAATCTA TTATTTTGCT
3061 TAATTCTGGA GTGTACACAT TTCTGTCCAG CACCCTGAAG TCTCTGGAAG AGAAGGACCA

3121 TATCCACCGA GTCCTGGACA AGATCACAGA CACTTTGATC CACCTGATGG CCAAGGCAGG
3181 CCTGACCCTG CAGCAGCAGC ACCAGCGGCT GGCCCAGCTC CTCCTCATCC TCTCCCACAT
3241 CAGGCACATG AGTAACAAAG GCATGGAGCA TCTGTACAGC ATGAAGTGCA AGAACGTGGT
3301 GCCCCTCTAT GACCTGCTGC TGGAGATGCT GGACGCCCAC CGCCTACATG CGCCCACTAG
3361 CCGTGGAGGG GCATCCGTGG AGGAGACGGA CCAAAGCCAC TTGGCCACTG CGGGCTCTAC
3421 TTCATCGCAT TCCTTGCAAA AGTATTACAT CACGGGGGAG GCAGAGGGTT TCCCTGCCAC
3481 AGTCTGAGAG CTCCCTGGCG AATTCCCAGA GATGTTAGCT GAAATCATCA CTAATCAGAT
3541 ACCAAAATAT TCAAATGGAA ATATCAAAAA GCTTCTGTTT CATCAAAAAT GACTCGACCT
3601 AACTGAGTAA GCTAGCTTGT TCGAGTATTA TGGCATTGGG AAAACTGTTT TTCTTGTACC
3661 ATTTGTTGTG CTTGTAATTT ACTGTGTTTT TTATTCGGTT TTCGCTATCG AACTGTGAAA
3721 TGGAAATGGA TGGAGAAGAG TTAATGAATG ATATGGTCCT TTTGTTCATT CTCAAATTAA
3781 TATTATTTGT TTTTTCTCTT ATTTGTTGTG TGTTGAATTT GAAATTATAA GAGATATGCA
3841 AACATTTTGT TTTGAGTAAA AATGTGTCAA ATCGTGGCCT CTAATGACCG AAGTTAATAT
3901 GAGGAGTAAA ACATCCCAAA CAAGCTTGGA AACTGAAGGC GGGAAACGAC AATCTGATCA
3961 TGAGCGGAGA ATTAAGGGAG TCACGTTATG ACCCCCGCCG ATGACGCGGG ACAAGCCGTT
4021 TTACGTTTGG AACTGACAGA ACCGCAACGA TTGAAGGAGC CACTCAGCCG CGGGTTTCTG
4081 GAGTTTAATG AGCTAAGCAC ATACGTCAGA AACCATTATT GCGCGTTCAA AAGTCGCCTA
4141 AGGTCACTAT CAGCTAGCAA ATATTTCTTG TCAAAAATGC TCCACTGACG TTCCATAAAT
4201 TCCCCTCGGT ATCCAATTAG AGTCTCATAT TCACTCTCAA TCCAAATAAT CTGCACCGGA
4261 TCCCCTAGAA TGAAAAAGCC TGAACTCACC GCGACGTCTG TCGAGAAGTT TCTGATCGAA
4321 AAGTTCGACA GCGTCTCCGA CCTGATGCAG CTCTCGGAGG GCGAAGAATC TCGTGCTTTC
4381 AGCTTCGATG TAGGAGGGCG TGGATATGTC CTGCGGGTAA ATAGCTGCGC CGATGGTTTC
4441 TACAAAGATC GTTATGTTTA TCGGCACTTT GCATCGGCCG CGCTCCCGAT TCCGGAAGTG
4501 CTTGACATTG GGGAATTCAG CGAGAGCCTG ACCTATTGCA TCTCCCGCCG TGCACAGGGT
4561 GTCACGTTGC AAGACCTGCC TGAAACCGAA CTGCCCGCTG TTCTGCAGCC GGTCGCGGAG
4621 GCCATGGATG CGATCGCTGC GGCCGATCTT AGCCAGACGA GCGGGTTCGG CCCATTCGGA
4681 CCGCAAGGAA TCGGTCAATA CACTACATGG CGTGATTTCA TATGCGCGAT TGCTGATCCC
4741 CATGTGTATC ACTGGCAAAC TGTGATGGAC GACACCGTCA GTGCGTCCGT CGCGCAGGCT
4801 CTCGATGAGC TGATGCTTTG GGCCGAGGAC TGCCCCGAAG TCCGGCACCT CGTGCACGCG
4861 GATTTCGGCT CCAACAATGT CCTGACGGAC AATGGCCGCA TAACAGCGGT CATTGACTGG
4921 AGCGAGGCGA TGTTCGGGGA TTCCCAATAC GAGGTCGCCA ACATCTTCTT CTGGAGGCCG
4981 TGGTTGGCTT GTATGGAGCA GCAGACGCGC TACTTCGAGC GGAGGCATCC GGAGCTTGCA
5041 GGATCGCCGC GGCTCCGGGC GTATATGCTC CGCATTGGTC TTGACCAACT CTATCAGAGC
5101 TTGGTTGACG GCAATTTCGA TGATGCAGCT TGGGCGCAGG GTCGATGCGA CGCAATCGTC
5161 CGATCCGGAG CCGGGACTGT CGGGCGTACA CAAATCGCCC GCAGAAGCGC GGCCGTCTGG
5221 ACCGATGGCT GTGTAGAAGT ACTCGCCGAT AGTGGAAACC GACGCCCCAG CACTCGTCCG
5281 AGGGCAAAGG AATAGCGATC GTTCAAACAT TTGGCAATAA AGTTTCTTAA GATTGAATCC
5341 TGTTGCCGGT CTTGCGATGA TTATCATATA ATTTCTGTTG AATTACGTTA AGCATGTAAT
5401 AATTAACATG TAATGCATGA CGTTATTTAT GAGATGGGTT TTTATGATTA GAGTCCCGCA
5461 ATTATACATT TAATACGCGA TAGAAAACAA AATATAGCGC GCAAACTAGG ATAAATTATC
5521 GCGCGCGGTG TCATCTATGT TACTAGATCG GGGAATTGAT CCCCCCTCGA CAGCTTGCAT
5581 GCCAGCTTGG GCTGCAGGTC GAGGCTAAAA AACTAATCGC ATTATCATCC CCTCGACGTA
5641 CTGTACATAT AACCACTGGT TTTATATACA GCAGTACTGT ACATATAACC ACTGGTTTTA
5701 TATACAGCAG TCGACGTACT GTACATATAA CCACTGGTTT TATATACAGC AGTACTGTAC
5761 ATATAACCAC TGGTTTTATA TACAGCAGTC GAGGTAAGAT TAGATATGGA TATGTATATG
5821 GATATGTATA TGGTGGTAAT GCCATGTAAT ATGCTCGACT CTAGGATCTT CGCAAGACCC
5881 TTCCTCTATA TAAGGAAGTT CATTTCATTT GGAGAGGACA CGCTGAAGCT AGTCGACTCT
5941 AGCCTCGAGG CGCGCCAAGC TATCAACAAG TTTGTACAAA AAAGCAGGCT ATGCTCGAGA
6001 TGGTGAGCAA GGGCGAGGAG CTGTTCACCG GGGTGGTGCC CATCCTGGTC GAGCTGGACG
6061 GCGACGTAAA CGGCCACAAG TTCAGCGTGT CCGGCGAGGG CGAGGGCGAT GCCACCTACG
6121 GCAAGCTGAC CCTGAAGTTC ATCTGCACCA CCGGCAAGCT GCCCGTGCCC TGGCCCACCC
6181 TCGTGACCAC CCTGACCTAC GGCGTGCAGT GCTTCAGCCG CTACCCCGAC CACATGAAGC
6241 AGCACGACTT CTTCAAGTCC GCCATGCCCG AAGGCTACGT CCAGGAGCGC ACCATCTTCT

6301 TCAAGGACGA CGGCAACTAC AAGACCCGCG CCGAGGTGAA GTTCGAGGGC GACACCCTGG
6361 TGAACCGCAT CGAGCTGAAG GGCATCGACT TCAAGGAGGA CGGCAACATC CTGGGGCACA
6421 AGCTGGAGTA CAACTACAAC AGCCACAACG TCTATATCAT GGCCGACAAG CAGAAGAACG
6481 GCATCAAGGT GAACTTCAAG ATCCGCCACA ACATCGAGGA CGGCAGCGTG CAGCTCGCCG
6541 ACCACTACCA GCAGAACACC CCCATCGGCG ACGGCCCCGT GCTGCTGCCC GACAACCACT
6601 ACCTGAGTAC TCAGTCCAAG CTGAGCAAAG ACCCCAACGA GAAGCGCGAT CACATGGTCC
6661 TGCTGGAGTT CGTGACCGCC GCCGGGATCA CTCTCGGCAT GGACGAGCTG TACAAGGTCG
6721 AGCAGAAGTT GATCTCAGAG GAGGACTTAG GCATGGCGCA ACAATCGTTG ATCTACAGTT
6781 TCGTAGCTCG CGGCACGGTG ATCCTCGTTG AGTTCACTGA TTTCAAAGGT AATTTCACCT
6841 CAATCGCTGC TCAGTGCCTC CAGAAGCTTC CGTCTTCGAA CAACAAGTTC ACCTACAACT
6901 GCGACGGTCA TACCTTCAAT TACCTTGTCG AAGATGGATT CAGTAAGTCA CTTTTCGTTT
6961 GATCTATGCA TAGGTTTTAA TCTTACACCA TTCGGTGCTT ACCTCGATTT GTTTCTAGGT
7021 TTATTTGCCT AACCTATCGA TTCGTATGGA TTTGTGCATC CTGAGTAGTC TGATTTCACT
7081 GAGGAACCGC ATGAACCAAG CTTTTAGGGT TAGATCTATA TTTATTAAAC GCAATAAATC
7141 TCTGTAGAGT CTTACCTGTA TAATCACTCA GATTTGGACA GAATCCGTGG TAGCTTTGGT
7201 TGATACTATG TGGAAGAACA CCAACATTGA TCTGAGACAG GGTTTTTACT GGTTGCTTTA
7261 TTTGCGTATA GTTTCCTAGG TCATGGGTTT TGGCTGGCTT CATTTTTCAA TGAAAATCTG
7321 GGGTCTTATT GTAAGCTTTT GTGACTATCC TTTAGTTGTT GTAGCCATAT CAATTTCATA
7381 TTAAATCAAT GTCACCTGAT TATCCCGTAA GGGAAGACAA GGAGATTGAT TTTGTTCCTG
7441 TTGTACAGCC TATTGTGTTG TTGCGGTTGA TTCTGCTGGG AGGCAAATTC CCATGTCCTT
7501 TTTGGAAAGA GTAAAAGAAG ATTTTAACAA GCGATATGGT GGTGGAAAGG CTGCAACTGC
7561 TCAAGCAAAC AGCTTGAATA AGGAGTTTGG GTACTTTTTC ATTATCTCTT CTATTTGGAT
7621 GATCTTCTCT TTTATATTCG TGGACTGACT CTTTTTGGTA ACTTTGAAGC TCTAAACTGA
7681 AAGAGCATAT GCAGTATTGC ATGGATCATC CTGATGAGAT TAGCAAGCTT GCTAAGGTGA
7741 AGGCGCAAGT GTCAGAAGTT AAGGGTGTAA TGATGGAAAA CATTGAGAAG GTTTGAATCT
7801 GACCCTTTCT GTCCTCAATG TATATATTTA CATCTATGGT TGACCCATCT GAAAGAGCTC
7861 ATCTAGTCAT ACTAAGTTAC TGTGAAATCA ATTACTAATA ATTCAATGCA TTACATCTTA
7921 CGGGAAATGT CCAGTTTACC AATGTAACAC CGCTTACATA TGCGACTCAT TATTTGCAGG
7981 TTCTTGACCG TGGTGAGAAA ATTGAGCTTT TGGTGGACAA AACCGAAAAT CTTCGCTCAC
8041 AGGTTAGACT CTCTCATACC CTTATCTCCT GCATCTATTT GCCTACACTC ACACACAAAG
8101 ATGAACATGC TTTTTGAGTT CATAGCTGAC TCATCCTATT ACCATATCTT GTGTCTAACT
8161 GAACTCAAAC TGCCACAGGC ACAAGATTTC AGAACAACAG GAACGCAGAT GAGAAGAAAG
8221 ATGTGGCTTC AGAACATGAA GATAAAACTC ATAGTGCTCG CCATCATTAT CGCACTGATT
8281 CTCATCATCG TGCTCTCAGT TTGCCATGGG TTTAAGTGTT AAACCCAGCT TTCTTGTACA
8341 AAGTGGTTGA TAATTCTTAA TTAACTAGTC GATCCAGGCC TCCCAGCTTT CGTCCGTATC
8401 ATCGGTTTCG ACAACGTTCG TCAAGTTCAA TGCATCAGTT TCATTGCCCA CACACCAGAA
8461 TCCTACTAAG TTTGAGTATT ATGGCATTGG AAAAGCTGTT TTCTTCTATC ATTTGTTCTG
8521 CTTGTAATTT ACTGTGTTCT TTCAGTTTTT GTTTTCGGAC ATCAAAATGC AAATGGATGG
8581 ATAAGAGTTA ATAAATGATA TGGTCCTTTT GTTCATTCTC AAATTATTAT TATCTGTTGT
8641 TTTTACTTTA ATGGGTTGAA TTTAAGTAAG AAAGGAACTA ACAGTGTGAT ATTAAGGTGC
8701 AATGTTAGAC ATATAAAACA GTCTTTCACC TCTCTTTGGT TATGTCTTGA ATTGGTTTGT
8761 TTCTTCACTT ATCTGTGTAA TCAAGTTTAC TATGAGTCTA TGATCAAGTA ATTATGCAAT
8821 CAAGTTAAGT ACAGTATAGG CTTTTTGTGT CGAGGGGGTA CCGAGTCGAG GAATTCACTG
8881 GCCGTCGTTT TACAACGTCG TGACTGGGAA AACCCTGGCG TTACCCAACT TAATCGCCTT
8941 GCAGCACATC CCCCTTTCGC CAGCTGGCGT AATAGCGAAG AGGCCCGCAC CGATCGCCCT
9001 TCCCAACAGT TGCGCAGCCT GAATGGCGGG TACCGAGCTC GAATTCAATT CGGCGTTAAT
9061 TCAGTACATT AAAAACGTCC GCAATGTGTT ATTAAGTTGT CTAAGCGTCA ATTTGTTTAC
9121 ACCACAATAT ATCCTGCCAC CAGCCAGCCA ACAGCTCCCC GACCGGCAGC TCGGCACAAA
9181 ATCACCACTC GATACAGGCA GCCCATCAGT CCGGGACGGC GTCAGCGGGA GAGCCGTTGT
9241 AAGGCGGCAG ACTTTGCTCA TGTTACCGAT GCTATTCGGA AGAACGGCAA CTAAGCTGCC
9301 GGGTTTGAAA CACGGATGAT CTCGCGGAGG GTAGCATGTT GATTGTAACG ATGACAGAGC
9361 GTTGCTGCCT GTGATCAATT CGGGCACGAA CCCAGTGGAC ATAAGCCTCG TTCGGTTCGT
9421 AAGCTGTAAT GCAAGTAGCG TAACTGCCGT CACGCAACTG GTCCAGAACC TTGACCGAAC

9481 GCAGCGGTGG TAACGGCGCA GTGGCGGTTT TCATGGCTTC TTGTTATGAC ATGTTTTTTT
9541 GGGGTACAGT CTATGCCTCG GGCATCCAAG CAGCAAGCGC GTTACGCCGT GGGTCGATGT
9601 TTGATGTTAT GGAGCAGCAA CGATGTTACG CAGCAGGGCA GTCGCCCTAA AACAAAGTTA
9661 AACATCATGG GGGAAGCGGT GATCGCCGAA GTATCGACTC AACTATCAGA GGTAGTTGGC
9721 GTCATCGAGC GCCATCTCGA ACCGACGTTG CTGGCCGTAC ATTTGTACGG CTCCGCAGTG
9781 GATGGCGGCC TGAAGCCACA CAGTGATATT GATTTGCTGG TTACGGTGAC CGTAAGGCTT
9841 GATGAAACAA CGCGGCGAGC TTTGATCAAC GACCTTTTGG AAACTTCGGC TTCCCCTGGA
9901 GAGAGCGAGA TTCTCCGCGC TGTAGAAGTC ACCATTGTTG TGCACGACGA CATCATTCCG
9961 TGGCGTTATC CAGCTAAGCG CGAACTGCAA TTTGGAGAAT GGCAGCGCAA TGACATTCTT
10021 GCAGGTATCT TCGAGCCAGC CACGATCGAC ATTGATCTGG CTATCTTGCT GACAAAAGCA
10081 AGAGAACATA GCGTTGCCTT GGTAGGTCCA GCGGCGGAGG AACTCTTTGA TCCGGTTCCT
10141 GAACAGGATC TATTTGAGGC GCTAAATGAA ACCTTAACGC TATGGAACTC GCCGCCCGAC
10201 TGGGCTGGCG ATGAGCGAAA TGTAGTGCTT ACGTTGTCCC GCATTTGGTA CAGCGCAGTA
10261 ACCGGCAAAA TCGCGCCGAA GGATGTCGCT GCCGACTGGG CAATGGAGCG CCTGCCGGCC
10321 CAGTATCAGC CCGTCATACT TGAAGCTAGA CAGGCTTATC TTGGACAAGA AGAAGATCGC
10381 TTGGCCTCGC GCGCAGATCA GTTGGAAGAA TTTGTCCACT ACGTGAAAGG CGAGATCACC
10441 AAGGTAGTCG GCAAATAATG TCTAGCTAGA AATTCGTTCA AGCCGACGCC GCTTCGCCGG
10501 CGTTAACTCA AGCGATTAGA TGCACTAAGC ACATAATTGC TCACAGCCAA ACTATCAGGT
10561 CAAGTCTGCT TTTATTATTT TTAAGCGTGC ATAATAAGCC CTACACAAAT TGGGAGATAT
10621 ATCATGCATG ACCAAAATCC CTTAACGTGA GTTTTCGTTC CACTGAGCGT CAGACCCCGT
10681 AGAAAAGATC AAAGGATCTT CTTGAGATCC TTTTTTTCTG CGCGTAATCT GCTGCTTGCA
10741 AACAAAAAAA CCACCGCTAC CAGCGGTGGT TTGTTTGCCG GATCAAGAGC TACCAACTCT
10801 TTTTCCGAAG GTAACTGGCT TCAGCAGAGC GCAGATACCA AATACTGTCC TTCTAGTGTA
10861 GCCGTAGTTA GGCCACCACT TCAAGAACTC TGTAGCACCG CCTACATACC TCGCTCTGCT
10921 AATCCTGTTA CCAGTGGCTG CTGCCAGTGG CGATAAGTCG TGTCTTACCG GGTTGGACTC
10981 AAGACGATAG TTACCGGATA AGGCGCAGCG GTCGGGCTGA ACGGGGGGTT CGTGCACACA
11041 GCCCAGCTTG GAGCGAACGA CCTACACCGA ACTGAGATAC CTACAGCGTG AGCTATGAGA
11101 AAGCGCCACG CTTCCCGAAG GGAGAAAGGC GGACAGGTAT CCGGTAAGCG GCAGGGTCGG
11161 AACAGGAGAG CGCACGAGGG AGCTTCCAGG GGGAAACGCC TGGTATCTTT ATAGTCCTGT
11221 CGGGTTTCGC CACCTCTGAC TTGAGCGTCG ATTTTTGTGA TGCTCGTCAG GGGGGCGGAG
11281 CCTATGGAAA AACGCCAGCA ACGCGGCCTT TTTACGGTTC CTGGCCTTTT GCTGGCCTTT
11341 TGCTCACATG TTCTTTCCTG CGTTATCCCC TGATTCTGTG GATAACCGTA TTACCGCCTT
11401 TGAGTGAGCT GATACCGCTC GCCGCAGCCG AACGACCGAG CGCAGCGAGT CAGTGAGCGA
11461 GGAAGCGGAA GAGCGCCTGA TGCGGTATTT TCTCCTTACG CATCTGTGCG GTATTTCACA
11521 CCGCATATGG TGCACTCTCA GTACAATCTG CTCTGATGCC GCATAGTTAA GCCAGTATAC
11581 ACTCCGCTAT CGCTACGTGA CTGGGTCATG GCTGCGCCCC GACACCCGCC AACACCCGCT
11641 GACGCGCCCT GACGGGCTTG TCTGCTCCCG GCATCCGCTT ACAGACAAGC TGTGACCGTC
11701 TCCGGGAGCT GCATGTGTCA GAGGTTTTCA CCGTCATCAC CGAAACGCGC GAGGCAGGGT
11761 GCCTTGATGT GGGCGCCGGC GGTCGAGTGG CGACGGCGCG GCTTGTCCGC GCCCTGGTAG
11821 ATTGCCTGGC CGTAGGCCAG CCATTTTTGA GCGGCCAGCG GCCGCGATAG GCCGACGCGA
11881 AGCGGCGGGG CGTAGGGAGC GCAGCGACCG AAGGGTAGGC GCTTTTTGCA GCTCTTCGGC
11941 TGTGCGCTGG CCAGACAGTT ATGCACAGGC CAGGCGGGTT TTAAGAGTTT TAATAAGTTT
12001 TAAAGAGTTT TAGGCGGAAA AATCGCCTTT TTTCTCTTTT ATATCAGTCA CTTACATGTG
12061 TGACCGGTTC CCAATGTACG GCTTTGGGTT CCCAATGTAC GGGTTCCGGT TCCCAATGTA
12121 CGGCTTTGGG TTCCCAATGT ACGTGCTATC CACAGGAAAG AGACCTTTTC GACCTTTTTC
12181 CCCTGCTAGG GCAATTTGCC CTAGCATCTG CTCCGTACAT TAGGAACCGG CGGATGCTTC
12241 GCCCTCGATC AGGTTGCGGT AGCGCATGAC TAGGATCGGG CCAGCCTGCC CCGCCTCCTC
12301 CTTCAAATCG TACTCCGGCA GGTCATTTGA CCCGATCAGC TTGCGCACGG TGAAACAGAA
12361 CTTCTTGAAC TCTCCGGCGC TGCCACTGCG TTCGTAGATC GTCTTGAACA ACCATCTGGC
12421 TTCTGCCTTG CCTGCGGCGC GGCGTGCCAG GCGGTAGAGA AAACGGCCGA TGCCGGGATC
12481 GATCAAAAAG TAATCGGGGT GAACCGTCAG CACGTCCGGG TTCTTGCCTT CTGTGATCTC
12541 GCGGTACATC CAATCAGCTA GCTCGATCTC GATGTACTCC GGCCGCCCGG TTTCGCTCTT
12601 TACGATCTTG TAGCGGCTAA TCAAGGCTTC ACCCTCGGAT ACCGTCACCA GGCGGCCGTT

12661 CTTGGCCTTC TTCGTACGCT GCATGGCAAC GTGCGTGGTG TTTAACCGAA TGCAGGTTTC
12721 TACCAGGTCG TCTTTCTGCT TTCCGCCATC GGCTCGCCGG CAGAACTTGA GTACGTCCGC
12781 AACGTGTGGA CGGAACACGC GGCCGGGCTT GTCTCCCTTC CCTTCCCGGT ATCGGTTCAT
12841 GGATTCGGTT AGATGGGAAA CCGCCATCAG TACCAGGTCG TAATCCCACA CACTGGCCAT
12901 GCCGGCCGGC CCTGCGGAAA CCTCTACGTG CCCGTCTGGA AGCTCGTAGC GGATCACCTC
12961 GCCAGCTCGT CGGTCACGCT TCGACAGACG GAAAACGGCC ACGTCCATGA TGCTGCGACT
13021 ATCGCGGGTG CCCACGTCAT AGAGCATCGG AACGAAAAAA TCTGGTTGCT CGTCGCCCTT
13081 GGGCGGCTTC CTAATCGACG GCGCACCGGC TGCCGGCGGT TGCCGGGATT CTTTGCGGAT
13141 TCGATCAGCG GCCGCTTGCC ACGATTCACC GGGGCGTGCT TCTGCCTCGA TGCGTTGCCG
13201 CTGGGCGGCC TGCGCGGCCT TCAACTTCTC CACCAGGTCA TCACCCAGCG CCGCGCCGAT
13261 TTGTACCGGG CCGGATGGTT TGCGACCGTC ACGCCGATTC CTCGGGCTTG GGGGTTCCAG
13321 TGCCATTGCA GGGCCGGCAG ACAACCCAGC CGCTTACGCC TGGCCAACCG CCCGTTCCTC
13381 CACACATGGG GCATTCCACG GCGTCGGTGC CTGGTTGTTC TTGATTTTCC ATGCCGCCTC
13441 CTTTAGCCGC TAAAATTCAT CTACTCATTT ATTCATTTGC TCATTTACTC TGGTAGCTGC
13501 GCGATGTATT CAGATAGCAG CTCGGTAATG GTCTTGCCTT GGCGTACCGC GTACATCTTC
13561 AGCTTGGTGT GATCCTCCGC CGGCAACTGA AAGTTGACCC GCTTCATGGC TGGCGTGTCT
13621 GCCAGGCTGG CCAACGTTGC AGCCTTGCTG CTGCGTGCGC TCGGACGGCC GGCACTTAGC
13681 GTGTTTGTGC TTTTGCTCAT TTTCTCTTTA CCTCATTAAC TCAAATGAGT TTTGATTTAA
13741 TTTCAGCGGC CAGCGCCTGG ACCTCGCGGG CAGCGTCGCC CTCGGGTTCT GATTCAAGAA
13801 CGGTTGTGCC GGCGGCGGCA GTGCCTGGGT AGCTCACGCG CTGCGTGATA CGGGACTCAA
13861 GAATGGGCAG CTCGTACCCG GCCAGCGCCT CGGCAACCTC ACCGCCGATG CGCGTGCCTT
13921 TGATCGCCCG CGACACGACA AAGGCCGCTT GTAGCCTTCC ATCCGTGACC TCAATGCGCT
13981 GCTTAACCAG CTCCACCAGG TCGGCGGTGG CCCATATGTC GTAAGGGCTT GGCTGCACCG
14041 GAATCAGCAC GAAGTCGGCT GCCTTGATCG CGGACACAGC CAAGTCCGCC GCCTGGGGCG
14101 CTCCGTCGAT CACTACGAAG TCGCGCCGGC CGATGGCCTT CACGTCGCGG TCAATCGTCG
14161 GGCGGTCGAT GCCGACAACG GTTAGCGGTT GATCTTCCCG CACGGCCGCC CAATCGCGGG
14221 CACTGCCCTG GGGATCGGAA TCGACTAACA GAACATCGGC CCCGGCGAGT TGCAGGGCGC
14281 GGGCTAGATG GGTTGCGATG GTCGTCTTGC CTGACCCGCC TTTCTGGTTA AGTACAGCGA
14341 TAACCTTCAT GCGTTCCCCT TGCGTATTTG TTTATTTACT CATCGCATCA TATACGCAGC
14401 GACCGCATGA CGCAAGCTGT TTTACTCAAA TACACATCAC CTTTTTAGAC GGCGGCGCTC
14461 GGTTTCTTCA GCGGCCAAGC TGGCCGGCCA GGCCGCCAGC TTGGCATCAG ACAAACCGGC
14521 CAGGATTTCA TGCAGCCGCA CGGTTGAGAC GTGCGCGGGC GGCTCGAACA CGTACCCGGC
14581 CGCGATCATC TCCGCCTCGA TCTCTTCGGT AATGAAAAAC GGTTCGTCCT GGCCGTCCTG
14641 GTGCGGTTTC ATGCTTGTTC CTCTTGGCGT TCATTCTCGG CGGCCGCCAG GGCGTCGGCC
14701 TCGGTCAATG CGTCCTCACG GAAGGCACCG CGCCGCCTGG CCTCGGTGGG CGTCACTTCC
14761 TCGCTGCGCT CAAGTGCGCG GTACAGGGTC GAGCGATGCA CGCCAAGCAG TGCAGCCGCC
14821 TCTTTCACGG TGCGGCCTTC CTGGTCGATC AGCTCGCGGG CGTGCGCGAT CTGTGCCGGG
14881 GTGAGGGTAG GGCGGGGGCC AAACTTCACG CCTCGGGCCT TGGCGGCCTC GCGCCCGCTC
14941 CGGGTGCGGT CGATGATTAG GGAACGCTCG AACTCGGCAA TGCCGGCGAA CACGGTCAAC
15001 ACCATGCGGC CGGCCGGCGT GGTGGTGTCG GCCCACGGCT CTGCCAGGCT ACGCAGGCCC
15061 GCGCCGGCCT CCTGGATGCG CTCGGCAATG TCCAGTAGGT CGCGGGTGCT GCGGGCCAGG
15121 CGGTCTAGCC TGGTCACTGT CACAACGTCG CCAGGGCGTA GGTGGTCAAG CATCCTGGCC
15181 AGCTCCGGGC GGTCGCGCCT GGTGCCGGTG ATCTTCTCGG AAAACAGCTT GGTGCAGCCG
15241 GCCGCGTGCA GTTCGGCCCG TTGGTTGGTC AAGTCCTGGT CGTCGGTGCT GACGCGGGCA
15301 TAGCCCAGCA GGCCAGCGGC GGCGCTCTTG TTCATGGCGT AATGTCTCCG GTTCTAGTCG
15361 CAAGTATTCT ACTTTATGCG ACTAAAACAC GCGACAAGAA AACGCCAGGA AAAGGGCAGG
15421 GCGGCAGCCT GTCGCGTAAC TTAGGACTTG TGCGACATGT CGTTTTCAGA AGACGGCTGC
15481 ACTGAACGTC AGAAGCCGAC TGCACTATAG CAGCGGAGGG GTTGGATCAA AGTACTTTGA
15541 TCCCGAGGGG AACCCTGTGG TTGGCATGCA CATACAAATG GACGAACGGA TAAACCTTTT
15601 CACGCCCTTT TAAATATCCG TTATTCTAAT AAACGCTCTT TTCTCTTAG
//
LOCUS D933 15649 bp DNA circular UNA 10-DEC-2018
COMMENT Insert from E444_pDONR207_GFP_myc_gVamp721_Y57D 414 to 2772

```
FEATURES             Location/Qualifiers
     misc_feature    1..26
                     /vntifkey="21"
                     /locus_tag="RB"
     promoter        44..1640
                     /vntifkey="29"
                     /locus_tag="5' Promoter-UBQ10"
     5'UTR           1530..2029
                     /vntifkey="52"
                     /locus_tag="5'UTR"
     misc_feature    1641..1725
                     /vntifkey="21"
                     /locus_tag="Exon 1.1"
     intron          1726..2029
                     /vntifkey="15"
                     /locus_tag="Intron 1.1"
     CDS             2037..3921
                     /vntifkey="4"
                     /locus_tag="XVE"
     promoter        3937..4269
                     /vntifkey="29"
                     /locus_tag="NOS promoter"
     CDS             3938..5573
                     /vntifkey="4"
                     /locus_tag="hygromycin resistance"
     promoter        5576..5853
                     /vntifkey="29"
                     /locus_tag="lexA -46 35S promoter"
     repeat_region   5961..5973
                     /vntifkey="34"
                     /locus_tag="attR1"
     CDS             5997..5999
                     /locus_tag="MYC(1)"
     misc_feature    6000..6716
                     /locus_tag="XhoI-meGFP-SalI"
     CDS             6720..6749
                     /locus_tag="MYC"
     misc_feature    join(6753..6896,6937..8322)
                     /locus_tag="gVamp721"
     misc_feature    6897..6936
                     /locus_tag="mutated sequence Y57D + MscI site"
     misc_feature    7980..7999
                     /locus_tag="g721 seq primer"
     misc_feature    8333..8418
                     /vntifkey="21"
                     /locus_tag="confirmed sequence"
     repeat_region   8333..8347
                     /vntifkey="34"
                     /locus_tag="attR2"
     terminator      8375..8854
                     /vntifkey="43"
                     /locus_tag="T3A"
     misc_feature    9115..9139
```

/vntifkey="21"
/locus_tag="LB"
ORIGIN
1 GTTTACCCGC CAATATATCC TGTCAAACAC TGATAGTTTA AACAGTCTAG CTCAACAGAG
61 CTTTTAACCC AAATTGGTAC AATAGAATAC AACTTTAGAT CATAATTCTC AAAAGAAAGA
121 GATTCCTTAG CTATTCTATC TGCCACTCCA TTTCCTTCTC GGCTTGTATG CACAAGCATA
181 AAATCCTCAA ACTTGCTAAG TAGATACTTT ATGTCTTGGA TAATTGGATT GAGACTTGAC
241 AAGCATAACT TTCATGTAAC CAAAGACACA AGTTGCTGAG AATCCACCTC AAAAATGATC
301 TTCCTATAAT TGAATCGGGA TAATGACAGC ACAGCCCATC TAAGAGCCTC CACTTCTACT
361 TCCAGCACGC TTCTTACTTT TACCACAGCT CTTGCACCTA ACCATAACAC CTTCCCTGTA
421 TGATCGCGAA GCACCCACCC TAAGCCACAT TTTAATCCTT CTGTTGGCCA TGCCCCATCA
481 AAGTTGCACT TAACCCAAGA TTGTGGTGGA GCTTCCCATG TTTCTCGTCT GTCCCGACGG
541 TGTTGTGGTT GGTGCTTTCC TTACATTCTG AGCCTCTTTC CTTCTAATCC ACTCATCTGC
601 ATCTTCTTGT GTCCTTACTA ATACCTCATT GGTTCCAAAT TCCCTCCCTT TAAGCACCAG
661 CTCGTTTCTG TTCTTCCACA GCCTCCCAAG TATCCAAGGG ACTAAAGCCT CCACATTCTT
721 CAGATCAGGA TATTCTTGTT TAAGATGTTG AACTCTATGG AGGTTTGTAT GAACTGATGA
781 TCTAGGACCG GATAAGTTCC CTTCTTCATA GCGAACTTAT TCAAAGAATG TTTTGTGTAT
841 CATTCTTGTT ACATTGTTAT TAATGAAAAA ATATTATTGG TCATTGGACT GAACACGAGT
901 GTTAAATATG GACCAGGCCC CAAATAAGAT CCATTGATAT ATGAATTAAA TAACAAGAAT
961 AAATCGAGTC ACCAAACCAC TTGCCTTTTT TAACGAGACT TGTTCACCAA CTTGATACAA
1021 AAGTCATTAT CCTATGCAAA TCAATAATCA TACAAAAATA TCCAATAACA CTAAAAAATT
1081 AAAAGAAATG GATAATTTCA CAATATGTTA TACGATAAAG AAGTTACTTT TCCAAGAAAT
1141 TCACTGATTT TATAAGCCCA CTTGCATTAG ATAAATGGCA AAAAAAAACA AAAAGGAAAA
1201 GAAATAAAGC ACGAAGAATT CTAGAAAATA CGAAATACGC TTCAATGCAG TGGGACCCAC
1261 GGTTCAATTA TTGCCAATTT TCAGCTCCAC CGTATATTTA AAAAATAAAA CGATAATGCT
1321 AAAAAAATAT AAATCGTAAC GATCGTTAAA TCTCAACGGC TGGATCTTAT GACGACCGTT
1381 AGAAATTGTG GTTGTCGACG AGTCAGTAAT AAACGGCGTC AAAGTGGTTG CAGCCGGCAC
1441 ACACGAGTCG TGTTTATCAA CTCAAAGCAC AAATACTTTT CCTCAACCTA AAAATAAGGC
1501 AATTAGCCAA AAACAACTTT GCGTGTAAAC AACGCTCAAT ACACGTGTCA TTTTATTATT
1561 AGCTATTGCT TCACCGCCTT AGCTTTCTCG TGACCTAGTC GTCCTCGTCT TTTCTTCTTC
1621 TTCTTCTATA AAACAATACC CAAAGAGCTC TTCTTCTTCA CAATTCAGAT TTCAATTTCT
1681 CAAAATCTTA AAAACTTTCT CTCAATTCTC TCTACCGTGA TCAAGGTAAA TTTCTGTGTT
1741 CCTTATTCTC TCAAAATCTT CGATTTTGTT TTCGTTCGAT CCCAATTTCG TATATGTTCT
1801 TTGGTTTAGA TTCTGTTAAT CTTAGATCGA AGACGATTTT CTGGGTTTGA TCGTTAGATA
1861 TCATCTTAAT TCTCGATTAG GGTTTCATAG ATATCATCCG ATTTGTTCAA ATAATTTGAG
1921 TTTTGTCGAA TAATTACTCT TCGATTTGTG ATTTCTATCT AGATCTGGTG TTAGTTTCTA
1981 GTTTGTGCGA TCGAATTTGT CGATTAATCT GAGTTTTTCT GATTAACAGT TCGAAATGAA
2041 AGCGTTAACG GCCAGGCAAC AAGAGGTGTT TGATCTCATC CGTGATCACA TCAGCCAGAC
2101 AGGTATGCCG CCGACGCGTG CGGAAATCGC GCAGCGTTTG GGGTTCCGTT CCCCAAACGC
2161 GGCTGAAGAA CATCTGAAGG CGCTGGCACG CAAAGGCGTT ATTGAAATTG TTTCCGGCGC
2221 ATCACGCGGG ATTCGTCTGT TGCAGGAAGA GGAAGAAGGG TTGCCGCTGG TAGGTCGTGT
2281 GGCTGCCGGT GAACCGTCGA GCGCCCCCCC GACCGATGTC AGCCTGGGGG ACGAGCTCCA
2341 CTTAGACGGC GAGGACGTGG CGATGGCGCA TGCCGACGCG CTAGACGATT TCGATCTGGA
2401 CATGTTGGGG GACGGGGATT CCCCGGGTCC GGGATTTACC CCCCACGACT CCGCCCCCTA
2461 CGGCGCTCTG GATATGGCCG ACTTCGAGTT TGAGCAGATG TTTACCGATG CCCTTGGAAT
2521 TGACGAGTAC GGTGGGGATC CGTCTGCTGG AGACATGAGA GCTGCCAACC TTTGGCCAAG
2581 CCCGCTCATG ATCAAACGCT CTAAGAAGAA CAGCCTGGCC TTGTCCCTGA CGGCCGACCA
2641 GATGGTCAGT GCCTTGTTGG ATGCTGAGCC CCCCATACTC TATTCCGAGT ATGATCCTAC
2701 CAGACCCTTC AGTGAAGCTT CGATGATGGG CTTACTGACC AACCTGGCAG ACAGGGAGCT
2761 GGTTCACATG ATCAACTGGG CGAAGAGGGT GCCAGGCTTT GTGGATTTGA CCCTCCATGA
2821 TCAGGTCCAC CTTCTAGAAT GTGCCTGGCT AGAGATCCTG ATGATTGGTC TCGTCTGGCG
2881 CTCCATGGAG CACCCAGTGA AGCTACTGTT TGCTCCTAAC TTGCTCTTGG ACAGGAACCA
2941 GGGAAAATGT GTAGAGGGCA TGGTGGAGAT CTTCGACATG CTGCTGGCTA CATCATCTCG

3001 GTTCCGCATG ATGAATCTGC AGGGAGAGGA GTTTGTGTGC CTCAAATCTA TTATTTTGCT
3061 TAATTCTGGA GTGTACACAT TTCTGTCCAG CACCCTGAAG TCTCTGGAAG AGAAGGACCA
3121 TATCCACCGA GTCCTGGACA AGATCACAGA CACTTTGATC CACCTGATGG CCAAGGCAGG
3181 CCTGACCCTG CAGCAGCAGC ACCAGCGGCT GGCCCAGCTC CTCCTCATCC TCTCCCACAT
3241 CAGGCACATG AGTAACAAAG GCATGGAGCA TCTGTACAGC ATGAAGTGCA AGAACGTGGT
3301 GCCCCTCTAT GACCTGCTGC TGGAGATGCT GGACGCCCAC CGCCTACATG CGCCCACTAG
3361 CCGTGGAGGG GCATCCGTGG AGGAGACGGA CCAAAGCCAC TTGGCCACTG CGGGCTCTAC
3421 TTCATCGCAT TCCTTGCAAA AGTATTACAT CACGGGGGAG GCAGAGGGTT TCCCTGCCAC
3481 AGTCTGAGAG CTCCCTGGCG AATTCCCAGA GATGTTAGCT GAAATCATCA CTAATCAGAT
3541 ACCAAAATAT TCAAATGGAA ATATCAAAAA GCTTCTGTTT CATCAAAAAT GACTCGACCT
3601 AACTGAGTAA GCTAGCTTGT TCGAGTATTA TGGCATTGGG AAAACTGTTT TTCTTGTACC
3661 ATTTGTTGTG CTTGTAATTT ACTGTGTTTT TTATTCGGTT TTCGCTATCG AACTGTGAAA
3721 TGGAAATGGA TGGAGAAGAG TTAATGAATG ATATGGTCCT TTTGTTCATT CTCAAATTAA
3781 TATTATTTGT TTTTTCTCTT ATTTGTTGTG TGTTGAATTT GAAATTATAA GAGATATGCA
3841 AACATTTTGT TTTGAGTAAA AATGTGTCAA ATCGTGGCCT CTAATGACCG AAGTTAATAT
3901 GAGGAGTAAA ACATCCCAAA CAAGCTTGGA AACTGAAGGC GGGAAACGAC AATCTGATCA
3961 TGAGCGGAGA ATTAAGGGAG TCACGTTATG ACCCCCGCCG ATGACGCGGG ACAAGCCGTT
4021 TTACGTTTGG AACTGACAGA ACCGCAACGA TTGAAGGAGC CACTCAGCCG CGGGTTTCTG
4081 GAGTTTAATG AGCTAAGCAC ATACGTCAGA AACCATTATT GCGCGTTCAA AAGTCGCCTA
4141 AGGTCACTAT CAGCTAGCAA ATATTTCTTG TCAAAAATGC TCCACTGACG TTCCATAAAT
4201 TCCCCTCGGT ATCCAATTAG AGTCTCATAT TCACTCTCAA TCCAAATAAT CTGCACCGGA
4261 TCCCCTAGAA TGAAAAAGCC TGAACTCACC GCGACGTCTG TCGAGAAGTT TCTGATCGAA
4321 AAGTTCGACA GCGTCTCCGA CCTGATGCAG CTCTCGGAGG GCGAAGAATC TCGTGCTTTC
4381 AGCTTCGATG TAGGAGGGCG TGGATATGTC CTGCGGGTAA ATAGCTGCGC CGATGGTTTC
4441 TACAAAGATC GTTATGTTTA TCGGCACTTT GCATCGGCCG CGCTCCCGAT TCCGGAAGTG
4501 CTTGACATTG GGGAATTCAG CGAGAGCCTG ACCTATTGCA TCTCCCGCCG TGCACAGGGT
4561 GTCACGTTGC AAGACCTGCC TGAAACCGAA CTGCCCGCTG TTCTGCAGCC GGTCGCGGAG
4621 GCCATGGATG CGATCGCTGC GGCCGATCTT AGCCAGACGA GCGGGTTCGG CCCATTCGGA
4681 CCGCAAGGAA TCGGTCAATA CACTACATGG CGTGATTTCA TATGCGCGAT TGCTGATCCC
4741 CATGTGTATC ACTGGCAAAC TGTGATGGAC GACACCGTCA GTGCGTCCGT CGCGCAGGCT
4801 CTCGATGAGC TGATGCTTTG GGCCGAGGAC TGCCCCGAAG TCCGGCACCT CGTGCACGCG
4861 GATTTCGGCT CCAACAATGT CCTGACGGAC AATGGCCGCA TAACAGCGGT CATTGACTGG
4921 AGCGAGGCGA TGTTCGGGGA TTCCCAATAC GAGGTCGCCA ACATCTTCTT CTGGAGGCCG
4981 TGGTTGGCTT GTATGGAGCA GCAGACGCGC TACTTCGAGC GGAGGCATCC GGAGCTTGCA
5041 GGATCGCCGC GGCTCCGGGC GTATATGCTC CGCATTGGTC TTGACCAACT CTATCAGAGC
5101 TTGGTTGACG GCAATTTCGA TGATGCAGCT TGGGCGCAGG GTCGATGCGA CGCAATCGTC
5161 CGATCCGGAG CCGGGACTGT CGGGCGTACA CAAATCGCCC GCAGAAGCGC GGCCGTCTGG
5221 ACCGATGGCT GTGTAGAAGT ACTCGCCGAT AGTGGAAACC GACGCCCCAG CACTCGTCCG
5281 AGGGCAAAGG AATAGCGATC GTTCAAACAT TTGGCAATAA AGTTTCTTAA GATTGAATCC
5341 TGTTGCCGGT CTTGCGATGA TTATCATATA ATTTCTGTTG AATTACGTTA AGCATGTAAT
5401 AATTAACATG TAATGCATGA CGTTATTTAT GAGATGGGTT TTTATGATTA GAGTCCCGCA
5461 ATTATACATT TAATACGCGA TAGAAAACAA AATATAGCGC GCAAACTAGG ATAAATTATC
5521 GCGCGCGGTG TCATCTATGT TACTAGATCG GGGAATTGAT CCCCCCTCGA CAGCTTGCAT
5581 GCCAGCTTGG GCTGCAGGTC GAGGCTAAAA AACTAATCGC ATTATCATCC CCTCGACGTA
5641 CTGTACATAT AACCACTGGT TTTATATACA GCAGTACTGT ACATATAACC ACTGGTTTTA
5701 TATACAGCAG TCGACGTACT GTACATATAA CCACTGGTTT TATATACAGC AGTACTGTAC
5761 ATATAACCAC TGGTTTTATA TACAGCAGTC GAGGTAAGAT TAGATATGGA TATGTATATG
5821 GATATGTATA TGGTGGTAAT GCCATGTAAT ATGCTCGACT CTAGGATCTT CGCAAGACCC
5881 TTCCTCTATA TAAGGAAGTT CATTTCATTT GGAGAGGACA CGCTGAAGCT AGTCGACTCT
5941 AGCCTCGAGG CGCGCCAAGC TATCAACAAG TTTGTACAAA AAAGCAGGCT ATGCTCGAGA
6001 TGGTGAGCAA GGGCGAGGAG CTGTTCACCG GGGTGGTGCC CATCCTGGTC GAGCTGGACG
6061 GCGACGTAAA CGGCCACAAG TTCAGCGTGT CCGGCGAGGG CGAGGGCGAT GCCACCTACG
6121 GCAAGCTGAC CCTGAAGTTC ATCTGCACCA CCGGCAAGCT GCCCGTGCCC TGGCCCACCC

6181 TCGTGACCAC CCTGACCTAC GGCGTGCAGT GCTTCAGCCG CTACCCCGAC CACATGAAGC
6241 AGCACGACTT CTTCAAGTCC GCCATGCCCG AAGGCTACGT CCAGGAGCGC ACCATCTTCT
6301 TCAAGGACGA CGGCAACTAC AAGACCCGCG CCGAGGTGAA GTTCGAGGGC GACACCCTGG
6361 TGAACCGCAT CGAGCTGAAG GGCATCGACT TCAAGGAGGA CGGCAACATC CTGGGGCACA
6421 AGCTGGAGTA CAACTACAAC AGCCACAACG TCTATATCAT GGCCGACAAG CAGAAGAACG
6481 GCATCAAGGT GAACTTCAAG ATCCGCCACA ACATCGAGGA CGGCAGCGTG CAGCTCGCCG
6541 ACCACTACCA GCAGAACACC CCCATCGGCG ACGGCCCCGT GCTGCTGCCC GACAACCACT
6601 ACCTGAGTAC TCAGTCCAAG CTGAGCAAAG ACCCCAACGA GAAGCGCGAT CACATGGTCC
6661 TGCTGGAGTT CGTGACCGCC GCCGGGATCA CTCTCGGCAT GGACGAGCTG TACAAGGTCG
6721 AGCAGAAGTT GATCTCAGAG GAGGACTTAG GCATGGCGCA ACAATCGTTG ATCTACAGTT
6781 TCGTAGCTCG CGGCACGGTG ATCCTCGTTG AGTTCACTGA TTTCAAAGGT AATTTCACCT
6841 CAATCGCTGC TCAGTGCCTC CAGAAGCTTC CGTCTTCGAA CAACAAGTTC ACCTACAACT
6901 GCGATGGCCA TACCTTCAAT GACCTTGTCG AAGATGGATT CAGTAAGTCA CTTTTCGTTT
6961 GATCTATGCA TAGGTTTTAA TCTTACACCA TTCGGTGCTT ACCTCGATTT GTTTCTAGGT
7021 TTATTTGCCT AACCTATCGA TTCGTATGGA TTTGTGCATC CTGAGTAGTC TGATTTCACT
7081 GAGGAACCGC ATGAACCAAG CTTTTAGGGT TAGATCTATA TTTATTAAAC GCAATAAATC
7141 TCTGTAGAGT CTTACCTGTA TAATCACTCA GATTTGGACA GAATCCGTGG TAGCTTTGGT
7201 TGATACTATG TGGAAGAACA CCAACATTGA TCTGAGACAG GGTTTTTACT GGTTGCTTTA
7261 TTTGCGTATA GTTTCCTAGG TCATGGGTTT TGGCTGGCTT CATTTTTCAA TGAAAATCTG
7321 GGGTCTTATT GTAAGCTTTT GTGACTATCC TTTAGTTGTT GTAGCCATAT CAATTTCATA
7381 TTAAATCAAT GTCACCTGAT TATCCCGTAA GGGAAGACAA GGAGATTGAT TTTGTTCCTG
7441 TTGTACAGCC TATTGTGTTG TTGCGGTTGA TTCTGCTGGG AGGCAAATTC CCATGTCCTT
7501 TTTGGAAAGA GTAAAAGAAG ATTTTAACAA GCGATATGGT GGTGGAAAGG CTGCAACTGC
7561 TCAAGCAAAC AGCTTGAATA AGGAGTTTGG GTACTTTTTC ATTATCTCTT CTATTTGGAT
7621 GATCTTCTCT TTTATATTCG TGGACTGACT CTTTTTGGTA ACTTTGAAGC TCTAAACTGA
7681 AAGAGCATAT GCAGTATTGC ATGGATCATC CTGATGAGAT TAGCAAGCTT GCTAAGGTGA
7741 AGGCGCAAGT GTCAGAAGTT AAGGGTGTAA TGATGGAAAA CATTGAGAAG GTTTGAATCT
7801 GACCCTTTCT GTCCTCAATG TATATATTTA CATCTATGGT TGACCCATCT GAAAGAGCTC
7861 ATCTAGTCAT ACTAAGTTAC TGTGAAATCA ATTACTAATA ATTCAATGCA TTACATCTTA
7921 CGGGAAATGT CCAGTTTACC AATGTAACAC CGCTTACATA TGCGACTCAT TATTTGCAGG
7981 TTCTTGACCG TGGTGAGAAA ATTGAGCTTT TGGTGGACAA AACCGAAAAT CTTCGCTCAC
8041 AGGTTAGACT CTCTCATACC CTTATCTCCT GCATCTATTT GCCTACACTC ACACACAAAG
8101 ATGAACATGC TTTTTGAGTT CATAGCTGAC TCATCCTATT ACCATATCTT GTGTCTAACT
8161 GAACTCAAAC TGCCACAGGC ACAAGATTTC AGAACAACAG GAACGCAGAT GAGAAGAAAG
8221 ATGTGGCTTC AGAACATGAA GATAAAACTC ATAGTGCTCG CCATCATTAT CGCACTGATT
8281 CTCATCATCG TGCTCTCAGT TTGCCATGGG TTTAAGTGTT AAACCCAGCT TTCTTGTACA
8341 AAGTGGTTGA TAATTCTTAA TTAACTAGTC GATCCAGGCC TCCCAGCTTT CGTCCGTATC
8401 ATCGGTTTCG ACAACGTTCG TCAAGTTCAA TGCATCAGTT TCATTGCCCA CACACCAGAA
8461 TCCTACTAAG TTTGAGTATT ATGGCATTGG AAAAGCTGTT TTCTTCTATC ATTTGTTCTG
8521 CTTGTAATTT ACTGTGTTCT TTCAGTTTTT GTTTTCGGAC ATCAAAATGC AAATGGATGG
8581 ATAAGAGTTA ATAAATGATA TGGTCCTTTT GTTCATTCTC AAATTATTAT TATCTGTTGT
8641 TTTTACTTTA ATGGGTTGAA TTTAAGTAAG AAAGGAACTA ACAGTGTGAT ATTAAGGTGC
8701 AATGTTAGAC ATATAAAACA GTCTTTCACC TCTCTTTGGT TATGTCTTGA ATTGGTTTGT
8761 TTCTTCACTT ATCTGTGTAA TCAAGTTTAC TATGAGTCTA TGATCAAGTA ATTATGCAAT
8821 CAAGTTAAGT ACAGTATAGG CTTTTTGTGT CGAGGGGGTA CCGAGTCGAG GAATTCACTG
8881 GCCGTCGTTT TACAACGTCG TGACTGGGAA AACCCTGGCG TTACCCAACT TAATCGCCTT
8941 GCAGCACATC CCCCTTTCGC CAGCTGGCGT AATAGCGAAG AGGCCCGCAC CGATCGCCCT
9001 TCCCAACAGT TGCGCAGCCT GAATGGCGGG TACCGAGCTC GAATTCAATT CGGCGTTAAT
9061 TCAGTACATT AAAAACGTCC GCAATGTGTT ATTAAGTTGT CTAAGCGTCA ATTTGTTTAC
9121 ACCACAATAT ATCCTGCCAC CAGCCAGCCA ACAGCTCCCC GACCGGCAGC TCGGCACAAA
9181 ATCACCACTC GATACAGGCA GCCCATCAGT CCGGGACGGC GTCAGCGGGA GAGCCGTTGT
9241 AAGGCGGCAG ACTTTGCTCA TGTTACCGAT GCTATTCGGA AGAACGGCAA CTAAGCTGCC
9301 GGGTTTGAAA CACGGATGAT CTCGCGGAGG GTAGCATGTT GATTGTAACG ATGACAGAGC

9361 GTTGCTGCCT GTGATCAATT CGGGCACGAA CCCAGTGGAC ATAAGCCTCG TTCGGTTCGT
9421 AAGCTGTAAT GCAAGTAGCG TAACTGCCGT CACGCAACTG GTCCAGAACC TTGACCGAAC
9481 GCAGCGGTGG TAACGGCGCA GTGGCGGTTT TCATGGCTTC TTGTTATGAC ATGTTTTTTT
9541 GGGGTACAGT CTATGCCTCG GGCATCCAAG CAGCAAGCGC GTTACGCCGT GGGTCGATGT
9601 TTGATGTTAT GGAGCAGCAA CGATGTTACG CAGCAGGGCA GTCGCCCTAA AACAAAGTTA
9661 AACATCATGG GGGAAGCGGT GATCGCCGAA GTATCGACTC AACTATCAGA GGTAGTTGGC
9721 GTCATCGAGC GCCATCTCGA ACCGACGTTG CTGGCCGTAC ATTTGTACGG CTCCGCAGTG
9781 GATGGCGGCC TGAAGCCACA CAGTGATATT GATTTGCTGG TTACGGTGAC CGTAAGGCTT
9841 GATGAAACAA CGCGGCGAGC TTTGATCAAC GACCTTTTGG AAACTTCGGC TTCCCCTGGA
9901 GAGAGCGAGA TTCTCCGCGC TGTAGAAGTC ACCATTGTTG TGCACGACGA CATCATTCCG
9961 TGGCGTTATC CAGCTAAGCG CGAACTGCAA TTTGGAGAAT GGCAGCGCAA TGACATTCTT
10021 GCAGGTATCT TCGAGCCAGC CACGATCGAC ATTGATCTGG CTATCTTGCT GACAAAAGCA
10081 AGAGAACATA GCGTTGCCTT GGTAGGTCCA GCGGCGGAGG AACTCTTTGA TCCGGTTCCT
10141 GAACAGGATC TATTTGAGGC GCTAAATGAA ACCTTAACGC TATGGAACTC GCCGCCCGAC
10201 TGGGCTGGCG ATGAGCGAAA TGTAGTGCTT ACGTTGTCCC GCATTTGGTA CAGCGCAGTA
10261 ACCGGCAAAA TCGCGCCGAA GGATGTCGCT GCCGACTGGG CAATGGAGCG CCTGCCGGCC
10321 CAGTATCAGC CCGTCATACT TGAAGCTAGA CAGGCTTATC TTGGACAAGA AGAAGATCGC
10381 TTGGCCTCGC GCGCAGATCA GTTGGAAGAA TTTGTCCACT ACGTGAAAGG CGAGATCACC
10441 AAGGTAGTCG GCAAATAATG TCTAGCTAGA AATTCGTTCA AGCCGACGCC GCTTCGCCGG
10501 CGTTAACTCA AGCGATTAGA TGCACTAAGC ACATAATTGC TCACAGCCAA ACTATCAGGT
10561 CAAGTCTGCT TTTATTATTT TTAAGCGTGC ATAATAAGCC CTACACAAAT TGGGAGATAT
10621 ATCATGCATG ACCAAAATCC CTTAACGTGA GTTTTCGTTC CACTGAGCGT CAGACCCCGT
10681 AGAAAAGATC AAAGGATCTT CTTGAGATCC TTTTTTTCTG CGCGTAATCT GCTGCTTGCA
10741 AACAAAAAAA CCACCGCTAC CAGCGGTGGT TTGTTTGCCG GATCAAGAGC TACCAACTCT
10801 TTTTCCGAAG GTAACTGGCT TCAGCAGAGC GCAGATACCA AATACTGTCC TTCTAGTGTA
10861 GCCGTAGTTA GGCCACCACT TCAAGAACTC TGTAGCACCG CCTACATACC TCGCTCTGCT
10921 AATCCTGTTA CCAGTGGCTG CTGCCAGTGG CGATAAGTCG TGTCTTACCG GGTTGGACTC
10981 AAGACGATAG TTACCGGATA AGGCGCAGCG GTCGGGCTGA ACGGGGGGTT CGTGCACACA
11041 GCCCAGCTTG GAGCGAACGA CCTACACCGA ACTGAGATAC CTACAGCGTG AGCTATGAGA
11101 AAGCGCCACG CTTCCCGAAG GGAGAAAGGC GGACAGGTAT CCGGTAAGCG GCAGGGTCGG
11161 AACAGGAGAG CGCACGAGGG AGCTTCCAGG GGGAAACGCC TGGTATCTTT ATAGTCCTGT
11221 CGGGTTTCGC CACCTCTGAC TTGAGCGTCG ATTTTTGTGA TGCTCGTCAG GGGGGCGGAG
11281 CCTATGGAAA AACGCCAGCA ACGCGGCCTT TTTACGGTTC CTGGCCTTTT GCTGGCCTTT
11341 TGCTCACATG TTCTTTCCTG CGTTATCCCC TGATTCTGTG GATAACCGTA TTACCGCCTT
11401 TGAGTGAGCT GATACCGCTC GCCGCAGCCG AACGACCGAG CGCAGCGAGT CAGTGAGCGA
11461 GGAAGCGGAA GAGCGCCTGA TGCGGTATTT TCTCCTTACG CATCTGTGCG GTATTTCACA
11521 CCGCATATGG TGCACTCTCA GTACAATCTG CTCTGATGCC GCATAGTTAA GCCAGTATAC
11581 ACTCCGCTAT CGCTACGTGA CTGGGTCATG GCTGCGCCCC GACACCCGCC AACACCCGCT
11641 GACGCGCCCT GACGGGCTTG TCTGCTCCCG GCATCCGCTT ACAGACAAGC TGTGACCGTC
11701 TCCGGGAGCT GCATGTGTCA GAGGTTTTCA CCGTCATCAC CGAAACGCGC GAGGCAGGGT
11761 GCCTTGATGT GGGCGCCGGC GGTCGAGTGG CGACGGCGCG GCTTGTCCGC GCCCTGGTAG
11821 ATTGCCTGGC CGTAGGCCAG CCATTTTTGA GCGGCCAGCG GCCGCGATAG GCCGACGCGA
11881 AGCGGCGGGG CGTAGGGAGC GCAGCGACCG AAGGGTAGGC GCTTTTTGCA GCTCTTCGGC
11941 TGTGCGCTGG CCAGACAGTT ATGCACAGGC CAGGCGGGTT TTAAGAGTTT TAATAAGTTT
12001 TAAAGAGTTT TAGGCGGAAA AATCGCCTTT TTTCTCTTTT ATATCAGTCA CTTACATGTG
12061 TGACCGGTTC CCAATGTACG GCTTTGGGTT CCCAATGTAC GGGTTCCGGT TCCCAATGTA
12121 CGGCTTTGGG TTCCCAATGT ACGTGCTATC CACAGGAAAG AGACCTTTTC GACCTTTTTC
12181 CCCTGCTAGG GCAATTTGCC CTAGCATCTG CTCCGTACAT TAGGAACCGG CGGATGCTTC
12241 GCCCTCGATC AGGTTGCGGT AGCGCATGAC TAGGATCGGG CCAGCCTGCC CCGCCTCCTC
12301 CTTCAAATCG TACTCCGGCA GGTCATTTGA CCCGATCAGC TTGCGCACGG TGAAACAGAA
12361 CTTCTTGAAC TCTCCGGCGC TGCCACTGCG TTCGTAGATC GTCTTGAACA ACCATCTGGC
12421 TTCTGCCTTG CCTGCGGCGC GGCGTGCCAG GCGGTAGAGA AAACGGCCGA TGCCGGGATC
12481 GATCAAAAAG TAATCGGGGT GAACCGTCAG CACGTCCGGG TTCTTGCCTT CTGTGATCTC

12541 GCGGTACATC CAATCAGCTA GCTCGATCTC GATGTACTCC GGCCGCCCGG TTTCGCTCTT
12601 TACGATCTTG TAGCGGCTAA TCAAGGCTTC ACCCTCGGAT ACCGTCACCA GGCGGCCGTT
12661 CTTGGCCTTC TTCGTACGCT GCATGGCAAC GTGCGTGGTG TTTAACCGAA TGCAGGTTTC
12721 TACCAGGTCG TCTTTCTGCT TTCCGCCATC GGCTCGCCGG CAGAACTTGA GTACGTCCGC
12781 AACGTGTGGA CGGAACACGC GGCCGGGCTT GTCTCCCTTC CCTTCCCGGT ATCGGTTCAT
12841 GGATTCGGTT AGATGGGAAA CCGCCATCAG TACCAGGTCG TAATCCCACA CACTGGCCAT
12901 GCCGGCCGGC CCTGCGGAAA CCTCTACGTG CCCGTCTGGA AGCTCGTAGC GGATCACCTC
12961 GCCAGCTCGT CGGTCACGCT TCGACAGACG GAAAACGGCC ACGTCCATGA TGCTGCGACT
13021 ATCGCGGGTG CCCACGTCAT AGAGCATCGG AACGAAAAAA TCTGGTTGCT CGTCGCCCTT
13081 GGGCGGCTTC CTAATCGACG GCGCACCGGC TGCCGGCGGT TGCCGGGATT CTTTGCGGAT
13141 TCGATCAGCG GCCGCTTGCC ACGATTCACC GGGGCGTGCT TCTGCCTCGA TGCGTTGCCG
13201 CTGGGCGGCC TGCGCGGCCT TCAACTTCTC CACCAGGTCA TCACCCAGCG CCGCGCCGAT
13261 TTGTACCGGG CCGGATGGTT TGCGACCGTC ACGCCGATTC CTCGGGCTTG GGGGTTCCAG
13321 TGCCATTGCA GGGCCGGCAG ACAACCCAGC CGCTTACGCC TGGCCAACCG CCCGTTCCTC
13381 CACACATGGG GCATTCCACG GCGTCGGTGC CTGGTTGTTC TTGATTTTCC ATGCCGCCTC
13441 CTTTAGCCGC TAAAATTCAT CTACTCATTT ATTCATTTGC TCATTTACTC TGGTAGCTGC
13501 GCGATGTATT CAGATAGCAG CTCGGTAATG GTCTTGCCTT GGCGTACCGC GTACATCTTC
13561 AGCTTGGTGT GATCCTCCGC CGGCAACTGA AAGTTGACCC GCTTCATGGC TGGCGTGTCT
13621 GCCAGGCTGG CCAACGTTGC AGCCTTGCTG CTGCGTGCGC TCGGACGGCC GGCACTTAGC
13681 GTGTTTGTGC TTTTGCTCAT TTTCTCTTTA CCTCATTAAC TCAAATGAGT TTTGATTTAA
13741 TTTCAGCGGC CAGCGCCTGG ACCTCGCGGG CAGCGTCGCC CTCGGGTTCT GATTCAAGAA
13801 CGGTTGTGCC GGCGGCGGCA GTGCCTGGGT AGCTCACGCG CTGCGTGATA CGGGACTCAA
13861 GAATGGGCAG CTCGTACCCG GCCAGCGCCT CGGCAACCTC ACCGCCGATG CGCGTGCCTT
13921 TGATCGCCCG CGACACGACA AAGGCCGCTT GTAGCCTTCC ATCCGTGACC TCAATGCGCT
13981 GCTTAACCAG CTCCACCAGG TCGGCGGTGG CCCATATGTC GTAAGGGCTT GGCTGCACCG
14041 GAATCAGCAC GAAGTCGGCT GCCTTGATCG CGGACACAGC CAAGTCCGCC GCCTGGGGCG
14101 CTCCGTCGAT CACTACGAAG TCGCGCCGGC CGATGGCCTT CACGTCGCGG TCAATCGTCG
14161 GGCGGTCGAT GCCGACAACG GTTAGCGGTT GATCTTCCCG CACGGCCGCC CAATCGCGGG
14221 CACTGCCCTG GGGATCGGAA TCGACTAACA GAACATCGGC CCCGGCGAGT TGCAGGGCGC
14281 GGGCTAGATG GGTTGCGATG GTCGTCTTGC CTGACCCGCC TTTCTGGTTA AGTACAGCGA
14341 TAACCTTCAT GCGTTCCCCT TGCGTATTTG TTTATTTACT CATCGCATCA TATACGCAGC
14401 GACCGCATGA CGCAAGCTGT TTTACTCAAA TACACATCAC CTTTTTAGAC GGCGGCGCTC
14461 GGTTTCTTCA GCGGCCAAGC TGGCCGGCCA GGCCGCCAGC TTGGCATCAG ACAAACCGGC
14521 CAGGATTTCA TGCAGCCGCA CGGTTGAGAC GTGCGCGGGC GGCTCGAACA CGTACCCGGC
14581 CGCGATCATC TCCGCCTCGA TCTCTTCGGT AATGAAAAAC GGTTCGTCCT GGCCGTCCTG
14641 GTGCGGTTTC ATGCTTGTTC CTCTTGGCGT TCATTCTCGG CGGCCGCCAG GGCGTCGGCC
14701 TCGGTCAATG CGTCCTCACG GAAGGCACCG CGCCGCCTGG CCTCGGTGGG CGTCACTTCC
14761 TCGCTGCGCT CAAGTGCGCG GTACAGGGTC GAGCGATGCA CGCCAAGCAG TGCAGCCGCC
14821 TCTTTCACGG TGCGGCCTTC CTGGTCGATC AGCTCGCGGG CGTGCGCGAT CTGTGCCGGG
14881 GTGAGGGTAG GGCGGGGGCC AAACTTCACG CCTCGGGCCT TGGCGGCCTC GCGCCCGCTC
14941 CGGGTGCGGT CGATGATTAG GGAACGCTCG AACTCGGCAA TGCCGGCGAA CACGGTCAAC
15001 ACCATGCGGC CGGCCGGCGT GGTGGTGTCG GCCCACGGCT CTGCCAGGCT ACGCAGGCCC
15061 GCGCCGGCCT CCTGGATGCG CTCGGCAATG TCCAGTAGGT CGCGGGTGCT GCGGGCCAGG
15121 CGGTCTAGCC TGGTCACTGT CACAACGTCG CCAGGGCGTA GGTGGTCAAG CATCCTGGCC
15181 AGCTCCGGGC GGTCGCGCCT GGTGCCGGTG ATCTTCTCGG AAAACAGCTT GGTGCAGCCG
15241 GCCGCGTGCA GTTCGGCCCG TTGGTTGGTC AAGTCCTGGT CGTCGGTGCT GACGCGGGCA
15301 TAGCCCAGCA GGCCAGCGGC GGCGCTCTTG TTCATGGCGT AATGTCTCCG GTTCTAGTCG
15361 CAAGTATTCT ACTTTATGCG ACTAAAACAC GCGACAAGAA AACGCCAGGA AAAGGGCAGG
15421 GCGGCAGCCT GTCGCGTAAC TTAGGACTTG TGCGACATGT CGTTTTCAGA AGACGGCTGC
15481 ACTGAACGTC AGAAGCCGAC TGCACTATAG CAGCGGAGGG GTTGGATCAA AGTACTTTGA
15541 TCCCGAGGGG AACCCTGTGG TTGGCATGCA CATACAAATG GACGAACGGA TAAACCTTTT
15601 CACGCCCTTT TAAATATCCG TTATTCTAAT AAACGCTCTT TTCTCTTAG

//

```
LOCUS       D934     15675 bp    DNA     circular UNA 10-DEC-2018
COMMENT     Insert from E445_pDONR207_GFP_myc_gVamp723 414 to 2798
FEATURES             Location/Qualifiers
     misc_feature    1..26
                     /vntifkey="21"
                     /locus_tag="RB"
     promoter        44..1640
                     /vntifkey="29"
                     /locus_tag="5' Promoter-UBQ10"
     5'UTR           1530..2029
                     /vntifkey="52"
                     /locus_tag="5'UTR"
     misc_feature    1641..1725
                     /vntifkey="21"
                     /locus_tag="Exon 1.1"
     intron          1726..2029
                     /vntifkey="15"
                     /locus_tag="Intron 1.1"
     CDS             2037..3921
                     /vntifkey="4"
                     /locus_tag="XVE"
     promoter        3937..4269
                     /vntifkey="29"
                     /locus_tag="NOS promoter"
     CDS             3938..5573
                     /vntifkey="4"
                     /locus_tag="hygromycin resistance"
     promoter        5576..5853
                     /vntifkey="29"
                     /locus_tag="lexA -46 35S promoter"
     repeat_region   5961..5973
                     /vntifkey="34"
                     /locus_tag="attR1"
     CDS             5997..5999
                     /locus_tag="MYC(1)"
     misc_feature    6000..6716
                     /locus_tag="XhoI-meGFP-SalI"
     CDS             6720..6749
                     /locus_tag="MYC"
     misc_feature    6753..8348
                     /locus_tag="gVamp723"
     misc_feature    7820..7841
                     /locus_tag="g723 seq primer"
     misc_feature    8359..8444
                     /vntifkey="21"
                     /locus_tag="confirmed sequence"
     repeat_region   8359..8373
                     /vntifkey="34"
                     /locus_tag="attR2"
     terminator      8401..8880
                     /vntifkey="43"
                     /locus_tag="T3A"
     misc_feature    9141..9165
```

/vntifkey="21"
/locus_tag="LB"

ORIGIN

1 GTTTACCCGC CAATATATCC TGTCAAACAC TGATAGTTTA AACAGTCTAG CTCAACAGAG
61 CTTTTAACCC AAATTGGTAC AATAGAATAC AACTTTAGAT CATAATTCTC AAAAGAAAGA
121 GATTCCTTAG CTATTCTATC TGCCACTCCA TTTCCTTCTC GGCTTGTATG CACAAGCATA
181 AAATCCTCAA ACTTGCTAAG TAGATACTTT ATGTCTTGGA TAATTGGATT GAGACTTGAC
241 AAGCATAACT TTCATGTAAC CAAAGACACA AGTTGCTGAG AATCCACCTC AAAAATGATC
301 TTCCTATAAT TGAATCGGGA TAATGACAGC ACAGCCCATC TAAGAGCCTC CACTTCTACT
361 TCCAGCACGC TTCTTACTTT TACCACAGCT CTTGCACCTA ACCATAACAC CTTCCCTGTA
421 TGATCGCGAA GCACCCACCC TAAGCCACAT TTTAATCCTT CTGTTGGCCA TGCCCCATCA
481 AAGTTGCACT TAACCCAAGA TTGTGGTGGA GCTTCCCATG TTTCTCGTCT GTCCCGACGG
541 TGTTGTGGTT GGTGCTTTCC TTACATTCTG AGCCTCTTTC CTTCTAATCC ACTCATCTGC
601 ATCTTCTTGT GTCCTTACTA ATACCTCATT GGTTCCAAAT TCCCTCCCTT TAAGCACCAG
661 CTCGTTTCTG TTCTTCCACA GCCTCCCAAG TATCCAAGGG ACTAAAGCCT CCACATTCTT
721 CAGATCAGGA TATTCTTGTT TAAGATGTTG AACTCTATGG AGGTTTGTAT GAACTGATGA
781 TCTAGGACCG GATAAGTTCC CTTCTTCATA GCGAACTTAT TCAAAGAATG TTTTGTGTAT
841 CATTCTTGTT ACATTGTTAT TAATGAAAAA ATATTATTGG TCATTGGACT GAACACGAGT
901 GTTAAATATG GACCAGGCCC CAAATAAGAT CCATTGATAT ATGAATTAAA TAACAAGAAT
961 AAATCGAGTC ACCAAACCAC TTGCCTTTTT TAACGAGACT TGTTCACCAA CTTGATACAA
1021 AAGTCATTAT CCTATGCAAA TCAATAATCA TACAAAAATA TCCAATAACA CTAAAAAATT
1081 AAAAGAAATG GATAATTTCA CAATATGTTA TACGATAAAG AAGTTACTTT TCCAAGAAAT
1141 TCACTGATTT TATAAGCCCA CTTGCATTAG ATAAATGGCA AAAAAAAACA AAAAGGAAAA
1201 GAAATAAAGC ACGAAGAATT CTAGAAAATA CGAAATACGC TTCAATGCAG TGGGACCCAC
1261 GGTTCAATTA TTGCCAATTT TCAGCTCCAC CGTATATTTA AAAAATAAAA CGATAATGCT
1321 AAAAAAATAT AAATCGTAAC GATCGTTAAA TCTCAACGGC TGGATCTTAT GACGACCGTT
1381 AGAAATTGTG GTTGTCGACG AGTCAGTAAT AAACGGCGTC AAAGTGGTTG CAGCCGGCAC
1441 ACACGAGTCG TGTTTATCAA CTCAAAGCAC AAATACTTTT CCTCAACCTA AAAATAAGGC
1501 AATTAGCCAA AAACAACTTT GCGTGTAAAC AACGCTCAAT ACACGTGTCA TTTTATTATT
1561 AGCTATTGCT TCACCGCCTT AGCTTTCTCG TGACCTAGTC GTCCTCGTCT TTTCTTCTTC
1621 TTCTTCTATA AAACAATACC CAAAGAGCTC TTCTTCTTCA CAATTCAGAT TTCAATTTCT
1681 CAAAATCTTA AAAACTTTCT CTCAATTCTC TCTACCGTGA TCAAGGTAAA TTTCTGTGTT
1741 CCTTATTCTC TCAAAATCTT CGATTTTGTT TTCGTTCGAT CCCAATTTCG TATATGTTCT
1801 TTGGTTTAGA TTCTGTTAAT CTTAGATCGA AGACGATTTT CTGGGTTTGA TCGTTAGATA
1861 TCATCTTAAT TCTCGATTAG GGTTTCATAG ATATCATCCG ATTTGTTCAA ATAATTTGAG
1921 TTTTGTCGAA TAATTACTCT TCGATTTGTG ATTTCTATCT AGATCTGGTG TTAGTTTCTA
1981 GTTTGTGCGA TCGAATTTGT CGATTAATCT GAGTTTTTCT GATTAACAGT TCGAAATGAA
2041 AGCGTTAACG GCCAGGCAAC AAGAGGTGTT TGATCTCATC CGTGATCACA TCAGCCAGAC
2101 AGGTATGCCG CCGACGCGTG CGGAAATCGC GCAGCGTTTG GGGTTCCGTT CCCCAAACGC
2161 GGCTGAAGAA CATCTGAAGG CGCTGGCACG CAAAGGCGTT ATTGAAATTG TTTCCGGCGC
2221 ATCACGCGGG ATTCGTCTGT TGCAGGAAGA GGAAGAAGGG TTGCCGCTGG TAGGTCGTGT
2281 GGCTGCCGGT GAACCGTCGA GCGCCCCCCC GACCGATGTC AGCCTGGGGG ACGAGCTCCA
2341 CTTAGACGGC GAGGACGTGG CGATGGCGCA TGCCGACGCG CTAGACGATT TCGATCTGGA
2401 CATGTTGGGG GACGGGGATT CCCCGGGTCC GGGATTTACC CCCCACGACT CCGCCCCCTA
2461 CGGCGCTCTG GATATGGCCG ACTTCGAGTT TGAGCAGATG TTTACCGATG CCCTTGGAAT
2521 TGACGAGTAC GGTGGGGATC CGTCTGCTGG AGACATGAGA GCTGCCAACC TTTGGCCAAG
2581 CCCGCTCATG ATCAAACGCT CTAAGAAGAA CAGCCTGGCC TTGTCCCTGA CGGCCGACCA
2641 GATGGTCAGT GCCTTGTTGG ATGCTGAGCC CCCCATACTC TATTCCGAGT ATGATCCTAC
2701 CAGACCCTTC AGTGAAGCTT CGATGATGGG CTTACTGACC AACCTGGCAG ACAGGGAGCT
2761 GGTTCACATG ATCAACTGGG CGAAGAGGGT GCCAGGCTTT GTGGATTTGA CCCTCCATGA
2821 TCAGGTCCAC CTTCTAGAAT GTGCCTGGCT AGAGATCCTG ATGATTGGTC TCGTCTGGCG
2881 CTCCATGGAG CACCCAGTGA AGCTACTGTT TGCTCCTAAC TTGCTCTTGG ACAGGAACCA
2941 GGGAAAATGT GTAGAGGGCA TGGTGGAGAT CTTCGACATG CTGCTGGCTA CATCATCTCG

3001 GTTCCGCATG ATGAATCTGC AGGGAGAGGA GTTTGTGTGC CTCAAATCTA TTATTTTGCT
3061 TAATTCTGGA GTGTACACAT TTCTGTCCAG CACCCTGAAG TCTCTGGAAG AGAAGGACCA
3121 TATCCACCGA GTCCTGGACA AGATCACAGA CACTTTGATC CACCTGATGG CCAAGGCAGG
3181 CCTGACCCTG CAGCAGCAGC ACCAGCGGCT GGCCCAGCTC CTCCTCATCC TCTCCCACAT
3241 CAGGCACATG AGTAACAAAG GCATGGAGCA TCTGTACAGC ATGAAGTGCA AGAACGTGGT
3301 GCCCCTCTAT GACCTGCTGC TGGAGATGCT GGACGCCCAC CGCCTACATG CGCCCACTAG
3361 CCGTGGAGGG GCATCCGTGG AGGAGACGGA CCAAAGCCAC TTGGCCACTG CGGGCTCTAC
3421 TTCATCGCAT TCCTTGCAAA AGTATTACAT CACGGGGGAG GCAGAGGGTT TCCCTGCCAC
3481 AGTCTGAGAG CTCCCTGGCG AATTCCCAGA GATGTTAGCT GAAATCATCA CTAATCAGAT
3541 ACCAAAATAT TCAAATGGAA ATATCAAAAA GCTTCTGTTT CATCAAAAAT GACTCGACCT
3601 AACTGAGTAA GCTAGCTTGT TCGAGTATTA TGGCATTGGG AAAACTGTTT TTCTTGTACC
3661 ATTTGTTGTG CTTGTAATTT ACTGTGTTTT TTATTCGGTT TTCGCTATCG AACTGTGAAA
3721 TGGAAATGGA TGGAGAAGAG TTAATGAATG ATATGGTCCT TTTGTTCATT CTCAAATTAA
3781 TATTATTTGT TTTTTCTCTT ATTTGTTGTG TGTTGAATTT GAAATTATAA GAGATATGCA
3841 AACATTTTGT TTTGAGTAAA AATGTGTCAA ATCGTGGCCT CTAATGACCG AAGTTAATAT
3901 GAGGAGTAAA ACATCCCAAA CAAGCTTGGA AACTGAAGGC GGGAAACGAC AATCTGATCA
3961 TGAGCGGAGA ATTAAGGGAG TCACGTTATG ACCCCCGCCG ATGACGCGGG ACAAGCCGTT
4021 TTACGTTTGG AACTGACAGA ACCGCAACGA TTGAAGGAGC CACTCAGCCG CGGGTTTCTG
4081 GAGTTTAATG AGCTAAGCAC ATACGTCAGA AACCATTATT GCGCGTTCAA AAGTCGCCTA
4141 AGGTCACTAT CAGCTAGCAA ATATTTCTTG TCAAAAATGC TCCACTGACG TTCCATAAAT
4201 TCCCCTCGGT ATCCAATTAG AGTCTCATAT TCACTCTCAA TCCAAATAAT CTGCACCGGA
4261 TCCCCTAGAA TGAAAAAGCC TGAACTCACC GCGACGTCTG TCGAGAAGTT TCTGATCGAA
4321 AAGTTCGACA GCGTCTCCGA CCTGATGCAG CTCTCGGAGG GCGAAGAATC TCGTGCTTTC
4381 AGCTTCGATG TAGGAGGGCG TGGATATGTC CTGCGGGTAA ATAGCTGCGC CGATGGTTTC
4441 TACAAAGATC GTTATGTTTA TCGGCACTTT GCATCGGCCG CGCTCCCGAT TCCGGAAGTG
4501 CTTGACATTG GGGAATTCAG CGAGAGCCTG ACCTATTGCA TCTCCCGCCG TGCACAGGGT
4561 GTCACGTTGC AAGACCTGCC TGAAACCGAA CTGCCCGCTG TTCTGCAGCC GGTCGCGGAG
4621 GCCATGGATG CGATCGCTGC GGCCGATCTT AGCCAGACGA GCGGGTTCGG CCCATTCGGA
4681 CCGCAAGGAA TCGGTCAATA CACTACATGG CGTGATTTCA TATGCGCGAT TGCTGATCCC
4741 CATGTGTATC ACTGGCAAAC TGTGATGGAC GACACCGTCA GTGCGTCCGT CGCGCAGGCT
4801 CTCGATGAGC TGATGCTTTG GGCCGAGGAC TGCCCCGAAG TCCGGCACCT CGTGCACGCG
4861 GATTTCGGCT CCAACAATGT CCTGACGGAC AATGGCCGCA TAACAGCGGT CATTGACTGG
4921 AGCGAGGCGA TGTTCGGGGA TTCCCAATAC GAGGTCGCCA ACATCTTCTT CTGGAGGCCG
4981 TGGTTGGCTT GTATGGAGCA GCAGACGCGC TACTTCGAGC GGAGGCATCC GGAGCTTGCA
5041 GGATCGCCGC GGCTCCGGGC GTATATGCTC CGCATTGGTC TTGACCAACT CTATCAGAGC
5101 TTGGTTGACG GCAATTTCGA TGATGCAGCT TGGGCGCAGG GTCGATGCGA CGCAATCGTC
5161 CGATCCGGAG CCGGGACTGT CGGGCGTACA CAAATCGCCC GCAGAAGCGC GGCCGTCTGG
5221 ACCGATGGCT GTGTAGAAGT ACTCGCCGAT AGTGGAAACC GACGCCCCAG CACTCGTCCG
5281 AGGGCAAAGG AATAGCGATC GTTCAAACAT TTGGCAATAA AGTTTCTTAA GATTGAATCC
5341 TGTTGCCGGT CTTGCGATGA TTATCATATA ATTTCTGTTG AATTACGTTA AGCATGTAAT
5401 AATTAACATG TAATGCATGA CGTTATTTAT GAGATGGGTT TTTATGATTA GAGTCCCGCA
5461 ATTATACATT TAATACGCGA TAGAAAACAA AATATAGCGC GCAAACTAGG ATAAATTATC
5521 GCGCGCGGTG TCATCTATGT TACTAGATCG GGGAATTGAT CCCCCCTCGA CAGCTTGCAT
5581 GCCAGCTTGG GCTGCAGGTC GAGGCTAAAA AACTAATCGC ATTATCATCC CCTCGACGTA
5641 CTGTACATAT AACCACTGGT TTTATATACA GCAGTACTGT ACATATAACC ACTGGTTTTA
5701 TATACAGCAG TCGACGTACT GTACATATAA CCACTGGTTT TATATACAGC AGTACTGTAC
5761 ATATAACCAC TGGTTTTATA TACAGCAGTC GAGGTAAGAT TAGATATGGA TATGTATATG
5821 GATATGTATA TGGTGGTAAT GCCATGTAAT ATGCTCGACT CTAGGATCTT CGCAAGACCC
5881 TTCCTCTATA TAAGGAAGTT CATTTCATTT GGAGAGGACA CGCTGAAGCT AGTCGACTCT
5941 AGCCTCGAGG CGCGCCAAGC TATCAACAAG TTTGTACAAA AAAGCAGGCT ATGCTCGAGA
6001 TGGTGAGCAA GGGCGAGGAG CTGTTCACCG GGGTGGTGCC CATCCTGGTC GAGCTGGACG
6061 GCGACGTAAA CGGCCACAAG TTCAGCGTGT CCGGCGAGGG CGAGGGCGAT GCCACCTACG
6121 GCAAGCTGAC CCTGAAGTTC ATCTGCACCA CCGGCAAGCT GCCCGTGCCC TGGCCCACCC

6181 TCGTGACCAC CCTGACCTAC GGCGTGCAGT GCTTCAGCCG CTACCCCGAC CACATGAAGC
6241 AGCACGACTT CTTCAAGTCC GCCATGCCCG AAGGCTACGT CCAGGAGCGC ACCATCTTCT
6301 TCAAGGACGA CGGCAACTAC AAGACCCGCG CCGAGGTGAA GTTCGAGGGC GACACCCTGG
6361 TGAACCGCAT CGAGCTGAAG GGCATCGACT TCAAGGAGGA CGGCAACATC CTGGGGCACA
6421 AGCTGGAGTA CAACTACAAC AGCCACAACG TCTATATCAT GGCCGACAAG CAGAAGAACG
6481 GCATCAAGGT GAACTTCAAG ATCCGCCACA ACATCGAGGA CGGCAGCGTG CAGCTCGCCG
6541 ACCACTACCA GCAGAACACC CCCATCGGCG ACGGCCCCGT GCTGCTGCCC GACAACCACT
6601 ACCTGAGTAC TCAGTCCAAG CTGAGCAAAG ACCCCAACGA GAAGCGCGAT CACATGGTCC
6661 TGCTGGAGTT CGTGACCGCC GCCGGGATCA CTCTCGGCAT GGACGAGCTG TACAAGGTCG
6721 AGCAGAAGTT GATCTCAGAG GAGGACTTAG GCATGGCGCA ACAATCGTTG TTCTACAGTT
6781 TCATCGCTCG CGGCACCGTA ATCCTCGTCG AGTTCACAGA TTTCAAAGGC AATTTCACAT
6841 CTGTCGCTGC TCAGTACCTT GAGAATCTTC CTTCCTCGAA CAACAAGTTT ACCTACAACT
6901 GCGATGGTCA TACGTTCAAC GACCTCGTCG AAAATGGATT CAGTGAGTCA AAATATTGCT
6961 CGTGATGTGT TTGTGATTGT GTTTTCGATT AGTCGATTTA TTCATTGTTT GGATTAGATT
7021 TCTTTTTACC TAATTGAGCA TTTGAGAATC GAGTTCTTAT GTTTGGTATA TCTCTATGAC
7081 TTATCTGAGT CGAGTTCTTA TGTGTTTACC GTTTGGTATA ATTTCAGGAA TAGTTATGTA
7141 TTGGGTTTGA TAATTAATTG ATGTTAGATC TATAATCAGT GATATGTTTG GTGGATCTAT
7201 TTAAGCATCT ATGCAATTCT TATTATGAAT CTATGATTCT TGGCTTCTTT GCGTGTAGCT
7261 TTTGTTTTCA TGAATTTGGC TGGTTTTTCA AATGAGAGTT GTTTGGGAAA TTTGTGGATA
7321 TATATCGCTC GAGTAAATTG CATTGGTGTA TTGTGTTTAT TATATTGCTC TGTTTTAATT
7381 AAACTTAGTT GGGTTTTGAT GACGTTGGAT ATGGTCCATA TCATATGGAC TCTTTATTAT
7441 ATTTTTTAAG TTTCTTATGA GGGATATGTT TGTGCAGCCT ATTGTGTTGT TGCAGTTGAT
7501 TCTGCTGGGA GGGAGATTCC TATGGCTTTC TTGGAACGCG TGAAGGAGGA TTTTTATAAG
7561 AGATATGGTG GTGAAAAGGC TGCAACTGAT CAAGCAAATA GCTTGAATAA AGAATTTGGG
7621 TATGTGTTTT TTGATCTATA TGATTGGTAT TGAGAAGTTT GTATTATTCA TGTCAATGAC
7681 AGACTCCTTT TTGGCTGCTC TAAAGGTCGA ATCTGAAAGA GCACATGCAG TATTGCATGG
7741 ATCATCCTGA TGAGATTAGC AACCTTGCTA AAGCTAAAGC TCAAGTGTCT GAAGTTAAAA
7801 GTTTAATGAT GGAAAACATT GAGAAGGTTC GATTCTGACC CTTTTTTCTG TTAATGGTAT
7861 AATTTTATGT TTTAATCTAT TTCCAGTAGA GTTAAGCTCG TATGAGTTAA GTGTCATGAT
7921 CTTTTGGTTC CTTAACCTTT AAATTGATGA GGGGAATGTC CAGTTCTGGG TTGTATAGGC
7981 ATGTATGCTG ACCGTTTTTA CATTGTGGAA CACATTCATT TGCAGGTTCT TGCCCGTGGT
8041 GTGATATGTG AGATGCTGGG TAGTTCAGAG GTTAGTTCTC TTAGGCATAT GGTTCTAAGT
8101 TCCATACACA TAGATATGCT TGAGTTGCCA GTTCTCTCTT ACTAATGTTG TCTTGGGTTT
8161 CCAACTTTTA ATTGAACTCG AAACTGTCAC AGTCACAGCC GCAAGCTTTC TATATAAAAA
8221 GAACTCAAAT GAAAAGGAAG AAGTGGTTTC AGAACATGAA GATAAAACTC ATTGTCCTTG
8281 CAATTATCAT TGCCTTGATT CTCATCATCA TCCTCTCGGT TTGTGGGGGA TTCAACTGCG
8341 GTAAATAAAC CCAGCTTTCT TGTACAAAGT GGTTGATAAT TCTTAATTAA CTAGTCGATC
8401 CAGGCCTCCC AGCTTTCGTC CGTATCATCG GTTTCGACAA CGTTCGTCAA GTTCAATGCA
8461 TCAGTTTCAT TGCCCACACA CCAGAATCCT ACTAAGTTTG AGTATTATGG CATTGGAAAA
8521 GCTGTTTTCT TCTATCATTT GTTCTGCTTG TAATTTACTG TGTTCTTTCA GTTTTTGTTT
8581 TCGGACATCA AAATGCAAAT GGATGGATAA GAGTTAATAA ATGATATGGT CCTTTTGTTC
8641 ATTCTCAAAT TATTATTATC TGTTGTTTTT ACTTTAATGG GTTGAATTTA AGTAAGAAAG
8701 GAACTAACAG TGTGATATTA AGGTGCAATG TTAGACATAT AAAACAGTCT TTCACCTCTC
8761 TTTGGTTATG TCTTGAATTG GTTTGTTTCT TCACTTATCT GTGTAATCAA GTTTACTATG
8821 AGTCTATGAT CAAGTAATTA TGCAATCAAG TTAAGTACAG TATAGGCTTT TTGTGTCGAG
8881 GGGGTACCGA GTCGAGGAAT TCACTGGCCG TCGTTTTACA ACGTCGTGAC TGGGAAAACC
8941 CTGGCGTTAC CCAACTTAAT CGCCTTGCAG CACATCCCCC TTTCGCCAGC TGGCGTAATA
9001 GCGAAGAGGC CCGCACCGAT CGCCCTTCCC AACAGTTGCG CAGCCTGAAT GGCGGGTACC
9061 GAGCTCGAAT TCAATTCGGC GTTAATTCAG TACATTAAAA ACGTCCGCAA TGTGTTATTA
9121 AGTTGTCTAA GCGTCAATTT GTTTACACCA CAATATATCC TGCCACCAGC CAGCCAACAG
9181 CTCCCCGACC GGCAGCTCGG CACAAAATCA CCACTCGATA CAGGCAGCCC ATCAGTCCGG
9241 GACGGCGTCA GCGGGAGAGC CGTTGTAAGG CGGCAGACTT TGCTCATGTT ACCGATGCTA
9301 TTCGGAAGAA CGGCAACTAA GCTGCCGGGT TTGAAACACG GATGATCTCG CGGAGGGTAG

9361 CATGTTGATT GTAACGATGA CAGAGCGTTG CTGCCTGTGA TCAATTCGGG CACGAACCCA
9421 GTGGACATAA GCCTCGTTCG GTTCGTAAGC TGTAATGCAA GTAGCGTAAC TGCCGTCACG
9481 CAACTGGTCC AGAACCTTGA CCGAACGCAG CGGTGGTAAC GGCGCAGTGG CGGTTTTCAT
9541 GGCTTCTTGT TATGACATGT TTTTTTGGGG TACAGTCTAT GCCTCGGGCA TCCAAGCAGC
9601 AAGCGCGTTA CGCCGTGGGT CGATGTTTGA TGTTATGGAG CAGCAACGAT GTTACGCAGC
9661 AGGGCAGTCG CCCTAAAACA AAGTTAAACA TCATGGGGGA AGCGGTGATC GCCGAAGTAT
9721 CGACTCAACT ATCAGAGGTA GTTGGCGTCA TCGAGCGCCA TCTCGAACCG ACGTTGCTGG
9781 CCGTACATTT GTACGGCTCC GCAGTGGATG GCGGCCTGAA GCCACACAGT GATATTGATT
9841 TGCTGGTTAC GGTGACCGTA AGGCTTGATG AAACAACGCG GCGAGCTTTG ATCAACGACC
9901 TTTTGGAAAC TTCGGCTTCC CCTGGAGAGA GCGAGATTCT CCGCGCTGTA GAAGTCACCA
9961 TTGTTGTGCA CGACGACATC ATTCCGTGGC GTTATCCAGC TAAGCGCGAA CTGCAATTTG
10021 GAGAATGGCA GCGCAATGAC ATTCTTGCAG GTATCTTCGA GCCAGCCACG ATCGACATTG
10081 ATCTGGCTAT CTTGCTGACA AAAGCAAGAG AACATAGCGT TGCCTTGGTA GGTCCAGCGG
10141 CGGAGGAACT CTTTGATCCG GTTCCTGAAC AGGATCTATT TGAGGCGCTA AATGAAACCT
10201 TAACGCTATG GAACTCGCCG CCCGACTGGG CTGGCGATGA GCGAAATGTA GTGCTTACGT
10261 TGTCCCGCAT TTGGTACAGC GCAGTAACCG GCAAAATCGC GCCGAAGGAT GTCGCTGCCG
10321 ACTGGGCAAT GGAGCGCCTG CCGGCCCAGT ATCAGCCCGT CATACTTGAA GCTAGACAGG
10381 CTTATCTTGG ACAAGAAGAA GATCGCTTGG CCTCGCGCGC AGATCAGTTG GAAGAATTTG
10441 TCCACTACGT GAAAGGCGAG ATCACCAAGG TAGTCGGCAA ATAATGTCTA GCTAGAAATT
10501 CGTTCAAGCC GACGCCGCTT CGCCGGCGTT AACTCAAGCG ATTAGATGCA CTAAGCACAT
10561 AATTGCTCAC AGCCAAACTA TCAGGTCAAG TCTGCTTTTA TTATTTTTAA GCGTGCATAA
10621 TAAGCCCTAC ACAAATTGGG AGATATATCA TGCATGACCA AAATCCCTTA ACGTGAGTTT
10681 TCGTTCCACT GAGCGTCAGA CCCCGTAGAA AAGATCAAAG GATCTTCTTG AGATCCTTTT
10741 TTTCTGCGCG TAATCTGCTG CTTGCAAACA AAAAAACCAC CGCTACCAGC GGTGGTTTGT
10801 TTGCCGGATC AAGAGCTACC AACTCTTTTT CCGAAGGTAA CTGGCTTCAG CAGAGCGCAG
10861 ATACCAAATA CTGTCCTTCT AGTGTAGCCG TAGTTAGGCC ACCACTTCAA GAACTCTGTA
10921 GCACCGCCTA CATACCTCGC TCTGCTAATC CTGTTACCAG TGGCTGCTGC CAGTGGCGAT
10981 AAGTCGTGTC TTACCGGGTT GGACTCAAGA CGATAGTTAC CGGATAAGGC GCAGCGGTCG
11041 GGCTGAACGG GGGGTTCGTG CACACAGCCC AGCTTGGAGC GAACGACCTA CACCGAACTG
11101 AGATACCTAC AGCGTGAGCT ATGAGAAAGC GCCACGCTTC CCGAAGGGAG AAAGGCGGAC
11161 AGGTATCCGG TAAGCGGCAG GGTCGGAACA GGAGAGCGCA CGAGGGAGCT TCCAGGGGGA
11221 AACGCCTGGT ATCTTTATAG TCCTGTCGGG TTTCGCCACC TCTGACTTGA GCGTCGATTT
11281 TTGTGATGCT CGTCAGGGGG GCGGAGCCTA TGGAAAAACG CCAGCAACGC GGCCTTTTTA
11341 CGGTTCCTGG CCTTTTGCTG GCCTTTTGCT CACATGTTCT TTCCTGCGTT ATCCCCTGAT
11401 TCTGTGGATA ACCGTATTAC CGCCTTTGAG TGAGCTGATA CCGCTCGCCG CAGCCGAACG
11461 ACCGAGCGCA GCGAGTCAGT GAGCGAGGAA GCGGAAGAGC GCCTGATGCG GTATTTTCTC
11521 CTTACGCATC TGTGCGGTAT TTCACACCGC ATATGGTGCA CTCTCAGTAC AATCTGCTCT
11581 GATGCCGCAT AGTTAAGCCA GTATACACTC CGCTATCGCT ACGTGACTGG GTCATGGCTG
11641 CGCCCCGACA CCCGCCAACA CCCGCTGACG CGCCCTGACG GGCTTGTCTG CTCCCGGCAT
11701 CCGCTTACAG ACAAGCTGTG ACCGTCTCCG GGAGCTGCAT GTGTCAGAGG TTTTCACCGT
11761 CATCACCGAA ACGCGCGAGG CAGGGTGCCT TGATGTGGGC GCCGGCGGTC GAGTGGCGAC
11821 GGCGCGGCTT GTCCGCGCCC TGGTAGATTG CCTGGCCGTA GGCCAGCCAT TTTTGAGCGG
11881 CCAGCGGCCG CGATAGGCCG ACGCGAAGCG GCGGGGCGTA GGGAGCGCAG CGACCGAAGG
11941 GTAGGCGCTT TTTGCAGCTC TTCGGCTGTG CGCTGGCCAG ACAGTTATGC ACAGGCCAGG
12001 CGGGTTTTAA GAGTTTTAAT AAGTTTTAAA GAGTTTTAGG CGGAAAAATC GCCTTTTTTC
12061 TCTTTTATAT CAGTCACTTA CATGTGTGAC CGGTTCCCAA TGTACGGCTT TGGGTTCCCA
12121 ATGTACGGGT TCCGGTTCCC AATGTACGGC TTTGGGTTCC CAATGTACGT GCTATCCACA
12181 GGAAAGAGAC CTTTTCGACC TTTTTCCCCT GCTAGGGCAA TTTGCCCTAG CATCTGCTCC
12241 GTACATTAGG AACCGGCGGA TGCTTCGCCC TCGATCAGGT TGCGGTAGCG CATGACTAGG
12301 ATCGGGCCAG CCTGCCCCGC CTCCTCCTTC AAATCGTACT CCGGCAGGTC ATTTGACCCG
12361 ATCAGCTTGC GCACGGTGAA ACAGAACTTC TTGAACTCTC CGGCGCTGCC ACTGCGTTCG
12421 TAGATCGTCT TGAACAACCA TCTGGCTTCT GCCTTGCCTG CGGCGCGGCG TGCCAGGCGG
12481 TAGAGAAAAC GGCCGATGCC GGGATCGATC AAAAAGTAAT CGGGGTGAAC CGTCAGCACG

12541 TCCGGGTTCT TGCCTTCTGT GATCTCGCGG TACATCCAAT CAGCTAGCTC GATCTCGATG
12601 TACTCCGGCC GCCCGGTTTC GCTCTTTACG ATCTTGTAGC GGCTAATCAA GGCTTCACCC
12661 TCGGATACCG TCACCAGGCG GCCGTTCTTG GCCTTCTTCG TACGCTGCAT GGCAACGTGC
12721 GTGGTGTTTA ACCGAATGCA GGTTTCTACC AGGTCGTCTT TCTGCTTTCC GCCATCGGCT
12781 CGCCGGCAGA ACTTGAGTAC GTCCGCAACG TGTGGACGGA ACACGCGGCC GGGCTTGTCT
12841 CCCTTCCCTT CCCGGTATCG GTTCATGGAT TCGGTTAGAT GGGAAACCGC CATCAGTACC
12901 AGGTCGTAAT CCCACACACT GGCCATGCCG GCCGGCCCTG CGGAAACCTC TACGTGCCCG
12961 TCTGGAAGCT CGTAGCGGAT CACCTCGCCA GCTCGTCGGT CACGCTTCGA CAGACGGAAA
13021 ACGGCCACGT CCATGATGCT GCGACTATCG CGGGTGCCCA CGTCATAGAG CATCGGAACG
13081 AAAAAATCTG GTTGCTCGTC GCCCTTGGGC GGCTTCCTAA TCGACGGCGC ACCGGCTGCC
13141 GGCGGTTGCC GGGATTCTTT GCGGATTCGA TCAGCGGCCG CTTGCCACGA TTCACCGGGG
13201 CGTGCTTCTG CCTCGATGCG TTGCCGCTGG GCGGCCTGCG CGGCCTTCAA CTTCTCCACC
13261 AGGTCATCAC CCAGCGCCGC GCCGATTTGT ACCGGGCCGG ATGGTTTGCG ACCGTCACGC
13321 CGATTCCTCG GGCTTGGGGG TTCCAGTGCC ATTGCAGGGC CGGCAGACAA CCCAGCCGCT
13381 TACGCCTGGC CAACCGCCCG TTCCTCCACA CATGGGGCAT TCCACGGCGT CGGTGCCTGG
13441 TTGTTCTTGA TTTTCCATGC CGCCTCCTTT AGCCGCTAAA ATTCATCTAC TCATTTATTC
13501 ATTTGCTCAT TTACTCTGGT AGCTGCGCGA TGTATTCAGA TAGCAGCTCG GTAATGGTCT
13561 TGCCTTGGCG TACCGCGTAC ATCTTCAGCT TGGTGTGATC CTCCGCCGGC AACTGAAAGT
13621 TGACCCGCTT CATGGCTGGC GTGTCTGCCA GGCTGGCCAA CGTTGCAGCC TTGCTGCTGC
13681 GTGCGCTCGG ACGGCCGGCA CTTAGCGTGT TTGTGCTTTT GCTCATTTTC TCTTTACCTC
13741 ATTAACTCAA ATGAGTTTTG ATTTAATTTC AGCGGCCAGC GCCTGGACCT CGCGGGCAGC
13801 GTCGCCCTCG GGTTCTGATT CAAGAACGGT TGTGCCGGCG GCGGCAGTGC CTGGGTAGCT
13861 CACGCGCTGC GTGATACGGG ACTCAAGAAT GGGCAGCTCG TACCCGGCCA GCGCCTCGGC
13921 AACCTCACCG CCGATGCGCG TGCCTTTGAT CGCCCGCGAC ACGACAAAGG CCGCTTGTAG
13981 CCTTCCATCC GTGACCTCAA TGCGCTGCTT AACCAGCTCC ACCAGGTCGG CGGTGGCCCA
14041 TATGTCGTAA GGGCTTGGCT GCACCGGAAT CAGCACGAAG TCGGCTGCCT TGATCGCGGA
14101 CACAGCCAAG TCCGCCGCCT GGGGCGCTCC GTCGATCACT ACGAAGTCGC GCCGGCCGAT
14161 GGCCTTCACG TCGCGGTCAA TCGTCGGGCG GTCGATGCCG ACAACGGTTA GCGGTTGATC
14221 TTCCCGCACG GCCGCCCAAT CGCGGGCACT GCCCTGGGGA TCGGAATCGA CTAACAGAAC
14281 ATCGGCCCCG GCGAGTTGCA GGGCGCGGGC TAGATGGGTT GCGATGGTCG TCTTGCCTGA
14341 CCCGCCTTTC TGGTTAAGTA CAGCGATAAC CTTCATGCGT TCCCCTTGCG TATTTGTTTA
14401 TTTACTCATC GCATCATATA CGCAGCGACC GCATGACGCA AGCTGTTTTA CTCAAATACA
14461 CATCACCTTT TTAGACGGCG GCGCTCGGTT TCTTCAGCGG CCAAGCTGGC CGGCCAGGCC
14521 GCCAGCTTGG CATCAGACAA ACCGGCCAGG ATTTCATGCA GCCGCACGGT TGAGACGTGC
14581 GCGGGCGGCT CGAACACGTA CCCGGCCGCG ATCATCTCCG CCTCGATCTC TTCGGTAATG
14641 AAAAACGGTT CGTCCTGGCC GTCCTGGTGC GGTTTCATGC TTGTTCCTCT TGGCGTTCAT
14701 TCTCGGCGGC CGCCAGGGCG TCGGCCTCGG TCAATGCGTC CTCACGGAAG GCACCGCGCC
14761 GCCTGGCCTC GGTGGGCGTC ACTTCCTCGC TGCGCTCAAG TGCGCGGTAC AGGGTCGAGC
14821 GATGCACGCC AAGCAGTGCA GCCGCCTCTT TCACGGTGCG GCCTTCCTGG TCGATCAGCT
14881 CGCGGGCGTG CGCGATCTGT GCCGGGGTGA GGGTAGGGCG GGGGCCAAAC TTCACGCCTC
14941 GGGCCTTGGC GGCCTCGCGC CCGCTCCGGG TGCGGTCGAT GATTAGGGAA CGCTCGAACT
15001 CGGCAATGCC GGCGAACACG GTCAACACCA TGCGGCCGGC CGGCGTGGTG GTGTCGGCCC
15061 ACGGCTCTGC CAGGCTACGC AGGCCCGCGC CGGCCTCCTG GATGCGCTCG GCAATGTCCA
15121 GTAGGTCGCG GGTGCTGCGG GCCAGGCGGT CTAGCCTGGT CACTGTCACA ACGTCGCCAG
15181 GGCGTAGGTG GTCAAGCATC CTGGCCAGCT CCGGGCGGTC GCGCCTGGTG CCGGTGATCT
15241 TCTCGGAAAA CAGCTTGGTG CAGCCGGCCG CGTGCAGTTC GGCCCGTTGG TTGGTCAAGT
15301 CCTGGTCGTC GGTGCTGACG CGGGCATAGC CCAGCAGGCC AGCGGCGGCG CTCTTGTTCA
15361 TGGCGTAATG TCTCCGGTTC TAGTCGCAAG TATTCTACTT TATGCGACTA AAACACGCGA
15421 CAAGAAAACG CCAGGAAAAG GGCAGGGCGG CAGCCTGTCG CGTAACTTAG GACTTGTGCG
15481 ACATGTCGTT TTCAGAAGAC GGCTGCACTG AACGTCAGAA GCCGACTGCA CTATAGCAGC
15541 GGAGGGGTTG GATCAAAGTA CTTTGATCCC GAGGGGAACC CTGTGGTTGG CATGCACATA
15601 CAAATGGACG AACGGATAAA CCTTTTCACG CCCTTTTAAA TATCCGTTAT TCTAATAAAC
15661 GCTCTTTTCT CTTAG

//
